# Exploring Neural Mechanisms of Language Switching: An fMRI Study Using a Functional Localizer Approach

**DOI:** 10.64898/2026.03.02.708926

**Authors:** Keng-Yu Lin, Agata Wolna, Jakub Szewczyk, Kalinka Timmer, Michele Diaz, Zofia Wodniecka

## Abstract

When bilinguals frequently switch between their first (L1) and second (L2) languages during speech production, we usually observe two phenomena: (i) *language switch cost*, where switching to a different language is more difficult than staying in the same one, and (ii) *reversed language dominance*, where L1 production becomes slower than L2 production. These effects are thought to reflect language control mechanisms, yet the underlying neural bases remain debated.

In this study, we addressed this question by using the precision functional magnetic resonance imaging (fMRI) based on functional localization. Forty-one Polish–English bilinguals performed a language switching task (LST), in which they named pictures in L1 or L2 based on color cues. We investigated mechanisms behind two indices of language control commonly observed in the LST. First, we asked whether the domain-general resources supporting *language switch cost* overlap with *nonverbal task switch cost*. Second, we asked whether *reversed language dominance* reflects changes in language activation in the language-specific system, or whether it is related to increased engagement of domain-general control mechanisms. Results indicated that the *language switch cost* and *nonverbal task switch cost* share overlapping domain-general neural mechanisms. Similar to the *language switch cost*, *reversed language dominance* primarily engages domain-general processes rather than language-specific resources.

**Highlights:**

- fMRI combined with functional localization approach is implemented to examine the neural mechanisms underlying *language switch cost* and *reversed language dominance*.
- *Language switch cost* relies on neural mechanisms shared with *nonverbal switch cost* within the Multiple Demand network.
- *Reversed language dominance* is primarily supported by the domain-general rather than the language-specific mechanisms.
- Domain-general neural mechanisms play a pivotal role in bilingual language switching in speech production.

## 1. Introduction

Bilinguals regularly use both their native language (L1) and a second language (L2) in daily communication, and both languages are omnipresent in their everyday experience. Consequently, both languages remain co-activated even when only one is being used (e.g., Chen et al., 2017; Meade, Midgley, Dijkstra, et al., 2018; Meade, Midgley, & Holcomb, 2018; Thierry & Wu, 2007). To produce the intended language in a given context, bilingual speakers rely on control mechanisms to manage this concurrent co-activation (e.g., Reverberi et al., 2015). This need for control is especially pronounced in challenging situations, particularly those involving frequent switching between languages. In laboratory settings, difficulties related to language switching have been observed as increased error rates, naming latencies, and slower speech rates (Bergmann et al., 2015; Bradlow et al., 2017; Broos et al., 2021; Gollan & Ferreira, 2009). On the neural level, frequently switching between languages has been linked to increased engagement of neural resources (Abutalebi et al., 2008; Abutalebi & Green, 2007, 2016; De Baene et al., 2015; de Bruin et al., 2014; Gollan & Ferreira, 2009; Green & Abutalebi, 2013; Meuter & Allport, 1999). It has been proposed that it reflects increased demands imposed over the bilingual language control (BLC) mechanisms, a set of mechanisms which allows bilinguals to efficiently manage the co-activation of the two languages (Abutalebi et al., 2008; Abutalebi & Green, 2007, 2008, 2016; Green & Abutalebi, 2013). However, despite substantial empirical evidence focused on understanding neurocognitive mechanisms of bilingual language control, it is still unclear (i) to what extent within the domain-general executive control system, mechanisms underlying language switching overlap with mechanisms supporting non-linguistic switching, and (ii) whether the differences in the language dominance, observed in situations requiring frequent switching, reflect mainly the engagement of the (domain-general) language control mechanisms or a change in how the two languages are processed by the language-specific system in the brain.

To address this gap, in this study, we used functional magnetic resonance imaging (fMRI) to investigate the neural mechanisms underlying bilingual language control in a language switching task (LST). Unlike previous studies, which employed group-averaging approaches, we used precision fMRI: an approach that identifies functional areas of interest in the brain on the individual subject level, allowing us to precisely assess the functional and spatial overlap between functional systems in the brain. In the following sections, we discuss the theoretical motivation for our experiment and highlight how the precision fMRI approach can help address the unresolved questions in research exploring neurocognitive basis of the bilingual language control.

### 1.1. Language switching paradigm and bilingual language control mechanisms

Bilingual language control has been most frequently investigated using a language switching task (Declerck & Koch, 2023; Declerck & Philipp, 2015). In this task, participants are asked to name pictures or numbers in either L1 or L2, and the language of response changes on a trial-by-trial basis, depending on the cue they receive. Two phenomena, exemplifying potentially different mechanisms of bilingual language control, have been observed in the LST: (1) *language switch cost* and (2) *reversed language dominance* (Goldrick & Gollan, 2023; Timmer et al., 2019, 2025).

The *language switch cost* refers to the additional behavioral cost associated with switching from one language to another. This cost is reflected in both slower naming latencies and increased error rates (e.g., Broos et al., 2021). Importantly, the switch cost is not limited to the language domain and is also reported in nonverbal task switching (TS) paradigms, where participants alternate between different nonverbal tasks instead of repeating the same one (Prior & Gollan, 2011; Weissberger et al., 2012, 2015). The *reversed language dominance*, on the other hand, refers to a paradoxical phenomenon in which speech production in the dominant language (typically L1) becomes slower relative to the less dominant language (Declerck & Philipp, 2015; Goldrick & Gollan, 2023). Unlike relatively robust evidence for *language switch cost* in bilingualism research, findings on the *reversed language dominance* are less consistent, and its robustness remains debated (see Declerck et al., 2020 for review; see Gade et al., 2021, and Goldrick & Gollan, 2023 for contrasting evidence).

The existing neurocognitive model of BLC provides very broad delineation of the brain basis of the mechanisms that support bilingual speech production and allow one to efficiently switch between languages. In particular, the BLC model proposes the engagement of a network of mechanisms, encompassing brain regions of the bilateral inferior frontal and parietal cortices, anterior cingulate cortex (ACC), pre-supplementary motor area (pre-SMA), basal ganglia, and thalamus (Abutalebi & Green, 2016; J. Wu et al., 2019; Y. J. Wu et al., 2021). However, it is unclear to what extent the mechanisms of bilingual language control correspond to specific adjustments of the neurocognitive system that selectively support switching between languages, and to what extent this ability relies on existing mechanisms of cognitive control which support a variety of non-linguistic tasks. Also, it is unclear how demanding situations, such as frequent switching between languages, affects domain-general and language-specific processing.

### 1.2. Neural evidence of bilingual language production and nonverbal task processing

A substantial body of research suggests that bilingual language production interacts with nonverbal task processing, implying that the mechanisms supporting bilingual language control partially overlap with domain-general executive functions. For example, Kałamała et al., (2022) demonstrated that L2 use modulates the P3 and N450 ERP components elicited by nonverbal inhibition tasks. Timmer et al., (2017) reported that bilinguals show greater overlap in scalp distributions between language switching and nonverbal task switching than monolinguals. Together, these EEG findings indicate that language switching and task switching rely on at least partly shared neural mechanisms.

Neuroimaging studies using group-averaging fMRI also support this view of shared neural mechanisms. Branzi, Della Rosa, et al., (2016) found that the left prefrontal cortex supports both language switching and nonverbal task switching. Similarly, De Baene et al., (2015) reported overlapping brain activation during both language switching and nonverbal task switching. Although this study also identified some task-specific brain activation, the authors claimed that the discrepancy could have resulted from differences in the tasks per se (e.g., response modality: speech production in the language switching task vs. button pressing in the nonverbal task switching task). A limitation of group-averaging approaches, however, is that it may not reveal neighboring but functionally distinct regions (see Fedorenko, 2021 for a discussion).

More recent evidence looking at the engagement of domain-general and language-specific resources in bilingual speech production comes from studies using precision fMRI methods based on functional localization of brain networks. This approach identifies functional systems in individual brains, without relying on between-subject voxel-wise averaging of functional signals, which allows to differentiate the relevant areas from nearby, functionally distinct regions and precisely quantify their individual contributions to the task at hand (Fedorenko et al., 2010; Kanwisher, 2010). For example, using this approach, Wolna et al. (2024a, 2024b) found that the domain-general fronto-parietal system in the brain (the Multiple Demand (MD) network) is sensitive to the increased difficulty of L2 speech production and the so-called L2 after-effect, a lingering difficulty in accessing L1 following L2 use. While L1 and L2 speech production engages the language system, the use of L2 appears to additionally draw on domain-general resources in speech production (Wolna et al., 2024a, 2024b) and comprehension (Malik-Moraleda et al., 2024). In language switching contexts, the opposite pattern is sometimes observed, where processing of the native language (L1) becomes temporarily more costly than processing of L2 (i.e., *reversed language dominance*). This raises a critical and intriguing question regarding such *reversed language dominance* observed in the LST: if frequent switching indeed affects language dominance, is this change reflected in the language system? Or does the effect which we refer to as *reversed language dominance* reflect, in fact, sustained, domain-general control mechanisms, and not directly affect the language-specific mechanisms underlying L1 and L2 production?

### 1.3. The present study

In the current study, we used a precision fMRI approach to investigate the extent to which *switch costs* in language switching and nonverbal task switching rely on shared neural mechanisms, as well as the neural basis of *reversed language dominance*. This approach offers key advantages over conventional group-averaging whole-brain analysis: it identifies functional networks of interest using “localizer” tasks, which are designed to robustly target specific cognitive processes and reliably identify the corresponding brain regions at the individual subject level (Fedorenko et al., 2010). This approach circumvents several issues which affects traditional group-level averaging approaches: first, it identifies brain regions based on its function, irrespective of the anatomical regions where activations are observed (the "reversed inference" problem, Poldrack, 2006). Second, it allows identification of brain networks at the individual subject level, which might otherwise be conflated in traditional group-level averaging approaches (Nieto-Castañón & Fedorenko, 2012; Wolna et al., 2025) given the inter-individual variability in functional organization of the neocortex (Braga & Buckner, 2017; DiNicola et al., 2023; Frost & Goebel, 2012; Tahmasebi et al., 2012). Here, we adapted this approach and used functional localizer tasks designed to identify distinct neural networks associated with language, domain-general, and switching mechanisms. We evaluated how these systems respond to a language switching task to address two research questions central to long-lasting debates in the neuroscience of bilingual language processing: (1) Are neural mechanisms supporting *language switch cost* fundamentally distinct from those supporting the switch cost in nonverbal task switching, or do they reflect generally overlapping neural mechanisms within the same network? (2) Does the *reversed language dominance* reflect increased engagement of language-specific resources, domain-general resources, or both? If the neural mechanisms underlying *language switch cost* overlap with the switch cost involved in nonverbal task switching, we should observe similar activation patterns within the same neural network for both types of switching tasks, with some degree of spatial correlation and overlap between the activation maps. If *reversed language dominance* relies on domain-general control, it should be reflected in an increased engagement of computations supported by the domain-general network. If the effect also engages language-specific mechanisms, which might be driven by the current activation of a given language, we should expect to see an increased engagement in the language network as well.

To our knowledge, this study is the first to use functional localizer analysis to investigate speech production in bilingual language switching. This approach enhances our ability to uncover the neural mechanisms underlying bilingual language control and provides valuable insights into the interplay between language-specific and domain-general cognitive processes.

## 2. Methods

### 2.1. Participants

Forty-five neurologically healthy Polish–English bilinguals participated in the study. Data from 4 participants were excluded due to technical issues during scanning (n=1), and artifacts of excessive motions (n=3). This left a final sample of 41 participants (35 females, 6 males, age=18-32 years of age, mean age=22.3 years, SD=2.93). All self-reported to be proficient in English (B2 or higher) and obtained at least 19/25 points on General English Test by Cambridge Assessment (https://www.cambridgeenglish.org/test-your-english/general-english/) (mean score=22.3, SD=1.77). Their proficiency in English was also assessed with LexTale (mean accuracy=0.78, SD=0.1) (Lemhöfer & Broersma, 2012). They also completed a language background questionnaire detailing their experience and use of all known languages (up to 4 languages), as shown in Table 1 below. All participants were right-handed (mean laterality quotient=69.8, range=0-100).

**Table 1.** Language experience of the participants. The table presents the averaged scores from the language background questionnaire including self-rated proficiency (rated for five language abilities on a scale from 1 to 10), age of acquisition (onset of learning and age at which participants could use a given language in a simple communication or comprehend a text or other materials), and daily language use representing % of time during the day they use a given language. Additionally, the mean number of words produced in category and letter verbal fluency tasks is reported for L1 and L2.

| Language | L1<br>(Polish) | L2<br>(English) | L3<br>(others) | L4<br>(others) |
| --- | --- | --- | --- | --- |
| N | 41 | 41 | 38 | 15 |
| **Self-rated proficiency (1-10)** |  |  |  |  |
| Accent | 9.73 | 6.59 | 3.79 | 3.93 |
| Listening | 9.93 | 7.88 | 4.21 | 3.40 |
| Reading | 9.90 | 8.32 | 4.63 | 4.20 |
| Speaking | 9.90 | 7.34 | 3.50 | 3.27 |
| Writing | 9.80 | 7.41 | 3.26 | 3.20 |
| **Age of acquisition** |  |  |  |  |
| Onset of learning | 0.00 | 5.41 | 14.84 | 15.00 |
| Being able to use it | 1.90 | 9.37 | 15.97 | 16.07 |
| **Daily use of language** |  |  |  |  |
| Active | 83.71% | 14.83% | 1.34% | 0.60% |
| Passive | 59.07% | 34.51% | 3.42% | 2.20% |
| **Verbal fluency**<br>(averaged number of words produced) |  |  |  |  |
| Category | 20.36 | 14.44 |  |  |
| Letter | 12.08 | 10.56 |  |  |

**Note:** Participants were Polish–English bilinguals, and thus L1 always corresponds to Polish and L2 to English. If they declared to know more languages, up to 4 languages were tracked for the language background questionnaire. The other languages varied across participants, including Czech, French, German, Hindi, Italian, Japanese, Latin, Portuguese, Russian, Spanish, Swedish, Turkish, Norwegian.

All participants gave informed consent prior to their participation in the experiments and received monetary remuneration for their participation. The study received approval from the Ethics Committee of the Institute of Psychology of Jagiellonian University concerning experimental studies with human participants.

### 2.2. Task design

Our study comprised three experimental sessions (see Figure 1 for an overview): one behavioral and two MRI sessions. In the behavioral session, participants completed verbal fluency tasks in L1 and L2, a LexTale task to assess their English proficiency (Lemhöfer & Broersma, 2012), a language questionnaire, and a short version of the picture naming task (using the same design and materials as in the MRI session). In one MRI session, participants completed a picture naming task, including single language runs and language switching runs, and in the other MRI session, they completed tasks which were used as functional localizers, including the auditory language localizer (Malik-Moraleda et al., 2022), Multiple Demand network localizer (Fedorenko et al., 2013), and the nonverbal task switching localizer.

**Figure 1.**
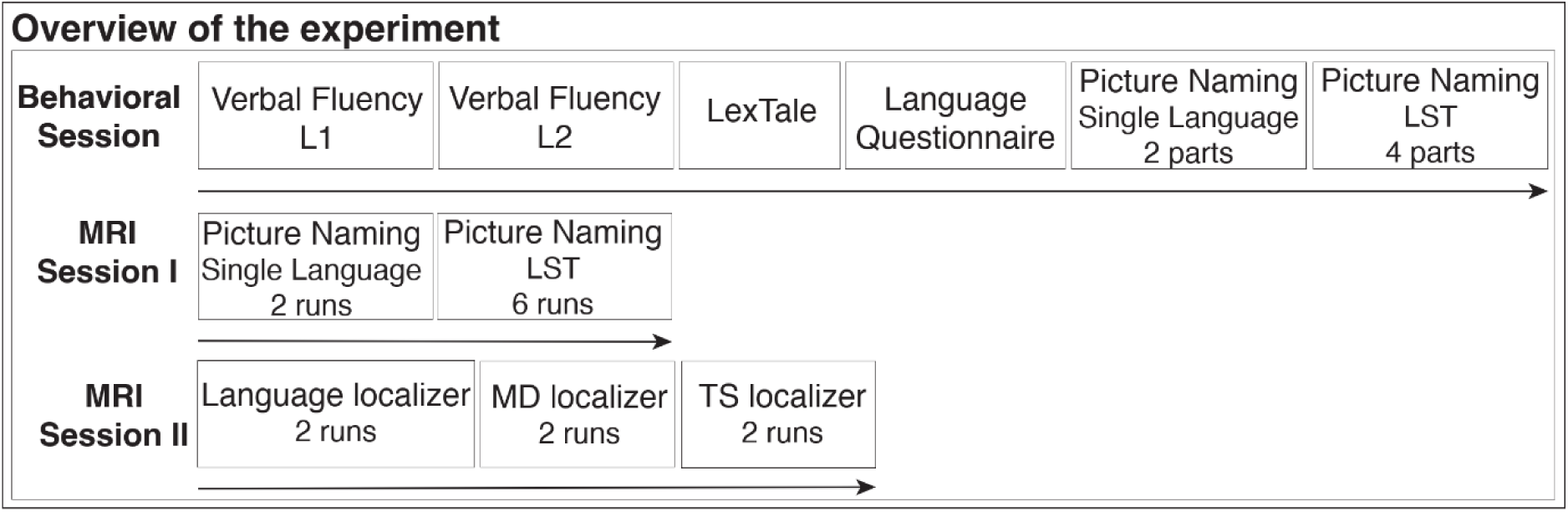
Overview of the design. The figure presents the task flow for the three sessions completed by each participant. The behavioral session and two MRI sessions were held over three different days. In the behavioral session, participants completed the tasks in a fixed order. In the first MRI session, the language order of the two single language runs, completed at the beginning, was counterbalanced, and followed by six language switching runs. In the second MRI session, participants completed the localizer tasks in a fixed order.

### 2.3. Behavioral session

#### 2.3.1. Verbal fluency task

In the verbal fluency task, participants were asked to produce as many words as possible belonging to a given category (fruit or animal) or words starting with a designated letter (B or M) within a 1-minute time limit. They completed the task in L1 first, followed by L2 (one category and one letter per language, with cues and their order counterbalanced between participants). The averaged words produced were shown in Table 1.

#### 2.3.2. LexTale task

The LexTale task served as a proxy of participants’ proficiency in English, which could supplement the subjective self-ratings. In each trial of the LexTale task, participants were presented with a string of letters and were instructed to indicate whether it was an existing English word via button pressing. We utilized the same experimental materials (40 words, 20 non-words) as in the original LexTale materials (Lemhöfer & Broersma, 2012).

#### 2.3.3. Language questionnaire

Participants were asked to fill out a questionnaire including detailed questions about their language experience. The language experience questionnaire required participants to provide information on their L1 (Polish) and L2 (English), as well as up to two additional languages if they knew any. For each language, they self-rated their proficiency in five language skills: reading, writing, listening, speaking, and accent. They also provided information on the age of acquisition (onset of exposure and onset of being able to use a given language in a simple conversation or to understand a simple text or video) as well as details of their language use (i.e., how much time during a typical day they passively and actively use each language) and language experience under different contexts, such as experience abroad, at home, and at work.

#### 2.3.4. Picture naming task

In the behavioral session, participants performed the picture naming task in which they name pictures in L1 or L2 based on the colored frame around it. The same materials and design were used in the behavioral and MRI sessions, with the behavioral session containing fewer parts (6 parts) than in the MRI session (8 parts, corresponding to 8 runs). For details of the picture naming task, see the task description in the MRI session (below).

### 2.4. MRI session

#### 2.4.1. Picture naming task

Participants were asked to name pictures in L1 or L2 based on a color-coded frame around the image. The color-language mapping was counterbalanced between participants. This design included two types of trials: *Switch* (named in a different language than the previous trial) and *Repeat* (named in the same language as previous trial), named in two languages (L1 and L2). In the single language runs, pictures were named consistently in one language. In the language switching task runs, each trial type was repeated the same number of times in each language within each run (12 per condition). Each run used the same set of 48 pictures. Note that in the behavioral session, participants completed 6 parts of the task using the same materials as the main task in the MRI session. They were therefore familiarized with the pictures and their names before getting into the scanner. Participants’ responses were recorded in the scanner using a FOMRI III, an MR-compatible, fiber-optic microphone system (Optoacoustics, Or-Yehuda, Israel).

Participants completed a total of 8 runs of the picture naming task. The first two runs consisted of blocks in which participants named pictures exclusively in either L1 or L2 within a given run (i.e., the same color frame was used for all 48 trials within a run). The language order was counterbalanced across participants, with some naming pictures in L1 during the first run and L2 during the second, while others completed the runs in the reversed order. The remaining six runs featured mixed language blocks and engaged language switching, where participants switched between L1 and L2 depending on the cue provided by the colored frame. The trials within each run were pseudo-randomized, so that the same language would be repeated 1-4 times in a row. The present study primarily focused on the six LST runs to investigate language switching.

Each run started with a 10s-long fixation period, followed by 48 picture naming trials. Each picture was displayed on the screen for 2.5s, enclosed by the color frame indicating the language to name it, which appeared on the screen simultaneously with the picture onset and remained there for the entire duration of the trial. After that, a fixation cross appeared with a jittered duration ranging from 0.75s to 13.4s (mean jittered duration=4.27s). The total duration of each run was 335 seconds. In the analysis looking at the overlap of language-specific and non-linguistic switching mechanism, the *Switch*>*Repeat* contrast was used as a localizer to target brain areas related to bilingual language switching. See Figure 2 for a schematic procedure for the task.

**Figure 2.**
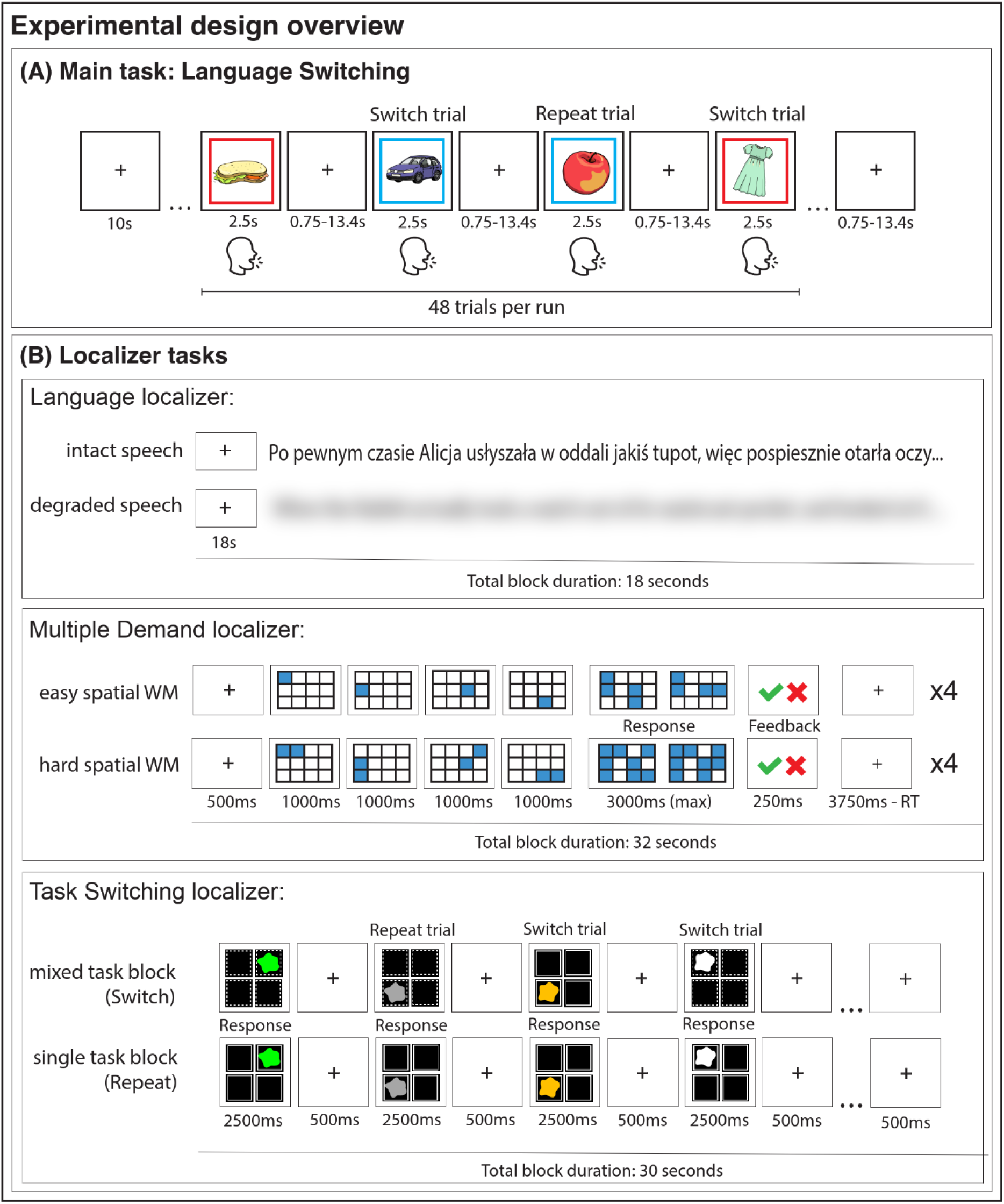
Overview of the experimental tasks. Panel (A) presents the main picture naming task where pictures are to be named in L1 or L2 according to the colored frame around the picture. The color-to-language mapping was counterbalanced between participants. The task includes two types of trials: the *Repeat* trials, when the language of naming in a trial is the same as in the previous trial, and the *Switch* trials, when the language of naming changes relative to the previous trial. Each run of this task includes 48 trials (12 per trial type and language combination). Panel (B) presents the schematic overview of the localizer tasks. In the language localizer, participants heard either *Intact* passages from Alice in Wonderland in L1, or a *Degraded* version of the audio. The Multiple Demand localizer was based on a spatial working memory task, where a 3x4 grid was presented, and a sequence of squares flashed up within the grid (one at a time in the easy condition, or two at a time in the hard condition). Participants were asked to keep track of the square locations (4 or 8 in total in *Easy* and *Hard* conditions, respectively) and, at the end of the trial, choose a grid with the correct locations from two alternatives, followed by visual feedback. Trials in each condition were grouped into sets of 4 to make experimental blocks. In the Task Switching localizer, participants were asked to respond to a stimulus based on its color or location: in each trial, a dashed or solid line indicated which rule they should follow to respond. The line type-to-task mapping was counterbalanced between participants. Within each block, the rule either changed from trial to trial (the *Switch* condition) or remained the same throughout all 10 trials (the *Repeat* condition).

#### 2.4.2. Auditory language localizer task

Participants passively listened to auditory stimuli corresponding either to the snippets from Alice in Wonderland in L1 (the *Intact* condition) or acoustically degraded, incomprehensible speech passages (the *Degraded* condition). The acoustically degraded speech passages were created based on the intact passages of Alice in Wonderland (Malik-Moraleda et al., 2022). Each trial began with a 0.5s fixation cross, followed by a 17.5s auditory stimulus. The task included twelve auditory blocks and four 12s fixation blocks.

Participants completed two runs of the task, each lasting 264 seconds. The *Intact*>*Degraded* contrast was used as a localizer to target brain areas supporting high-level language processing; see Figure 2.

#### 2.4.3. Multiple demand localizer task

Participants performed a visual working memory task consisting of blocks of either easy or hard series of trials. Each trial began with a 0.5s fixation cross, followed by four 3x4 grids where different locations were marked, each presented sequentially for 1s. In the *Easy* condition, only one location was marked per grid (four in total); in the *Hard* condition, two locations were marked per grid (eight in total). Participants were required to memorize the locations marked in all four consecutive grids. After viewing the four grids, participants were prompted to choose the correct pattern from two alternatives during a 4s response frame. This frame consisted of up to 3.75s for the response, followed by 0.25s for the feedback. If participants responded before the 3.75s timeout, a fixation cross was displayed after the feedback for the remainder of the 4s frame; see Figure 2.

The task followed a blocked design. Each run included 10 blocks (five per condition), and each block contained 4 trials. Additionally, six 16s fixation blocks were inserted in each run. Participants completed two runs of the multiple demand localizer task, with each run lasting 436 seconds. The contrast *Hard*>*Easy* was used as a localizer to target brain areas supporting domain-general cognitive demands.

#### 2.4.4. Task switching localizer task

Participants performed a color-location judgment task. In each trial, an irregular pattern was displayed for 2.5s, followed by a 0.5s fixation cross. The pattern was either colored (green, yellow) or colorless (gray, white) and appeared randomly in one of four screen locations (upper left, upper right, lower left, lower right). A rectangular frame (solid or dashed) enclosed the pattern and served as a cue, directing participants to respond to the pattern based on either its color (colored vs. colorless) or its location (left vs. right); see Figure 2.

The task followed a blocked design. Participants completed two types of blocks: the *Repeat* blocks, in which the task remained the same throughout the block (three blocks requiring responses based on stimulus location and three based on stimulus color), and the *Switch* blocks, in which the response rule alternated between trials. Each run included twelve blocks (6 per condition), and each block contained 10 trials. Additionally, five 14s fixation blocks were included in each run. Participants completed two runs of the task, with each run lasting 430 seconds. The *Switch*>*Repeat* contrast was used as a localizer to target brain areas supporting the task switching mechanism.

### 2.5. MRI acquisition

Structural MRI data were acquired using a Siemens MAGNETOM Prisma 3T scanner (Siemens Healthineers, Erlangen, Germany) with a 64-channel head coil at the Centre for Brain Research at the Jagiellonian University in Krakow, Poland. High-resolution T1-weighted anatomical images were acquired using a 3D MPRAGE sequence (192 sagittal slices, TR=2300 ms, TE=2.32 ms, TI=900 ms, flip angle=8°, voxel size=0.9x0.94x0.94 mm). Functional T2*-weighted images were acquired using a 2D multiband EPI pulse sequences from the Center of Magnetic Resonance Research (CMRR), University of Minnesota (Xu et al., 2013) (TR=1400 ms, TE=27 ms, flip angle=70°, voxel size=3x3x3 mm, multiband acceleration factor=2, GRAPPA acceleration factor=2, phase encoding=A>>P). Fieldmap images were acquired using a dual-echo gradient echo sequence, matched spatially to the fMRI scans (TE1=4.92 ms, TE2=7.38 ms, TR=485 ms, flip angle=60°).

### 2.6. fMRI data processing

#### 2.6.1. Preprocessing

The functional and anatomical images were preprocessed using the fMRIPrep preprocessing pipeline (version 23.2.1) (Esteban et al., 2019; Markiewicz et al., 2025). In addition, MRIQC (version 24.1.0) (Esteban et al., 2017) was used to supplement the outputs from fMRIPrep to assess our data quality.

The T1-weighted (T1w) image was corrected for intensity non-uniformity using N4BiasFieldCorrection (Tustison et al., 2010) and skull-stripped with ANTs-based brain extraction. Brain tissue segmentation was applied and the image was segmented into cerebrospinal fluid (CSF), white-matter (WM) and gray-matter (GM), using the FAST tool (FSL, RRID:SCR_002823, Zhang et al., 2001). Volume-based spatial normalization was carried out and the image was normalized to the standard space of MNI152NLin2009cAsym through nonlinear registration with antsRegistration (ANTs 2.5.0).

The following steps were used to preprocess the functional images. First, a reference volume was generated, using a custom methodology of fMRIPrep, for head motion correction. Head-motion parameters with respect to the blood-oxygen-level-dependent (BOLD) reference (transformation matrices, and six corresponding rotation and translation parameters) were estimated before any spatiotemporal filtering using mcflirt (FSL, Jenkinson et al., 2002). Frames that exceeded a threshold of 1.5 mm framewise displacement (FD) or 5.0 standardized DVARS were annotated as motion outliers that were later used as censoring in the first level modeling. The estimated fieldmap was then aligned with rigid registration to the target echo planar imaging (EPI) reference run. Subsequently, fieldmap images were applied to correct distortions induced by B0 field inhomogeneity. The BOLD reference was then co-registered to the T1w reference using bbregister (FreeSurfer) which implements boundary-based registration with six degrees of freedom (Greve & Fischl, 2009). Following co-registration, functional images were normalized to MNI152NLin2009cAsym standard space via nonlinear registration with antsRegistration (ANTs 2.5.0). For each run, confound time-series were also calculated, including FD, DVARS, three region-wise global signals (whole-brain, white-matter, and CSF). In addition, a set of physiological regressors were extracted to allow for component-based noise correction (CompCor, Behzadi et al., 2007). Finally, smoothing was applied to the preprocessed data using SPM12 using a Gaussian kernel with an FWHM of 4 mm, to the fMRIPrep-processed data before statistical modeling.

#### 2.6.2. First level modeling

Prior to the first level analysis, the fMRIPrep-processed data was visually inspected for artifacts. Missing data and experimental runs that contained excessive motion artifacts were excluded. Visual inspection of the fMRIPrep and MRIQC outputs followed Provins et al., (2023)’s guidelines. Visible peaks and periodic fluctuations as well as prolonged deflections in the carpet plot were considered excessive motion. As a result of this quality check, 4 subjects were entirely excluded from further analyses (1 due to missing data caused by technical issues during scanning, and 3 due to excessive motions). Additionally, individual picture naming LST runs from 4 other subjects were excluded due to excessive motion (totaling 7 runs across these 4 subjects). Subsequently, responses in individual voxels were modeled using CONN toolbox, (https://web.conn-toolbox.org/) (Whitfield-Gabrieli & Nieto-Castanon, 2012). A General Linear Model (GLM) with predictors corresponding to experimental conditions in each task was fitted independently to each voxel’s BOLD time series. Experimental conditions were modeled with a boxcar function convolved with a double-gamma hemodynamic response function. Fixation was modeled implicitly. Temporal autocorrelations in the BOLD signal timeseries were accounted for by a combination of high-pass filtering with a 128 seconds cutoff, and whitening using an AR(1) model with autoregressive coefficient being 0.2 (first-order autoregressive model linearized around the coefficient a=0.2) to approximate the observed covariance of the functional data in the context of Restricted Maximum Likelihood estimation (ReML). In addition to experimental conditions, the GLM included the first-order temporal derivatives of each condition and the nuisance regressors including 6 standard motion parameters, and regressors corresponding to outlier volumes identified by fMRIPrep. Additionally, in the language switching task, an additional regressor corresponding to accuracy was included in the model to regress out incorrect trials from the signal.

#### 2.6.3. Group-level functional parcels and subject-specific functional ROIs

In the functional localization analysis, we defined individual subject functional regions of interest (fROIs) within group-level parcels, based on selected contrasts from the localizer tasks. We used previously validated group-level parcels corresponding to the language network and the domain-general MD network (available at https://www.evlab.mit.edu/resources) to ensure comparability with prior research. The language system included five left hemisphere parcels, and the MD system included twenty symmetrical parcels (10 per hemisphere).

Within each group-level parcel, we defined subject-specific fROIs by selecting the top 10% of voxels showing the strongest response to the relevant localizer task contrast: *Intact*>*Degraded* speech for the language localizer, *Hard*>*Easy* visual working memory for the MD localizer, *Switch*>*Repeat* for the nonverbal task switching and language switching tasks (for an overview of individual-level fROIs creation steps addressing both research questions, see Figure 3). Subsequently, (i) from each fROI, we extracted parameter estimates (beta estimates, which given the signal scaling in SPM are approximately reflecting the percent signal change) for all the conditions of interest and (ii) within the MD parcels, we evaluated the number of overlapping voxels and spatial correlations of activation patterns between language switching and nonverbal task switching tasks, defined based on the switching contrasts (i.e., *Repeat* and *Switch* in TS and LST). The fROI creation as well as the two analysis types described below were implemented with the spm_ss toolbox in Matlab (available at: https://www.evlab.mit.edu/resources).

**Figure 3.**
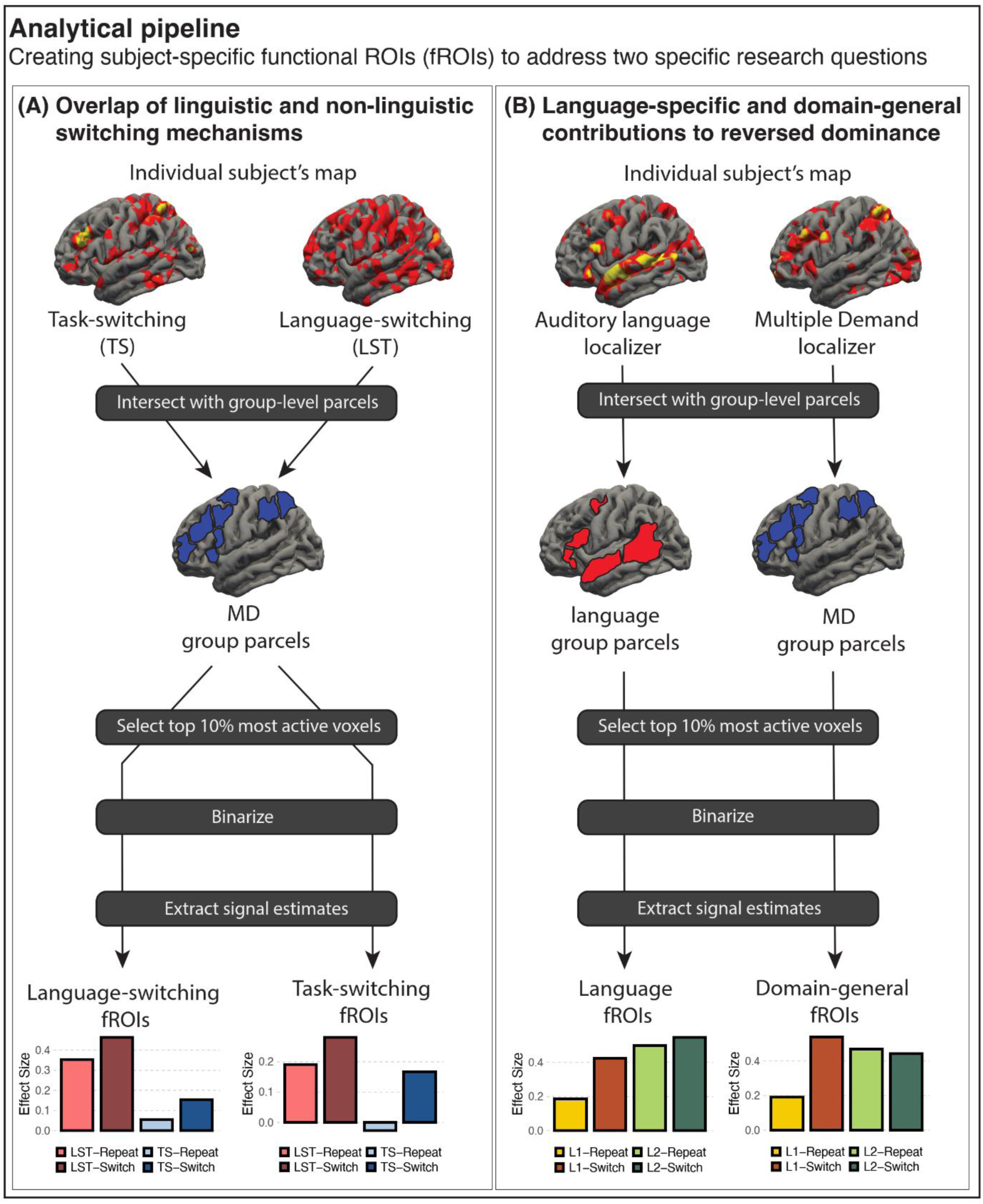
Overview of the analytical pipeline. Panel (A) presents the workflow used to address the question of overlap between language switching and nonverbal task switching mechanisms. Panel (B) presents the workflow used to address the question of language-specific and domain-general components supporting switching between languages. In both workflows, we intersected the statistical maps with group-level parcels: in workflow (A) we used the MD parcels to constrain both LST- and TS-fROIs and in workflow (B) we used language parcels to constrain the language fROIs and MD parcels to constrain domain-general fROIs. Subsequently, we selected the top 10% of the most active voxels within each parcel. We binarized the fROIs and extracted responses corresponding to the % BOLD signal change for the tasks of interest from each individual fROI.

#### 2.6.4. Estimation of responses to critical tasks

To investigate the origins of *switch costs*, we used the group-level MD network as a constraint to create subject-specific fROIs based on the language switching and nonverbal task switching tasks (Figure 3A). Subsequently, from both types of fROIs we extracted response estimates to the repeat and switch trials in both language switching task and nonverbal task switching task. To investigate the origins of *reversed language dominance*, we used group-level language and MD parcels as a constraint to create subject-specific fROIs based on the language and MD localizer tasks (Figure 3B). Subsequently, we extracted estimates of responses to L1 and L2 trials from both trial types (*Repeat*, *Switch*) in the LST. When estimating responses to tasks which were also used as localizers for a specific fROI, we relied on an across-runs cross-validation procedure, which avoids double-dipping by ensuring that different subsets of the data are used for functional ROI definition and response estimation (Kriegeskorte, 2011).

#### 2.6.5. Spatial evaluation of language-switching and nonverbal task-switching fROIs

In addition to examining the functional profiles of fROIs created based on language switching and nonverbal task switching contrasts (i.e., *Switch*>*Repeat*), we computed the voxel overlap for each pair of fROIs within each MD group-level parcel using Dice coefficient, calculated using the following formula:

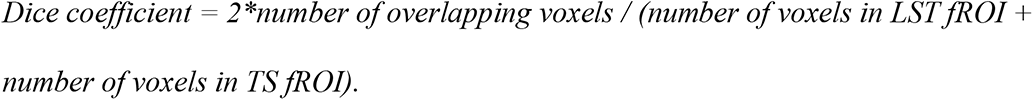

Furthermore, within each group-level MD parcel, we assessed the degree of voxel-wise spatial correlation between un-thresholded statistical maps corresponding to the language-switching and task-switching switching contrast (i.e., *Switch*>*Repeat* in LST and TS). For each subject, the correlation score was computed as the mean of voxel-to-voxel correlation (using the Fisher-transformed correlation coefficients) within each group-level MD parcel.

To estimate the ceiling for voxel overlap as well as the spatial correlations measures, we calculated these metrics within each task (i.e., separately for TS and LST), between-runs. To match the amount of data between the within-contrast and across-contrast comparisons, for the across-contrast comparisons, we obtained two correlation scores (between the first run of the first task and the second run of the second task, and between the second run of the first task and the first run of the second task) and averaged them to obtain a single across-contrast score. Finally, within-tasks estimates were averaged across the LST and TS tasks.

### 2.7. Statistical data analysis

Statistical analyses were conducted in R (R Core Team, 2025), using the lmerTest package (Kuznetsova et al., 2017). Pairwise comparisons relevant to our research questions were conducted using the emmeans package (Lenth, 2025) when statistically significant interactions of interest were observed. For all statistical analyses, we initially specified the maximal mixed effects model, following the recommendations outlined in Barr et al. (2013), and performed model reductions if the model could not converge. In cases of convergence issues, first, we removed correlations within the random structure. If necessary, we then progressively eliminated variables explaining the least variance in the random slopes until the model successfully converged. Mixed effects models were implemented using the default parameters and the optimizer bobyqa in the lmer() function. Statistical significance was determined using a threshold of p<.05.

#### 2.7.1. Behavioral data analysis

Responses to picture naming in the main language switching task were transcribed by a native Polish speaker and naming latencies were manually set to speech onset in each trial. Naming latency during the MRI session was analyzed for LST runs using linear mixed effects models.

Prior to statistical analysis, raw naming latencies were log-transformed to better meet the model’s assumption of normally distributed residuals. Additionally, incorrect responses and non-responses were excluded (4.92% of the data). The model included two categorical predictors, which were coded using the sum contrast: *Language* (L1=-0.5, L2=0.5) *and Trial*

*Type* (*Repeat*=-0.5, *Switch*=0.5). The data were analyzed using the following model:

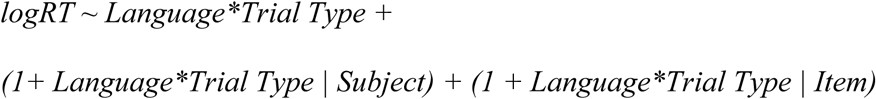

#### 2.7.2. Neuroimaging data analysis

For the fMRI data, we explored *switch costs* based on (1) the language switching fROIs within the MD network, and (2) the task switching fROIs within the MD network (Figure 3A). In addition, we also looked at the spatial overlap and correlation between LST and TS. To examine *reversed language dominance*, we focused on (1) the language fROIs within the language network, and (2) the MD fROIs within the MD network (Figure 3B).

##### 2.7.2.1. Is the switching mechanism shared between non-linguistic and linguistic switching within the MD system?

To explore whether language switching and nonverbal task switching share the same neural mechanisms, we directly compared brain activity between different trial types (*Repeat*, *Switch*) in both LST and TS within the MD network. Our dependent variable was the *Effect Size* (i.e., parameter estimates extracted from GLM, corresponding to % BOLD signal change), which was extracted based on LST-defined and TS-defined fROIs respectively, and was spatially constrained to the MD network parcels (Figure 3A). The model included two categorical predictors, which were coded using the sum contrast: *Trial Type* (*Repeat*=-0.5, *Switch*=0.5) and *Task* (*LST*=-0.5, *TS*=0.5). Subject and ROI were included as grouping factors in the random structure. The data were analyzed using the following model:

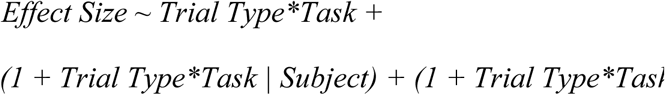

Additionally, we also examined the Trial Type×Task interaction effect within each group-level MD parcel using the following model, fitted to the responses in each ROI separately:

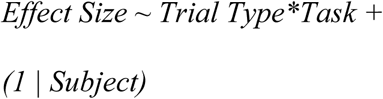

In addition to examining the functional profiles of the LST and TS, we compared their (i) spatial overlap and (ii) spatial correlations of the activation patterns. In the spatial overlap analysis, the dependent variable was the Dice coefficient and in the spatial correlation analysis, it was the Fisher-transformed correlation coefficients. Both analyses included one categorical predictor, *Task Pair*, corresponding to between-task comparisons (i.e., LST-TS) and the within-task comparisons (i.e., the average of within tasks comparisons for LST and TS). *Task Pair* was coded using a sum contrast (within-task average=-0.5, between-task comparison=0.5). The mixed effects model included Subject and ROI as random grouping factors. The data were analyzed using the following models:

To evaluate voxel overlap:

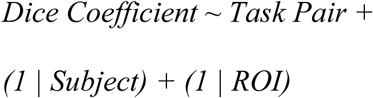

To evaluate spatial correlation:

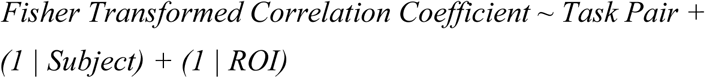

As with the functional profiles, we also investigated the *Task Pair* effect for spatial overlap and correlation within each parcel of the group-level MD network using the following models, fitted to the values in each ROI separately:

To evaluate voxel overlap:

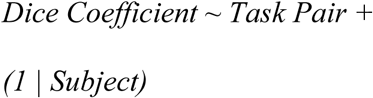

To evaluate spatial correlation:

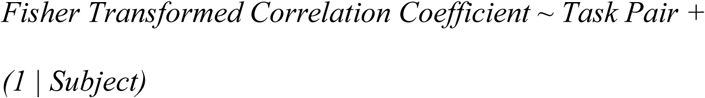

##### 2.7.2.2. Is the reversed language dominance driven by language-specific, domain-general control mechanisms, or both?

To examine the extent to which language-specific and domain-general mechanisms drive the *reversed language dominance*, we compared brain responses for L1 and L2 in the LST within the group-level language network and the MD network, respectively. Our dependent variable was the *Effect Size* (i.e., parameter estimates extracted from GLM, corresponding to % BOLD signal change), which was extracted based on fROIs in the language system and MD system (Figure 3B). The model included two categorical predictors, which were coded using the sum contrast: *Language* (L1=-0.5, L2=0.5), and *Trial Type* (*Repeat*=-0.5, *Switch*=0.5). Subject and ROI were included as grouping factors in the random structure. The data were analyzed using the following model:

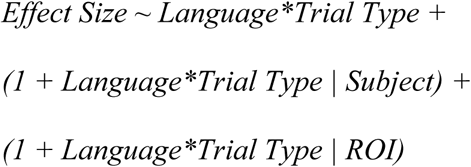

Furthermore, we also examined the Language effect within each individual parcel in both the language network and the MD network using the following model, fitted to the responses in each ROI separately:

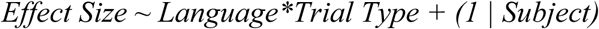

## 3. Results

### 3.1. Behavioral results

In the analysis of behavioral reaction times in the LST, we found a significant main effect of *Language*, with L1 exhibiting longer RTs than L2 (75ms, β=-0.07, SE=0.01, t-value=-7.41, p<.001), indicating the *reversed language dominance*. Additionally, a significant main effect of *Trial Type* was found, with the switch trials eliciting longer RTs than the repeat trials (62ms, β=0.059, SE=0.004, t-value=14.49, p<.001), indicating the *language switch cost*. No significant interaction was observed between *Language* and *Trial Type*.

### 3.2. fMRI results

#### 3.2.1. Is the switching mechanism shared between non-linguistic and linguistic switching within the MD system?

To address the first question, we examined the *switch costs* within the group-level MD parcels, using subject-specific fROIs defined by the *Switch*>*Repeat* contrast in both LST and TS. In the LST fROIs, a main effect of *Task* was found in which LST elicited increased brain activations than TS (β=-0.34, SE=0.10, t-value=-3.25, p<.010). Besides, a main effect of *Trial Type* was found, where switch trials elicited greater brain activation than repeat trials (β=0.09, SE=0.03, t-value=3.22, p<.010). No interaction effects were observed. See Table 2 and Figure 4A.

**Figure 4.**
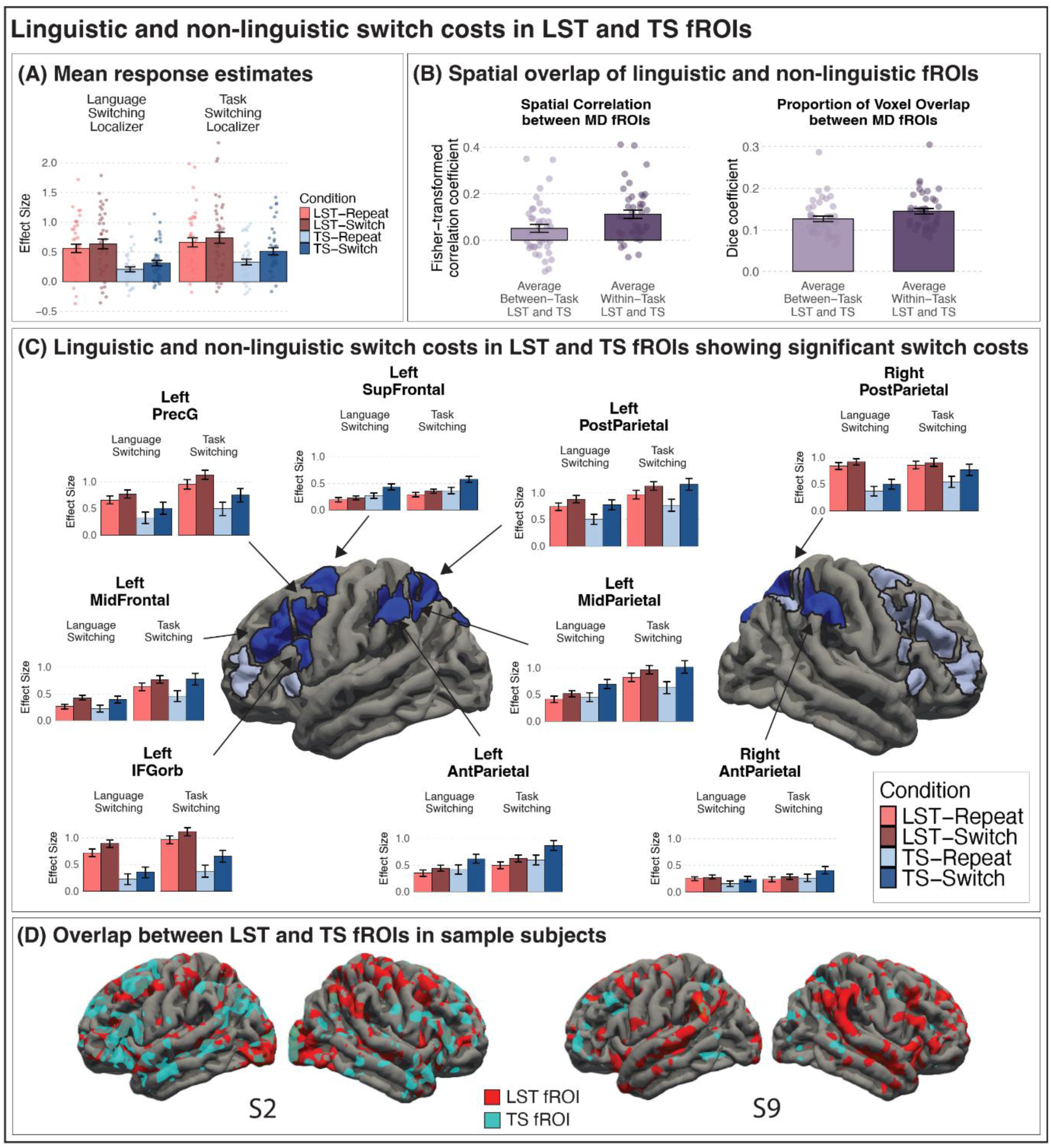
Summary of the analyses exploring the functional and spatial overlap between language-switching and task-switching switch costs. Panel (A) presents responses to the *language switch cost* (red bars) and *nonverbal switch cost* (blue bars) in the fROIs defined based on the LST (left) and TS (right). The bars correspond to effect sizes averaged over all fROIs in the MD system; individual points correspond to means by participant; error bars correspond to standard error of the mean. Panel (B) presents results of two analyses evaluating the spatial similarity based on the spatial correlations between activation patterns in each of the MD fROI (left) and voxel-level overlap between fROIs defined based on LST and TS (right). In both plots, the bars correspond to the within-task correlation or overlap between even and odd runs, averaged over LST and TS (dark violet bar) and to the average between-tasks correlation or overlap (light violet bar). Individual points correspond to means by participant; error bars correspond to the standard error of the mean. Panel (C) presents the profiles of responses to the language and nonverbal *switch costs* in MD parcels including fROIs that showed a significant *switch cost* in at least one of the switch tasks (LST or TS). These parcels are marked in dark blue, and the color coding of bars reflects the same information as in Panel (A). Panel (D) presents the surface projection of LST-based fROIs (red) and TS-based fROIs (blue) for two sample participants.

**Table 2.**
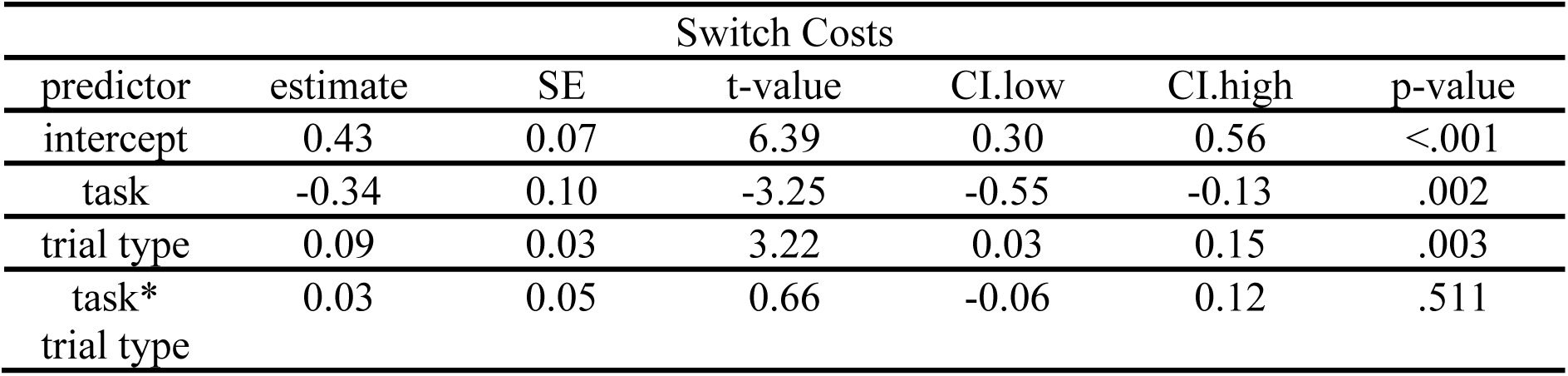
Results of switch costs in language switching and nonverbal task switching. The results present the *switch costs* in the LST fROIs within the MD network.

Separate analyses of each MD parcel focusing on the *Trial Type* effect revealed that 7 out of 20 parcels showed the main effect of *Trial Type* in which the switch trials elicited larger brain activations than the repeat trials, corresponding to the parcels falling into the posterior parietal, mid-parietal, anterior parietal, superior frontal, precentral gyrus, inferior frontal gyrus orbital part, and mid-frontal regions in the left hemisphere. No interaction effect between *Trial Type* (*Repeat*, *Switch*) and *Task* (LST, TS) was observed in the ROI-based analysis. See Figure 4C and supplementary materials for details of statistics.

In the TS fROIs, a main effect of *Task* was observed, with the LST eliciting greater brain activation than the TS (β=-0.28, SE=0.10, t-value=-2.81, p<.010). A main effect of *Trial Type* was also found, where the switch trials elicited larger brain activation than the repeat trials, consistent with the patterns observed in the LST fROIs (β=0.13, SE=0.03, t-value=3.80, p<.001). Furthermore, an interaction effect between *Task* and *Trial Type* was observed (β=0.10, SE=0.03, t-value=3.11, p<.010). Follow-up pairwise comparisons using the emmeans() package (Lenth, 2025) for the effect of interest (i.e., *Switch* vs. *Repeat* across tasks) showed that the switch trials elicited stronger brain activation than the repeat trials in both LST (β=-0.08, SE=0.04, z-ratio=-2.19, p<.050) and TS (β=-0.18, SE=0.04, z-ratio = - 4.50, p<.001), with the effect being larger in TS than in LST. See Table 3 and Figure 4A. Separate analyses of each individual MD parcel focusing on the *Trial Type* effect revealed that 9 out of 20 fROIs showed the main effect of *Trial Type* in which the switch trials had greater brain activations than the repeat trials. Similar to the results from the LST fROIs, these parcels included posterior parietal, mid-parietal, anterior parietal, superior frontal, precentral gyrus, inferior frontal gyrus orbital part, and mid-frontal regions in the left hemisphere. In addition, in the TS fROIs we observed the effect of *Trial Type* in the posterior and anterior parietal regions in the right hemisphere. No interaction effect between *Trial Type* and *Task* was observed in the ROI-based analysis. See Figure 4C and supplementary materials for details of statistics.

**Table 3.**
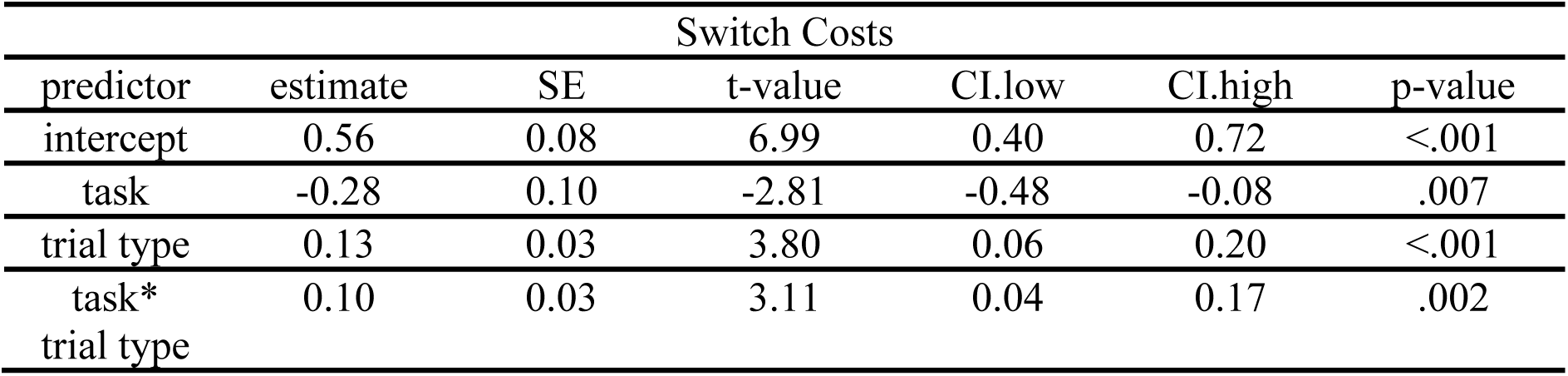
Results of switch costs in language switching and nonverbal task switching. The results present the *switch costs* in the TS fROIs within the MD network.

In addition to characterizing the functional profiles of the LST and TS fROIs within the MD system, we evaluated the spatial overlap between these two types of fROIs and spatial correlations between the activations elicited by language-switching and task-switching tasks. The results of both analyses showed similar patterns: the average between-task overlap and spatial correlation was above zero (Dice coefficient = 0.13; β=0.13, SE=0.01, t-value=11.86, p<.001; Fisher-transformed correlation coefficient = 0.05; β=0.05, SE=0.03, t-value=2.00, p=0.05). As expected, these values were smaller for the between-task comparison compared to the within-task comparison, with average between-task voxel overlap in between-task comparison corresponding to 87.5% of the average within-task voxel overlap (β=-0.02, SE=0.005, t-value=-3.92, p<.001) and average between-task correlation corresponding to 45.3% of the average with-task correlation (β=-0.06, SE=0.01, t-value=-5.76, p<.001). See Figure 4B.

#### 3.2.2. Is the reversed language dominance driven by language-specific, domain-general control mechanisms, or both?

To address this question, we evaluated the effect of *Language* in the language and domain-general MD systems. In the language network, we found a main effect of *Language* with L2 eliciting greater brain responses than L1 (β=0.22, SE=0.06, t-value=3.45, p<.050). In addition, a main effect of *Trial Type* was observed, with the switch trials eliciting greater responses compared to the repeat trials (β=0.14, SE=0.05, t-value=2.64, p<.010). See Table 4 and Figure 5A.

**Figure 5.**
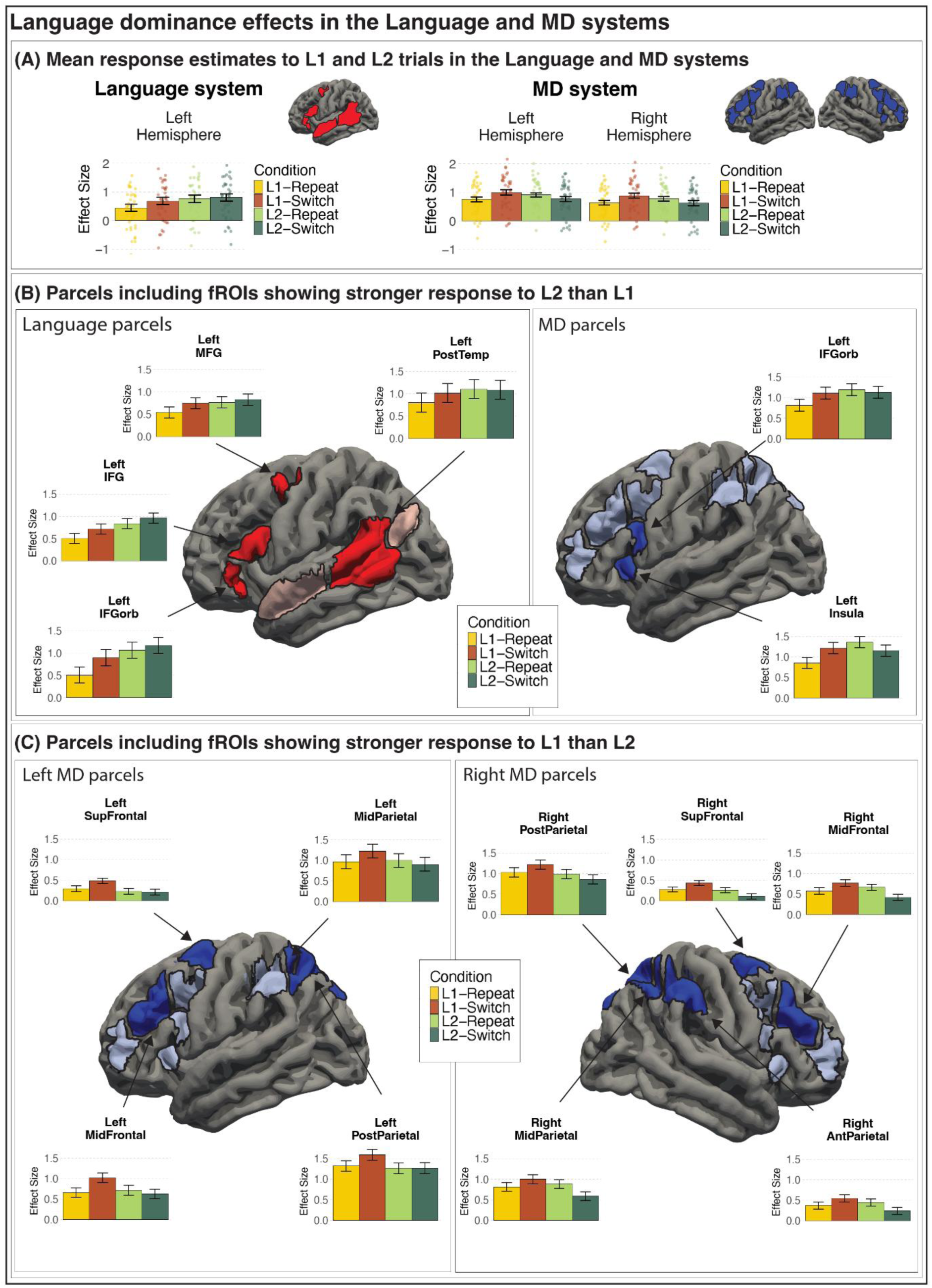
Summary of the analyses exploring the reversed language dominance. Panel (A) presents responses to languages across different trial types in the language (left) and Multiple Demand (right) systems. Panel (B) presents results of stronger responses to L2 than L1 in brain regions that reached statistical significance (dark red, dark blue). Panel (C) presents results of stronger responses to L1 than L2 (i.e., *reversed language dominance*) in brain regions that reached statistical significance (dark blue).

**Table 4.**
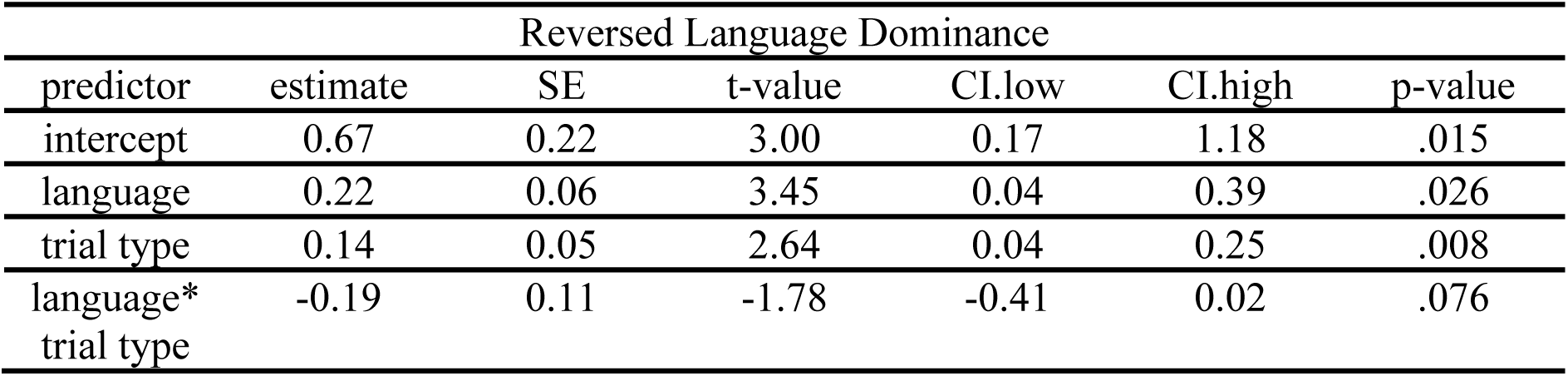
Results of reversed language dominance in the language network. The results present the *reversed language dominance* in the language fROIs within the language network.

Analysis of individual regions within the language network revealed a main effect of *Language* in which L2 elicited larger activations than L1 in inferior frontal gyrus orbital, inferior frontal gyrus, mid-frontal gyrus, and posterior temporal regions. Interaction effects between *Language* and *Trial Type* were also observed in the ROI-based analysis. In particular, the effect of L2 eliciting larger brain activations than L1 appeared exclusively in the repeat trials in the anterior and posterior temporal cortex. See Figure 5B and supplementary materials for details of statistics.

In the MD network, we found no main effect of *Language* or *Trial type*, but an interaction effect between *Language* and *Trial Type* (β=-0.38, SE=0.09, t-value=-4.25, p<.001). Follow-up pairwise comparisons revealed that L2 elicited greater brain activation than L1 in the repeat condition (β=-0.14, SE=0.07, z-ratio=-2.20, p<.050), while L1 elicited greater brain activation than L2 in the switch condition (β=0.24, SE=0.06, z-ratio=3.72, p<.001). See Table 5 and Figure 5A.

**Table 5.** Results of reversed language dominance in the Multiple Demand network. The results present the *reversed language dominance* in the MD fROIs within the MD network.

| Reversed Language Dominance |  |  |  |  |  |  |
| --- | --- | --- | --- | --- | --- | --- |
| predictor | estimate | SE | t-value | CI.low | CI.high | p-value |
| intercept | 0.79 | 0.11 | 7.35 | 0.58 | 1.01 | <.001 |
| language | -0.05 | 0.05 | -1.06 | -0.14 | 0.05 | .297 |
| trial type | 0.05 | 0.04 | 1.13 | -0.04 | 0.14 | .267 |
| language*<br>trial type | -0.38 | 0.09 | -4.25 | -0.57 | -0.20 | <.001 |

Analysis of individual regions within the MD network revealed a main effect of *Language* in which L1 elicited larger brain activations than L2 in 9 out of 20 regions, corresponding to posterior parietal, mid-parietal, superior frontal, mid-frontal regions at the left hemisphere, and posterior parietal, mid-parietal, anterior parietal, superior frontal, mid-frontal regions at the right hemisphere and L2 elicited larger brain activations than L1 in 2 out of 20 regions, corresponding to inferior frontal gyrus orbital part and insula at the left hemisphere. See Figure 5B, Figure 5C, and supplementary materials for details of statistics.

Interaction effects between *Language* and *Trial Type* were also observed in the ROI-based analysis. Similar to the language network, the effect of L2 eliciting larger brain activations than L1 appeared exclusively in the repeat trials in 9 out of 20 regions, corresponding to anterior parietal cortex, precentral gyrus, inferior frontal gyrus orbital part, insular cortex, medial frontal corte at the left hemisphere, and precentral gyrus, inferior frontal gyrus orbital part, insular cortex, medial frontal cortex at the right hemisphere. The effect of L1 eliciting larger brain activations than L2, on the other hand, appeared exclusively in the switch trials in most regions (15 out of 20), except for anterior parietal cortex, precentral gyrus, inferior frontal gyrus orbital part, insular cortex, medial frontal cortex at the left hemisphere. See supplementary materials for details of statistics.

## 4. Discussion

This fMRI study investigated the neural mechanisms of bilingual language control, focusing on two phenomena commonly observed in a language switching paradigm: the *language switch cost* and *reversed language dominance*. To evaluate the neural basis of *language switch cost*, we examined their functional and spatial overlap with the nonverbal switch cost within the domain-general MD network. To further understand the neural basis of *reversed language dominance*, we investigated how this effect is reflected in both the language network and the domain-general MD network. Our behavioral results replicated *language switch cost* and *reversed language dominance*. After confirming that we observed the critical behavioral effects, we proceeded to identify their underlying neural mechanisms. For the *switch costs*, we found evidence of overlapping neural mechanisms for both language switching and nonverbal task switching. For the *reversed language dominance*, we found stronger responses to L1 compared to L2 in several nodes of the domain-general MD network but not in the language network. We also observed an intriguing interaction in which L2 elicited greater brain responses than L1 during repeat trials, whereas L1 elicited greater responses than L2 during switch trials. Taken together, our findings highlight the critical involvement of the domain-general MD network in supporting language switching.

### 4.1. Switch costs across language and nonverbal domains rely on a shared neural mechanism

Consistent with previous neuroimaging studies (Branzi, Calabria, et al., 2016; De Baene et al., 2015), our results support the view that both language switching and nonverbal task switching recruit shared resources within the broader domain-general neural network.

First, we were able to identify brain activations related to nonverbal task switching and language switching in multiple nodes of the MD system, and second, they show very similar functional profiles. Moreover, we observed that the brain regions supporting task switching and language switching showed spatial overlap within the MD system. Our results are in line with recent claims suggesting that the MD system shows highly similar neural patterns across different tasks that engage domain-general cognitive control processes (e.g., the n-back task, the rule switching task) (Assem et al., 2022, 2024; Shashidhara et al., 2024). In the case of bilingual language control, our results show that task-switching and language-switching rely on largely shared, domain-general mechanisms, which is in line with many previous findings. However, in this study for the first time we evaluate the overlap between these two tasks on individual-subject level, which provides a much higher level of precision than any of the previous empirical reports.

In our analyses of *switch costs*, we sought the effects for the language and nonverbal switching within the MD network, a system well-established as a domain-general hub for cognitive control. However, the BLC network (Abutalebi & Green, 2016; Green & Abutalebi, 2013) includes additional nodes that do not fully overlap with the MD system, including the subcortical nodes and the medial regions. To ensure that our inferences are not constrained by assuming that BLC is simply part of the MD network, we also examined *switch costs* in regions identified as part of the BLC network (Abutalebi & Green, 2016). The results show that the response patterns were similar to those obtained when looking at the MD system and no additional regions outside of this system were engaged in the linguistic switch costs (for details of the analytical approach and results, see the supplemental materials). We found no evidence that *switch costs* are reflected differently across the two networks, suggesting that the BLC network is not a functionally distinct system in the brain, but rather that the bilingual language control mechanisms rely on mechanisms supported by the broader, general-purpose MD system.

### 4.2. Reversed language dominance reflects a control process, not changes in language-specific representations

As discussed in the Introduction, *reversed language dominance*, which is a paradoxical slow-down of L1 observed in LST, is oftentimes considered to be a reflection of different mechanisms than *switch costs* (Declerck & Koch, 2023). Still, similar to the *switch costs*, we observed the *reversed language dominance* (i.e., stronger responses to L1 than to L2) in the domain-general MD network. This suggests that despite potential differences in the underlying cognitive mechanism, reversal of language dominance in LST relies as well on a domain-general, general-purpose neural network. Interestingly, in some of the MD nodes, including the precentral/inferior frontal gyrus (orbital part) and insula, we found the opposite effect: stronger activations in response to L2 than L1 (i.e., the typical language dominance effect for unbalanced bilinguals). This result suggests that the reversal of the language dominance does not reflect a modulation of the entire system that bilinguals engage in speech production, but instead, there seems to be an interesting dichotomy: while some elements of the neural network are responsible for reversal of dominance (presumably to facilitate the production under a demanding context), others focus on resolving the typically-observed difficulty related to speaking in a less proficient, non-dominant language (e.g., Wolna et al., 2024a). In contrast to the results within the domain-general MD system, within the language network, we did not find any trace of mechanisms supporting the *reversed language dominance*. However, in line with the sensitivity of this system to L2 vs. L1 contrast (Malik-Moraleda et al., 2024; Wolna et al., 2024a), we observed greater activation in response to L2 than L1. These findings suggest that the *reversed language dominance* in fact primarily reflects a control mechanism, a domain-general process within the repertoire of the postulated mechanisms of BLC. This mechanism regulates the co-activation of the two languages and re-configures the cognitive system of a bilingual to an optimal state, enabling it to support situations that require frequent switching between languages. This explanation is in line with the assumption of the Adaptive Control Hypothesis (Green & Abutalebi, 2013), which posits that depending on the surrounding language context, bilinguals tailor their control mechanisms to efficiently support a given conversational situation. This explanation is supported by data showing that the domain-general control network in bilinguals re-configures by strengthening the connections between its nodes to support a task at hand (J. Wu et al., 2019).

At the same time, the language network processes L1 and L2 by allocating more resources to the less automated and less familiar language, L2 (Malik-Moraleda et al., 2024; Wolna et al., 2024a), and it is insensitive to a temporary reversal of dominance, observed behaviorally in the LST. The effect of native vs. foreign language in the language system has been proposed to reflect general differences in the processing difficulty between L1 and L2: words in L2 are overall more difficult to access, as their baseline activation is lower (Casado et al., 2022; Green, 1998). As the activation level of a language is a difficult concept to capture experimentally, the difficulty in language processing has been operationalized using different measures that are supposed to approximate the effort in interpreting a given lexical stimulus. Relying on this approach, responses of the language system have been shown to be influenced by factors, such as predictability (stronger responses to more surprising items) and frequency (stronger responses to less frequent words (Tuckute et al., 2024)). In the population of late, unbalanced bilinguals, the dominant L1 constitutes a strong prior in language processing (for similar results in bilingual language comprehension, see Malik-Moraleda et al., 2022), and given a much longer and substantial exposure, it can be argued to have an overall larger frequency, since words in L1 are used and encountered more than those in L2.

Such factors might explain why the language system shows an overall stronger response to words in L2, without having to assume categorical distinctions between the two languages, such as “language tags” (Green, 1998), but by treating the level of processing difficulty as a continuous dimension, which might be influenced by multiple factors, in line with models proposing a shared selection mechanism between languages (Blanco-Elorrieta & Caramazza, 2021).

## 5. Conclusion

The present study investigated bilingual language switching as a lens for exploring neural mechanisms of bilingual language control, focusing on *language switch cost* and *reversed language dominance*. Our finding of *switch costs* in the MD network reinforces the view that bilingual language control is supported by domain-general cognitive mechanisms and that language switching shares overlapping neural mechanisms with nonverbal switching. In addition, we found no evidence that the BLC network is unique to language switching specifically and thus argue that BLC is subserved by a broader domain-general MD network. As for the *reversed language dominance*, our findings show that this effect is also reflected by the MD network. This suggests that *reversed language dominance* during language switching results from the engagement of control mechanisms rather than from changes in language-specific representations in the language network (in which we found a typical effect with L2 evoking stronger activation than L1). Overall, the two indices of bilingual language control found in LST (i.e. *language switch cost* and *reversed language dominance*) seem to be subserved by largely the same Multiple Demand system of the brain rather than the language network. As such, this suggests that behaviorally observed reversal of language dominance reflects increased demands of domain general control rather than reduced availability (or inhibition) of language-specific representations. These findings advance our understanding of how language-specific and general mechanisms interact to support bilingual language control.

## Acknowledgments

We want to express our sincere thanks to all members of the Psychology of Language and Bilingualism Laboratory, LangUsta, for their contributions to the research project and to the technical team at Center for Brain Research for their assistance with fMRI data acquisition. Special thanks are given to Kinga Masłowska, Piotr Górniak, and Anna Meliksetian for their help in data collection and coding. We also gratefully acknowledge the support of Poland’s high-performance computing infrastructure PLGrid (ACK Cyfronet Ares) for providing computer facilities and assistance under computational grant no. PLG/2024/017151. This research was made possible by an OPUS grant (UMO-2017/27/B/HS6/00959) from the National Science Center Poland awarded to Zofia Wodniecka.

## 6. Supplementary materials

The supplementary_materials.pdf contains details of the statistical information of the analysis as well as the results of individual regions of each network. Data and codes necessary to reproduce the results and plots presented in this paper are available at https://osf.io/g8723/overview. Raw neuroimaging data in the brain imaging data structure (BIDS-compliant format) are available at https://openneuro.org/datasets/ds006240.

