## supplementary materials for "Exploring Neural Mechanisms of Language Switching: An fMRI Study Using a Functional Localizer Approach"

**Table of Contents**

The Behavioral Results of the fMRI Session
- Language Switching Task
- Task Switching Task

Switch Costs within the Bilingual Language Control (BLC) System
- Functional Results of Pooled ROI (Localizer: Language Switching vs. Task Switching)
- Functional Results of Unpooled ROI (Localizer: Language Switching vs. Task Switching)
- Spatial Correlation of Pooled ROI
- Spatial Correlation of Unpooled ROI
- Spatial Overlap of Pooled ROI
- Spatial Overlap of Unpooled ROI

Switch Costs within the Multiple Demand (MD) System
- Functional Results of Pooled ROI (Localizer: Language Switching vs. Task Switching)
- Functional Results of Unpooled ROI (Localizer: Language Switching vs. Task Switching)
- Spatial Correlation of Pooled ROI
- Spatial Correlation of Unpooled ROI
- Spatial Overlap of Pooled ROI
- Spatial Overlap of Unpooled ROI

Language Switching within the Language and Multiple Demand (MD) Systems
- Functional Results of Pooled ROI
- Functional Results of Unpooled ROI

Experiment Stimuli

**The Behavioral Results of the Language Switching Task (LST) of the fMRI Session**

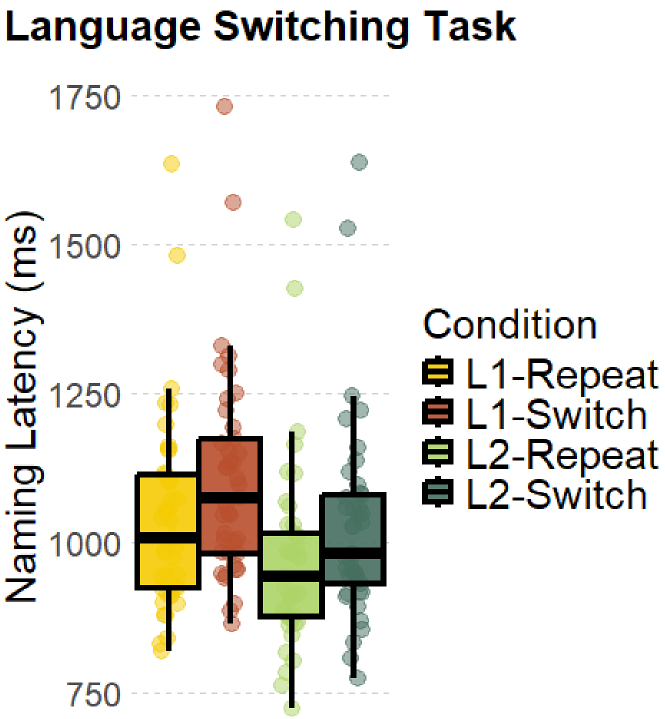


**Figure S1. Naming latencies in the Language Switching Task across Conditions.** The figure shows a boxplot of naming latencies in the LST across conditions. The horizontal lines of each box represent the first quartile (Q1), the median (Q2), and the third quartile (Q3), counted from the bottom. The interquartile range (IQR) is calculated as Q3 − Q1. The whiskers extend to the lower and upper bounds, defined as Q1 − 1.5 × IQR and Q3 + 1.5 × IQR, respectively.

**Table S1. Naming latencies in the Language Switching Task across Conditions.** The table presents the results of a mixed-effects model examining the effects of Language, Trial Type, and their interaction on naming latencies in the LST during the fMRI session.

| term | estimate | SE | t-value | p-value | CI |
|---|---|---|---|---|---|
| Intercept | 6.93 | 0.02 | 286.50 | < 0.001 | [6.88 6.98] |
| Language | -0.07 | 0.01 | -7.41 | < 0.001 | [-0.09 -0.05] |
| Trial Type | 0.06 | 0.004 | 14.49 | < 0.001 | [0.05 0.07] |
| Language x Trial Type | -0.002 | 0.01 | -0.27 | 0.79 | [-0.02 0.01] |

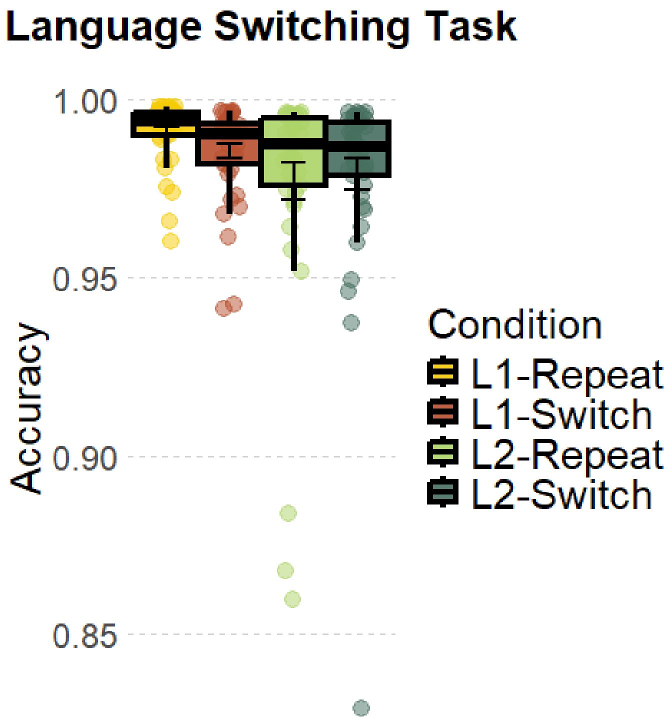


**Figure S2. Naming Accuracy in the Language Switching Task across Conditions.** The figure shows a boxplot of the naming accuracy in the LST across conditions. The horizontal lines of each box represent the first quartile (Q1), the median (Q2), and the third quartile (Q3), counted from the bottom. The interquartile range (IQR) is calculated as Q3 − Q1. The whiskers extend to the lower and upper bounds, defined as Q1 − 1.5 × IQR and Q3 + 1.5 × IQR, respectively.

**Table S2. Naming Accuracy in the Language Switching Task across Conditions.** The table presents the results of a mixed-effects model examining the effects of Language, Trial Type, and their interaction on the naming accuracy in the LST during the fMRI session.

| term | estimate | SE | z-value | p-value | CI |
|---|---|---|---|---|---|
| Intercept | 5.31 | 0.27 | 19.97 | < 0.001 | [4.79 5.83] |
| Language | -0.05 | 0.35 | -0.14 | 0.89 | [-0.74 0.64] |
| Trial Type | -0.32 | 0.21 | -1.51 | 0.13 | [-0.73 0.09] |
| Language x Trial Type | 0.40 | 0.37 | 1.08 | 0.28 | [-0.33 1.13] |

The Behavioral Results of the Task Switching (TS) of the fMRI Session

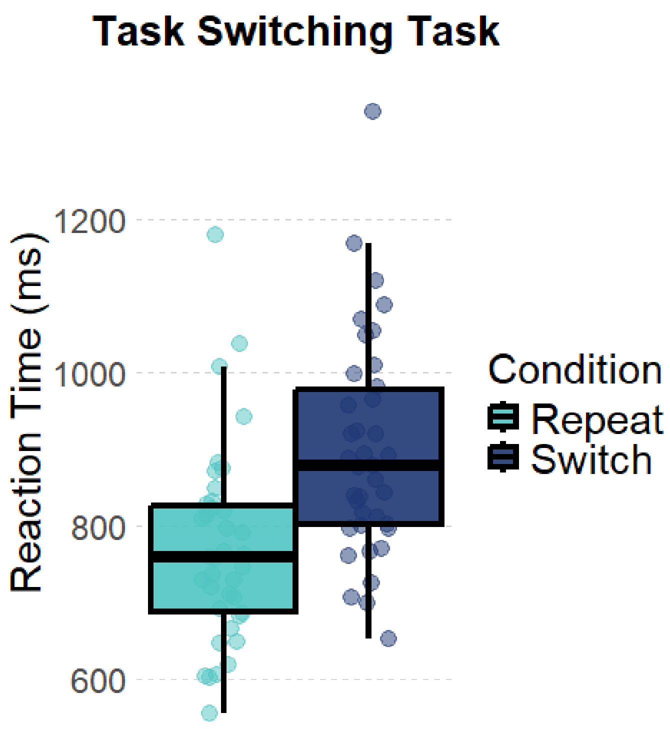


**Figure S3. Reaction Times in the Nonverbal Task Switching Task across Conditions.** The figure shows a boxplot of the reaction times in the nonverbal task switching task across conditions. The horizontal lines of each box represent the first quartile (Q1), the median (Q2), and the third quartile (Q3), counted from the bottom. The interquartile range (IQR) is calculated as Q3 − Q1. The whiskers extend to the lower and upper bounds, defined as Q1 − 1.5 × IQR and Q3 + 1.5 × IQR, respectively.

**Table S3. Reaction Times in the Nonverbal Task Switching Task across Conditions.** The table presents the results of a mixed-effects model examining the effects of Trial Type on the reaction times in the nonverbal task switching task during the fMRI session.

| term | estimate | SE | t-value | p-value | CI |
|---|---|---|---|---|---|
| Intercept | 6.71 | 0.03 | 263.06 | $< 0.001$ | [6.66 6.76] |
| Trial Type | 0.15 | 0.01 | 13.47 | $< 0.001$ | [0.13 0.18] |

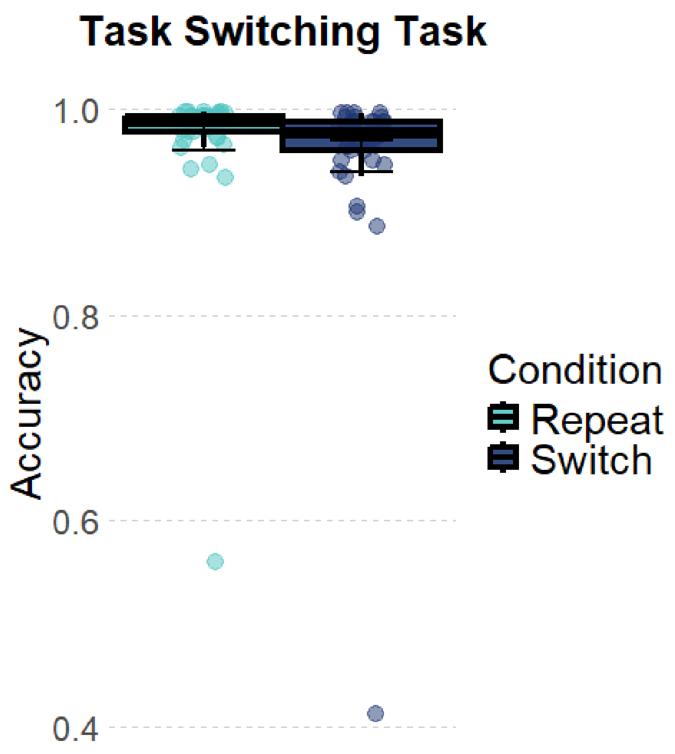


**Figure S4. Response Accuracy in the Nonverbal Task Switching Task across Conditions.** The figure shows a boxplot of the response accuracy in the nonverbal task switching task across conditions. The horizontal lines of each box represent the first quartile (Q1), the median (Q2), and the third quartile (Q3), counted from the bottom. The interquartile range (IQR) is calculated as Q3 − Q1. The whiskers extend to the lower and upper bounds, defined as Q1 − 1.5 × IQR and Q3 + 1.5 × IQR, respectively.

**Table S4. Response Accuracy in the Nonverbal Task Switching Task across Conditions.** The table presents the results of a mixed-effects model examining the effects of Trial Type on the response accuracy in the nonverbal task switching task during the fMRI session.

| term | estimate | SE | z-value | p-value | CI |
|---|---|---|---|---|---|
| Intercept | 4.05 | 0.22 | 18.19 | < 0.001 | [3.61 4.49] |
| Trial Type | -0.59 | 0.13 | -4.70 | < 0.001 | [-0.84 -0.34] |

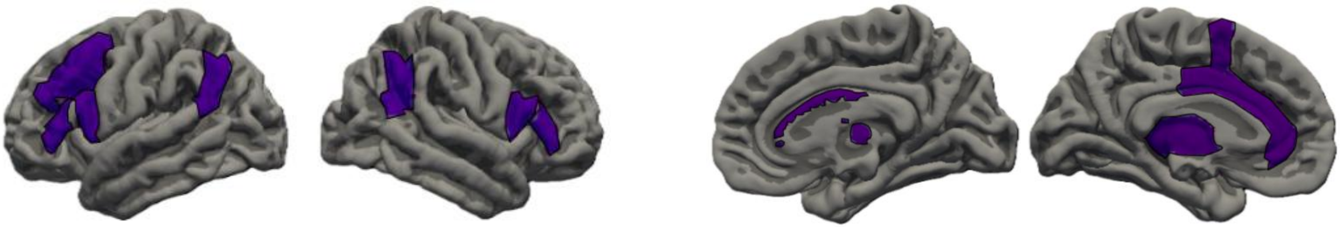

**Figure S5. The Bilingual Language Control Parcels.** The BLC parcels are created based on Abutalebi and Green (2016). The two plots on the left display the lateral view, whereas the two plots on the right display the medial view.

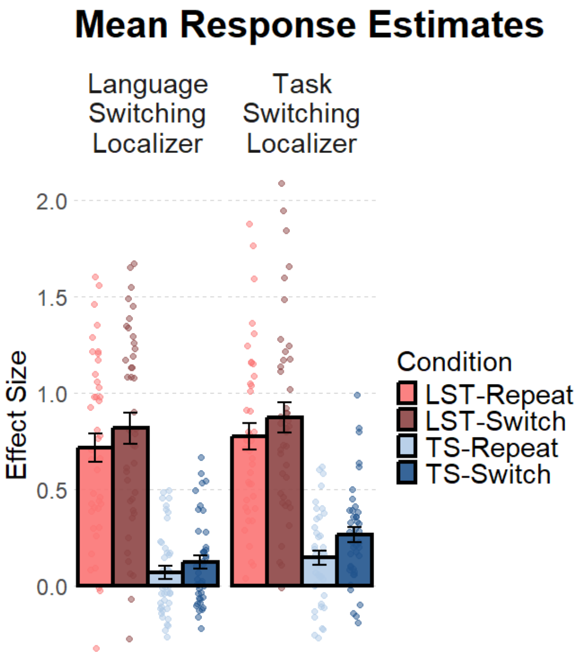


**Figure S6. Mean Response Estimates across Tasks and Trial Types.** The figure shows a bar plot of the mean response estimates, corresponding to the % BOLD signal change, across tasks and trial types. The 4 bars in the left panel depict results with the language-switching localizer, and the 4 bars in the right panel depict results with the nonverbal task-switching localizer. LST: language switching task, TS: nonverbal task switching task.

**Table S5. Mean Response Estimates across Tasks and Trial Types.** The table presents the results of a mixed-effects model examining the effects of Task, Trial Type, and their interaction on response estimates, using the language-switching localizer.

**Language Switching Localizer (BLC)**

| term | estimate | SE | t-value | p-value | CI |
|---|---|---|---|---|---|
| Intercept | 0.43 | 0.10 | 4.35 | < 0.001 | [0.22 0.64] |
| Task | -0.67 | 0.15 | -4.47 | < 0.001 | [-0.99 -0.36] |
| Trial Type | 0.08 | 0.03 | 2.68 | < 0.05 | [0.02 0.14] |
| Task x Trial Type | -0.05 | 0.05 | -0.89 | 0.38 | [-0.15 0.06] |

**Table S6. Mean Response Estimates across Tasks and Trial Types.** The table presents the results of a mixed-effects model examining the effects of Task, Trial Type, and their interaction on response estimates, using the nonverbal task-switching localizer.

**Task Switching Localizer (BLC)**

| term | estimate | SE | t-value | p-value | CI |
|---|---|---|---|---|---|
| Intercept | 0.51 | 0.09 | 5.75 | < 0.001 | [0.33 0.70] |
| Task | -0.62 | 0.13 | -4.60 | < 0.001 | [-0.90 -0.34] |
| Trial Type | 0.11 | 0.03 | 3.08 | < 0.01 | [0.04 0.18] |
| Task x Trial Type | 0.02 | 0.05 | 0.48 | 0.63 | [-0.07 0.11] |

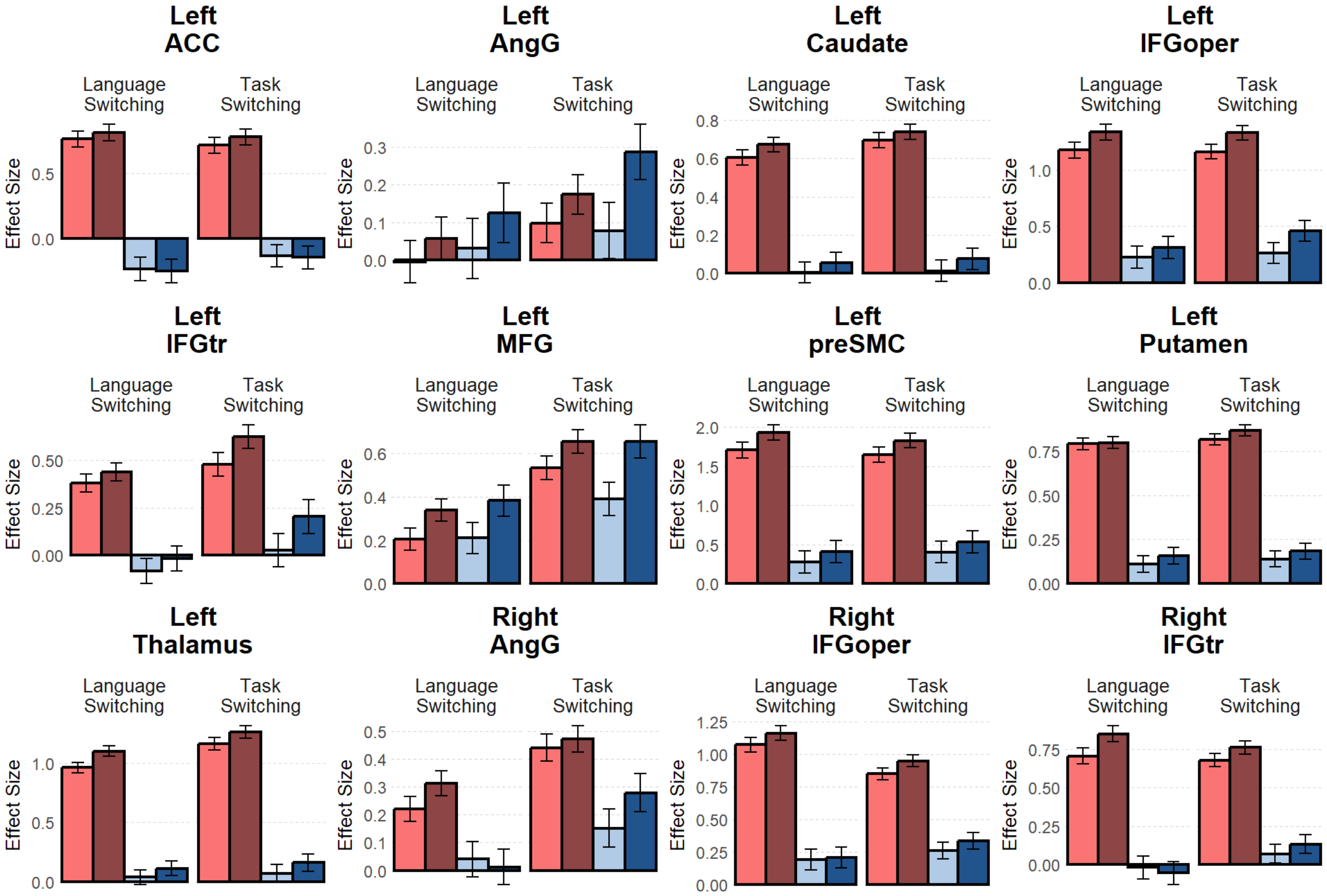


**Figure S7 By-ROI Response Estimates within the BLC Parcels.** The figure shows bar plots of mean response estimates, expressed as % BOLD signal change, across tasks and trial types. Each sub-figure corresponds to an individual fROI within the BLC parcels. Within each sub-figure, the 4 bars in the left panel depict results from the language-switching localizer, and the 4 bars in the right panel depict results from the nonverbal task-switching localizer.

**Table S7. By-ROI Response Estimates within BLC Parcels Using the Language-Switching Localizer.** The table presents the by-ROI results of a mixed-effects model examining the effects of Task, Trial Type, and their interaction on response estimates, using the language-switching localizer.

| ROI | term | estimate | SE | t-value | p-value | CI |
|---|---|---|---|---|---|---|
| leftACC | Intercept | 0.27 | 0.10 | 2.68 | < 0.05 | [0.07 0.48] |
| leftACC | Task | -1.04 | 0.07 | -14.76 | < 0.001 | [-1.18 -0.90] |
| leftACC | Trial Type | 0.02 | 0.07 | 0.25 | 0.80 | [-0.12 0.16] |
| leftACC | Task x Trial Type | -0.07 | 0.14 | -0.48 | 0.63 | [-0.34 0.21] |
| leftAngG | Intercept | 0.05 | 0.09 | 0.59 | 0.56 | [-0.13 0.23] |
| leftAngG | Task | 0.05 | 0.06 | 0.88 | 0.38 | [-0.06 0.17] |
| leftAngG | Trial Type | 0.08 | 0.06 | 1.34 | 0.18 | [-0.04 0.19] |
| leftAngG | Task x Trial Type | 0.03 | 0.12 | 0.27 | 0.78 | [-0.20 0.26] |
| leftCaudate | Intercept | 0.34 | 0.07 | 5.07 | < 0.001 | [0.20 0.47] |
| leftCaudate | Task | -0.61 | 0.06 | -10.99 | < 0.001 | [-0.72 -0.50] |

| ROI | term | estimate | SE | t-value | p-value | CI |
|---|---|---|---|---|---|---|
| leftCaudate | Trial Type | 0.06 | 0.06 | 1.06 | 0.29 | [-0.05 0.17] |
| leftCaudate | Task x Trial Type | -0.02 | 0.11 | -0.14 | 0.89 | [-0.23 0.20] |
| leftIFGoper | Intercept | 0.76 | 0.11 | 6.70 | < 0.001 | [0.53 0.99] |
| leftIFGoper | Task | -0.98 | 0.09 | -11.41 | < 0.001 | [-1.15 -0.81] |
| leftIFGoper | Trial Type | 0.12 | 0.09 | 1.43 | 0.15 | [-0.05 0.29] |
| leftIFGoper | Task x Trial Type | -0.07 | 0.17 | -0.42 | 0.68 | [-0.41 0.27] |
| leftIFGtr | Intercept | 0.18 | 0.08 | 2.24 | < 0.05 | [0.02 0.34] |
| leftIFGtr | Task | -0.46 | 0.07 | -6.80 | < 0.001 | [-0.59 -0.33] |
| leftIFGtr | Trial Type | 0.06 | 0.07 | 0.92 | 0.36 | [-0.07 0.20] |
| leftIFGtr | Task x Trial Type | 0.005 | 0.14 | 0.04 | 0.97 | [-0.26 0.27] |
| leftMFG | Intercept | 0.28 | 0.08 | 3.60 | < 0.001 | [0.12 0.44] |
| leftMFG | Task | 0.03 | 0.05 | 0.51 | 0.61 | [-0.07 0.12] |
| leftMFG | Trial Type | 0.15 | 0.05 | 3.11 | < 0.01 | [0.06 0.25] |
| leftMFG | Task x Trial Type | 0.04 | 0.10 | 0.38 | 0.70 | [-0.16 0.23] |
| leftpreSMC | Intercept | 1.08 | 0.16 | 6.93 | < 0.001 | [0.76 1.39] |
| leftpreSMC | Task | -1.48 | 0.10 | -15.35 | < 0.001 | [-1.67 -1.29] |
| leftpreSMC | Trial Type | 0.18 | 0.10 | 1.86 | 0.06 | [-0.01 0.37] |
| leftpreSMC | Task x Trial Type | -0.09 | 0.19 | -0.46 | 0.65 | [-0.47 0.29] |
| leftPutamen | Intercept | 0.47 | 0.06 | 8.55 | < 0.001 | [0.36 0.58] |
| leftPutamen | Task | -0.66 | 0.04 | -16.36 | < 0.001 | [-0.74 -0.58] |
| leftPutamen | Trial Type | 0.03 | 0.04 | 0.66 | 0.51 | [-0.05 0.11] |
| leftPutamen | Task x Trial Type | 0.04 | 0.08 | 0.51 | 0.61 | [-0.12 0.20] |
| leftThalamus | Intercept | 0.55 | 0.07 | 7.44 | < 0.001 | [0.40 0.70] |
| leftThalamus | Task | -0.96 | 0.06 | -15.84 | < 0.001 | [-1.07 -0.84] |
| leftThalamus | Trial Type | 0.10 | 0.06 | 1.75 | 0.08 | [-0.01 0.22] |
| leftThalamus | Task x Trial Type | -0.06 | 0.12 | -0.52 | 0.61 | [-0.30 0.18] |
| rightAngG | Intercept | 0.15 | 0.07 | 2.03 | < 0.05 | [0.0009 0.29] |
| rightAngG | Task | -0.24 | 0.05 | -4.58 | < 0.001 | [-0.34 -0.14] |
| rightAngG | Trial Type | 0.03 | 0.05 | 0.60 | 0.55 | [-0.07 0.14] |
| rightAngG | Task x Trial Type | -0.12 | 0.10 | -1.14 | 0.26 | [-0.33 0.09] |
| rightIFGoper | Intercept | 0.66 | 0.09 | 7.21 | < 0.001 | [0.47 0.84] |
| rightIFGoper | Task | -0.92 | 0.07 | -13.88 | < 0.001 | [-1.05 -0.79] |
| rightIFGoper | Trial Type | 0.05 | 0.07 | 0.81 | 0.42 | [-0.08 0.18] |

Table of Contents

| ROI | term | estimate | SE | t-value | p-value | CI |
|---|---|---|---|---|---|---|
| rightIFGoper | Task x Trial Type | -0.07 | 0.13 | -0.54 | 0.59 | [-0.33 0.19] |
| rightIFGtr | Intercept | 0.37 | 0.09 | 4.21 | < 0.001 | [0.19 0.55] |
| rightIFGtr | Task | -0.81 | 0.07 | -11.97 | < 0.001 | [-0.95 -0.68] |
| rightIFGtr | Trial Type | 0.05 | 0.07 | 0.80 | 0.43 | [-0.08 0.19] |
| rightIFGtr | Task x Trial Type | -0.18 | 0.14 | -1.32 | 0.19 | [-0.45 0.09] |

**Table S8. By-ROI Response Estimates within BLC Parcels Using the Nonverbal Task-Switching Localizer.** The table presents the by-ROI results of a mixed-effects model examining the effects of Task, Trial Type, and their interaction on response estimates, using the nonverbal task-switching localizer.

| ROI | term | estimate | SE | t-value | p-value | CI |
|---|---|---|---|---|---|---|
| leftACC | Intercept | 0.30 | 0.10 | 3.10 | < 0.01 | [0.11 0.50] |
| leftACC | Task | -0.89 | 0.06 | -13.78 | < 0.001 | [-1.02 -0.76] |
| leftACC | Trial Type | 0.03 | 0.06 | 0.40 | 0.69 | [-0.10 0.15] |
| leftACC | Task x Trial Type | -0.08 | 0.13 | -0.59 | 0.56 | [-0.33 0.18] |
| leftAngG | Intercept | 0.16 | 0.09 | 1.88 | 0.07 | [-0.01 0.33] |
| leftAngG | Task | 0.05 | 0.06 | 0.73 | 0.46 | [-0.08 0.17] |
| leftAngG | Trial Type | 0.14 | 0.06 | 2.26 | < 0.05 | [0.02 0.27] |
| leftAngG | Task x Trial Type | 0.13 | 0.13 | 1.05 | 0.29 | [-0.12 0.38] |
| leftCaudate | Intercept | 0.38 | 0.07 | 5.71 | < 0.001 | [0.25 0.52] |
| leftCaudate | Task | -0.67 | 0.05 | -12.40 | < 0.001 | [-0.78 -0.57] |
| leftCaudate | Trial Type | 0.05 | 0.05 | 0.99 | 0.32 | [-0.05 0.16] |
| leftCaudate | Task x Trial Type | 0.02 | 0.11 | 0.18 | 0.86 | [-0.19 0.23] |
| leftIFGoper | Intercept | 0.80 | 0.11 | 7.52 | < 0.001 | [0.59 1.02] |
| leftIFGoper | Task | -0.88 | 0.08 | -10.62 | < 0.001 | [-1.04 -0.72] |
| leftIFGoper | Trial Type | 0.18 | 0.08 | 2.21 | < 0.05 | [0.02 0.35] |
| leftIFGoper | Task x Trial Type | 0.03 | 0.17 | 0.15 | 0.88 | [-0.30 0.35] |

| ROI | term | estimate | SE | t-value | p-value | CI |
|---|---|---|---|---|---|---|
| leftIFGtr | Intercept | 0.33 | 0.10 | 3.36 | < 0.01 | [0.13 0.53] |
| leftIFGtr | Task | -0.44 | 0.07 | -6.43 | < 0.001 | [-0.57 -0.30] |
| leftIFGtr | Trial Type | 0.16 | 0.07 | 2.38 | < 0.05 | [0.03 0.30] |
| leftIFGtr | Task x Trial Type | 0.03 | 0.14 | 0.23 | 0.81 | [-0.24 0.30] |
| leftMFG | Intercept | 0.56 | 0.09 | 6.43 | < 0.001 | [0.38 0.74] |
| leftMFG | Task | -0.07 | 0.06 | -1.23 | 0.22 | [-0.19 0.04] |
| leftMFG | Trial Type | 0.19 | 0.06 | 3.26 | < 0.001 | [0.08 0.31] |
| leftMFG | Task x Trial Type | 0.14 | 0.12 | 1.21 | 0.23 | [-0.09 0.38] |
| leftpreSMC | Intercept | 1.10 | 0.15 | 7.33 | < 0.001 | [0.80 1.41] |
| leftpreSMC | Task | -1.27 | 0.09 | -13.98 | < 0.001 | [-1.45 -1.09] |
| leftpreSMC | Trial Type | 0.16 | 0.09 | 1.71 | 0.09 | [-0.02 0.34] |
| leftpreSMC | Task x Trial Type | -0.05 | 0.18 | -0.26 | 0.79 | [-0.41 0.31] |
| leftPutamen | Intercept | 0.50 | 0.05 | 9.67 | < 0.001 | [0.40 0.61] |
| leftPutamen | Task | -0.68 | 0.04 | -16.73 | < 0.001 | [-0.76 -0.60] |
| leftPutamen | Trial Type | 0.05 | 0.04 | 1.16 | 0.25 | [-0.03 0.13] |
| leftPutamen | Task x Trial Type | -0.01 | 0.08 | -0.08 | 0.94 | [-0.17 0.16] |
| leftThalamus | Intercept | 0.66 | 0.08 | 8.01 | < 0.001 | [0.50 0.83] |
| leftThalamus | Task | -1.10 | 0.06 | -18.38 | < 0.001 | [-1.22 -0.98] |
| leftThalamus | Trial Type | 0.10 | 0.06 | 1.58 | 0.12 | [-0.02 0.21] |
| leftThalamus | Task x Trial Type | -0.01 | 0.12 | -0.09 | 0.93 | [-0.25 0.22] |
| rightAngG | Intercept | 0.34 | 0.08 | 4.34 | < 0.001 | [0.18 0.49] |
| rightAngG | Task | -0.24 | 0.06 | -4.33 | < 0.001 | [-0.35 -0.13] |
| rightAngG | Trial Type | 0.08 | 0.06 | 1.42 | 0.16 | [-0.03 0.19] |

| ROI | term | estimate | SE | t-value | p-value | CI |
|---|---|---|---|---|---|---|
| rightAngG | Task x Trial Type | 0.10 | 0.11 | 0.86 | 0.39 | [-0.12 0.32] |
| rightIFGoper | Intercept | 0.60 | 0.07 | 8.10 | < 0.001 | [0.45 0.75] |
| rightIFGoper | Task | -0.60 | 0.06 | -10.64 | < 0.001 | [-0.71 -0.49] |
| rightIFGoper | Trial Type | 0.09 | 0.06 | 1.50 | 0.13 | [-0.03 0.20] |
| rightIFGoper | Task x Trial Type | -0.03 | 0.11 | -0.24 | 0.81 | [-0.25 0.20] |
| rightIFGtr | Intercept | 0.41 | 0.07 | 5.68 | < 0.001 | [0.26 0.56] |
| rightIFGtr | Task | -0.62 | 0.06 | -10.45 | < 0.001 | [-0.74 -0.50] |
| rightIFGtr | Trial Type | 0.07 | 0.06 | 1.26 | 0.21 | [-0.04 0.19] |
| rightIFGtr | Task x Trial Type | -0.02 | 0.12 | -0.17 | 0.87 | [-0.25 0.21] |

### Spatial Correlation of Switch Costs in the BLC System

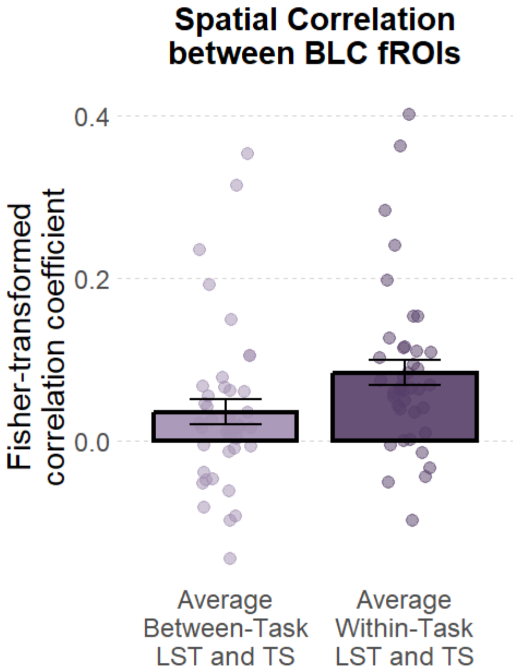


**Figure S8. Spatial Correlation between BLC fROIs.** The figure shows a bar plot of the fisher-transformed correlation coefficient between BLC fROIs. For the between-task condition, we computed the correlation coefficients across different tasks (e.g., LST-run1 with TS-run1) and averaged the values. For the within-task condition, we computed the correlation coefficients between runs of the same task (e.g., LST-run1 with LST-run2) and averaged the two values. LST: language switching task, TS: nonverbal task switching task.

**Table S9. Spatial Correlation between BLC fROIs.** The table presents the results of a mixed-effects model examining the effects of Task Type on spatial correlation between the BLC fROIs.

| term | estimate | SE | t-value | p-value | CI |
|---|---|---|---|---|---|
| Intercept | 0.06 | 0.02 | 2.91 | < 0.01 | [0.02 0.10] |
| Task Type | -0.05 | 0.01 | -3.48 | < 0.001 | [-0.07 -0.02] |

**Figure S9. By-ROI Spatial Correlation between BLC fROIs.** The figure shows bar plots of the fisher-transformed correlation coefficient in BLC fROIs. For the between-task condition, we computed the correlation coefficients across different tasks (e.g., LST-run1 with TS-run1) and averaged the values. For the within-task condition, we computed the correlation coefficients between runs of the same task (e.g., LST-run1 with LST-run2) and averaged the two values. LST: language switching task, TS: nonverbal task switching task.

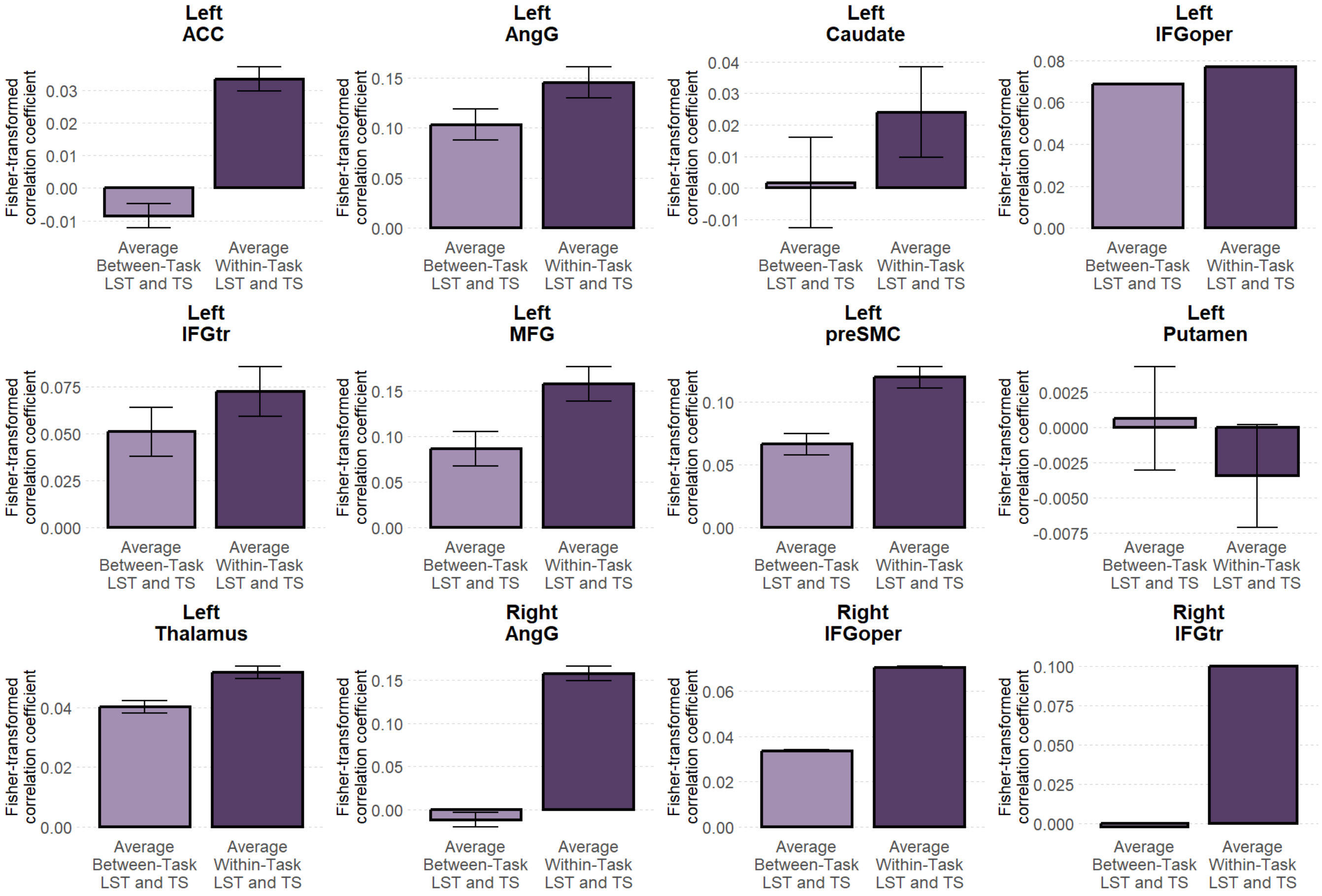


**Table S10. By-ROI Spatial Correlation between BLC fROIs.** The table presents the by-ROI results of a mixed-effects model examining the effects of Task Type on spatial correlation in each BLC fROI.

| ROI | term | estimate | SE | t-value | p-value | CI |
|---|---|---|---|---|---|---|
| leftACC | Intercept | 0.01 | 0.02 | 0.54 | 0.59 | [-0.03 0.06] |
| leftACC | Task Task | -0.04 | 0.04 | -0.98 | 0.33 | [-0.13 0.04] |
| leftAngG | Intercept | 0.12 | 0.03 | 3.69 | < 0.001 | [0.06 0.19] |
| leftAngG | Task Task | -0.04 | 0.05 | -0.86 | 0.40 | [-0.14 0.06] |
| leftCaudate | Intercept | 0.01 | 0.02 | 0.55 | 0.58 | [-0.03 0.06] |
| leftCaudate | Task Task | -0.02 | 0.03 | -0.78 | 0.44 | [-0.08 0.04] |

Table of Contents

| ROI | term | estimate | SE | t-value | p-value | CI |
|---|---|---|---|---|---|---|
| leftIFGoper | Intercept | 0.07 | 0.04 | 2.10 | < 0.05 | [0.004 0.14] |
| leftIFGoper | Task Task | -0.01 | 0.07 | -0.12 | 0.91 | [-0.15 0.13] |
| leftIFGtr | Intercept | 0.06 | 0.03 | 1.96 | 0.06 | [-0.002 0.13] |
| leftIFGtr | Task Task | -0.02 | 0.05 | -0.45 | 0.66 | [-0.12 0.08] |
| leftMFG | Intercept | 0.12 | 0.03 | 3.88 | < 0.001 | [0.06 0.19] |
| leftMFG | Task Task | -0.07 | 0.04 | -1.78 | 0.08 | [-0.15 0.01] |
| leftpreSMC | Intercept | 0.09 | 0.04 | 2.38 | < 0.05 | [0.01 0.17] |
| leftpreSMC | Task Task | -0.05 | 0.07 | -0.77 | 0.45 | [-0.19 0.09] |
| leftPutamen | Intercept | -0.001 | 0.01 | -0.13 | 0.90 | [-0.02 0.02] |
| leftPutamen | Task Task | 0.004 | 0.02 | 0.24 | 0.81 | [-0.03 0.04] |
| leftThalamus | Intercept | 0.05 | 0.02 | 2.09 | < 0.05 | [0.002 0.09] |
| leftThalamus | Task Task | -0.01 | 0.04 | -0.27 | 0.79 | [-0.10 0.07] |
| rightAngG | Intercept | 0.07 | 0.03 | 2.86 | < 0.01 | [0.02 0.12] |
| rightAngG | Task Task | -0.17 | 0.04 | -4.05 | < 0.001 | [-0.25 -0.09] |
| rightIFGoper | Intercept | 0.05 | 0.03 | 1.86 | 0.07 | [-0.004 0.11] |
| rightIFGoper | Task Task | -0.04 | 0.06 | -0.66 | 0.51 | [-0.15 0.07] |
| rightIFGtr | Intercept | 0.05 | 0.03 | 1.65 | 0.10 | [-0.01 0.11] |
| rightIFGtr | Task Task | -0.10 | 0.06 | -1.73 | 0.09 | [-0.22 0.02] |

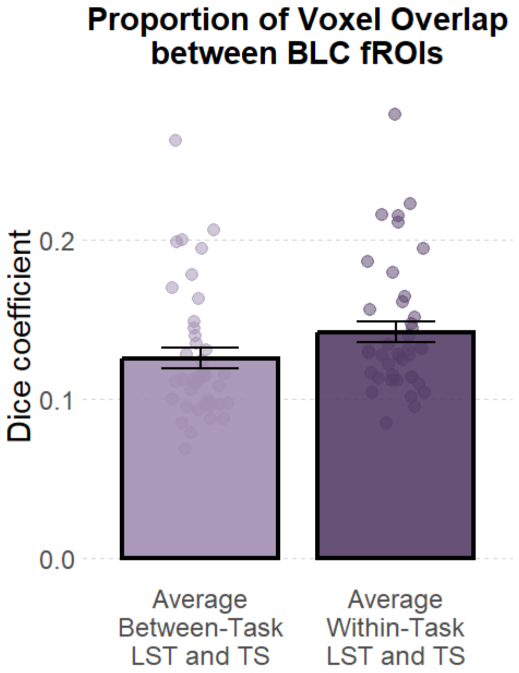


**Figure S10. Proportion of Spatial Overlap between BLC fROIs.** The figure shows a bar plot of the proportion of spatial overlap between BLC fROIs. For the between-task condition, we computed the dice coefficients across different tasks (e.g., LST-run1 with TS-run1) and averaged the values. For the within-task condition, we computed the dice coefficients between runs of the same task (e.g., LST-run1 with LST-run2) and averaged the two values. LST: language switching task, TS: nonverbal task switching task.

**Table S11. Proportion of Spatial Overlap between BLC fROIs.** The table presents the results of a mixed-effects model examining the effects of Task Type on spatial overlap between the BLC fROIs.

| term | estimate | SE | t-value | p-value | CI |
|---|---|---|---|---|---|
| Intercept | 0.13 | 0.01 | 15.76 | < 0.001 | [0.12 0.15] |
| Task Type | -0.02 | 0.01 | -2.76 | < 0.01 | [-0.03 -0.005] |

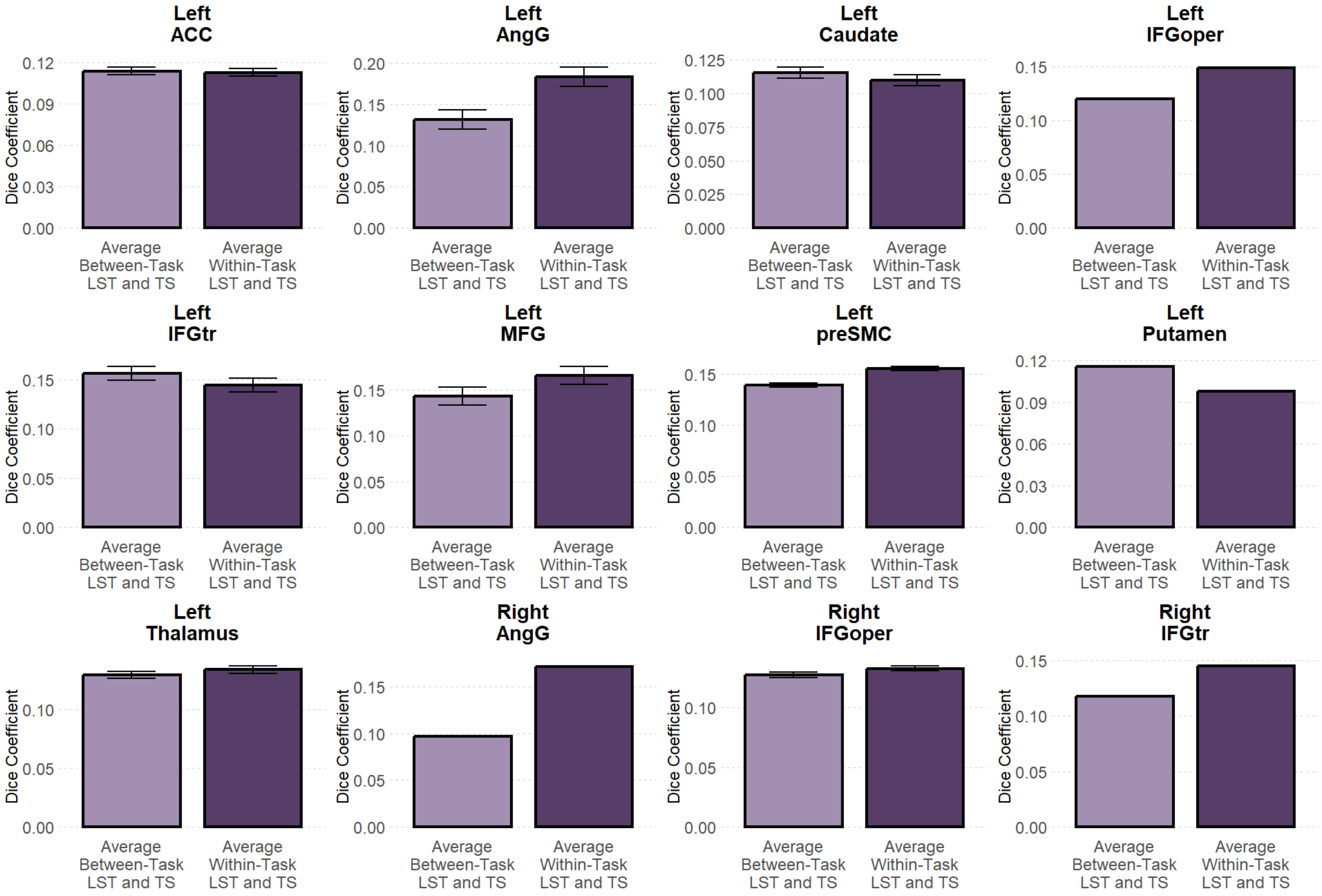


**Figure S11. By-ROI Proportion of Spatial Overlap in BLC fROIs.** The figure shows bar plots of the dice coefficient in BLC fROIs. For the between-task condition, we computed the dice coefficients across different tasks (e.g., LST-run1 with TS-run1) and averaged the values. For the within-task condition, we computed the dice coefficients between runs of the same task (e.g., LST-run1 with LST-run2) and averaged the two values. LST: language switching task, TS: nonverbal task switching task.

**Table S12. By-ROI Proportion of Spatial Overlap in BLC fROIs.** The table presents the by-ROI results of a mixed-effects model examining the effects of Task Type on proportion of spatial overlap in each BLC fROI.

| ROI | term | estimate | SE | t-value | p-value | CI |
|---|---|---|---|---|---|---|
| leftACC | Intercept | 0.11 | 0.01 | 12.30 | < 0.001 | [0.10 0.13] |
| leftACC | Task Type | 0.001 | 0.01 | 0.07 | 0.95 | [-0.03 0.03] |
| leftAngG | Intercept | 0.16 | 0.02 | 8.55 | < 0.001 | [0.12 0.20] |
| leftAngG | Task Type | -0.05 | 0.02 | -2.32 | < 0.05 | [-0.10 -0.01] |
| leftCaudate | Intercept | 0.11 | 0.01 | 11.38 | < 0.001 | [0.09 0.13] |
| leftCaudate | Task Type | 0.005 | 0.01 | 0.36 | 0.72 | [-0.03 0.04] |
| leftIFGoper | Intercept | 0.13 | 0.01 | 12.76 | < 0.001 | [0.11 0.16] |
| leftIFGoper | Task Type | -0.03 | 0.02 | -1.42 | 0.16 | [-0.07 0.01] |
| leftIFGtr | Intercept | 0.15 | 0.02 | 9.17 | < 0.001 | [0.12 0.18] |

Table of Contents

| ROI | term | estimate | SE | t-value | p-value | CI |
|---|---|---|---|---|---|---|
| leftIFGtr | Task Type | 0.01 | 0.03 | 0.48 | 0.64 | [-0.04 0.06] |
| leftMFG | Intercept | 0.16 | 0.01 | 10.15 | < 0.001 | [0.12 0.18] |
| leftMFG | Task Type | -0.02 | 0.02 | -1.25 | 0.22 | [-0.06 0.01] |
| leftpreSMC | Intercept | 0.15 | 0.01 | 10.46 | < 0.001 | [0.12 0.18] |
| leftpreSMC | Task Type | -0.02 | 0.03 | -0.62 | 0.54 | [-0.07 0.04] |
| leftPutamen | Intercept | 0.11 | 0.01 | 18.08 | < 0.001 | [0.10 0.12] |
| leftPutamen | Task Type | 0.02 | 0.01 | 1.51 | 0.14 | [-0.01 0.04] |
| leftThalamus | Intercept | 0.13 | 0.01 | 13.61 | < 0.001 | [0.11 0.15] |
| leftThalamus | Task Type | -0.005 | 0.02 | -0.29 | 0.78 | [-0.04 0.03] |
| rightAngG | Intercept | 0.14 | 0.01 | 12.61 | < 0.001 | [0.11 0.16] |
| rightAngG | Task Type | -0.07 | 0.02 | -3.56 | < 0.001 | [-0.12 -0.03] |
| rightIFGoper | Intercept | 0.13 | 0.01 | 11.16 | < 0.001 | [0.11 0.15] |
| rightIFGoper | Task Type | -0.005 | 0.02 | -0.26 | 0.80 | [-0.05 0.04] |
| rightIFGtr | Intercept | 0.13 | 0.01 | 11.06 | < 0.001 | [0.11 0.16] |
| rightIFGtr | Task Type | -0.03 | 0.02 | -1.14 | 0.26 | [-0.07 0.02] |

#### Mean Response estimates collapsed over MD ROIs

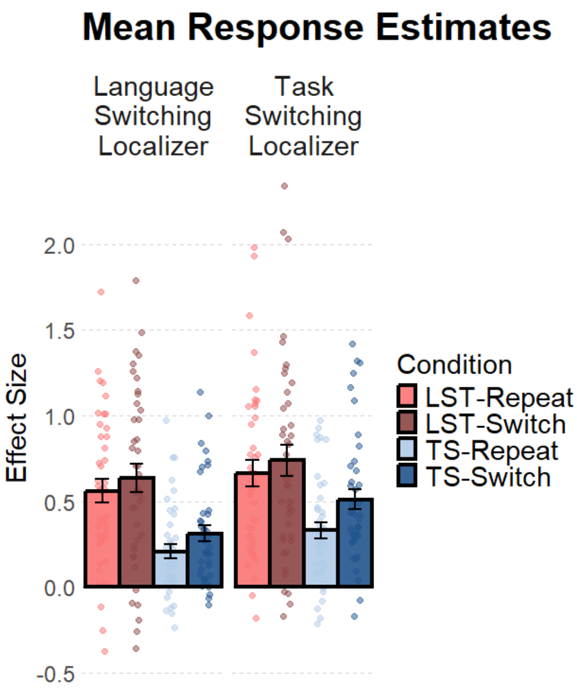


**Figure S12. Mean Response Estimates across Tasks and Trial Types.** The figure shows a bar plot of the mean response estimates, corresponding to the % BOLD signal change, across tasks and trial types. The 4 bars in the left panel depict results with the language-switching localizer, and 4 bars in the right panel depict results with the nonverbal task-switching localizer. LST: language switching task, TS: nonverbal task switching task.

**Table S13. Mean Response Estimates across Tasks and Trial Types.** The table presents the results of a mixed-effects model examining the effects of Task, Trial Type, and their interaction on response estimates, using the language-switching localizer.

| term | estimate | SE | t-value | p-value | CI |
|---|---|---|---|---|---|
| Intercept | 0.43 | 0.07 | 6.39 | < 0.001 | [0.29 0.56] |
| Task | -0.34 | 0.10 | -3.25 | < 0.01 | [-0.55 -0.13] |
| Trial Type | 0.09 | 0.03 | 3.22 | < 0.01 | [0.03 0.15] |
| Task x Trial Type | 0.03 | 0.05 | 0.66 | 0.51 | [-0.06 0.12] |

**Table S14. Mean Response Estimates across Tasks and Trial Types.** The table presents the results of a mixed-effects model examining the effects of Task, Trial Type, and their interaction on response estimates,

Table of Contents

using the task-switching localizer.

| **term** | **estimate** | **SE** | **t-value** | **p-value** | **CI** |
|---|---|---|---|---|---|
| Intercept | 0.56 | 0.08 | 6.99 | < 0.001 | [0.40 0.72] |
| Task | -0.28 | 0.10 | -2.81 | < 0.01 | [-0.48 -0.08] |
| Trial Type | 0.13 | 0.03 | 3.80 | < 0.001 | [0.06 0.20] |
| Task x Trial Type | 0.10 | 0.03 | 3.11 | < 0.01 | [0.04 0.17] |

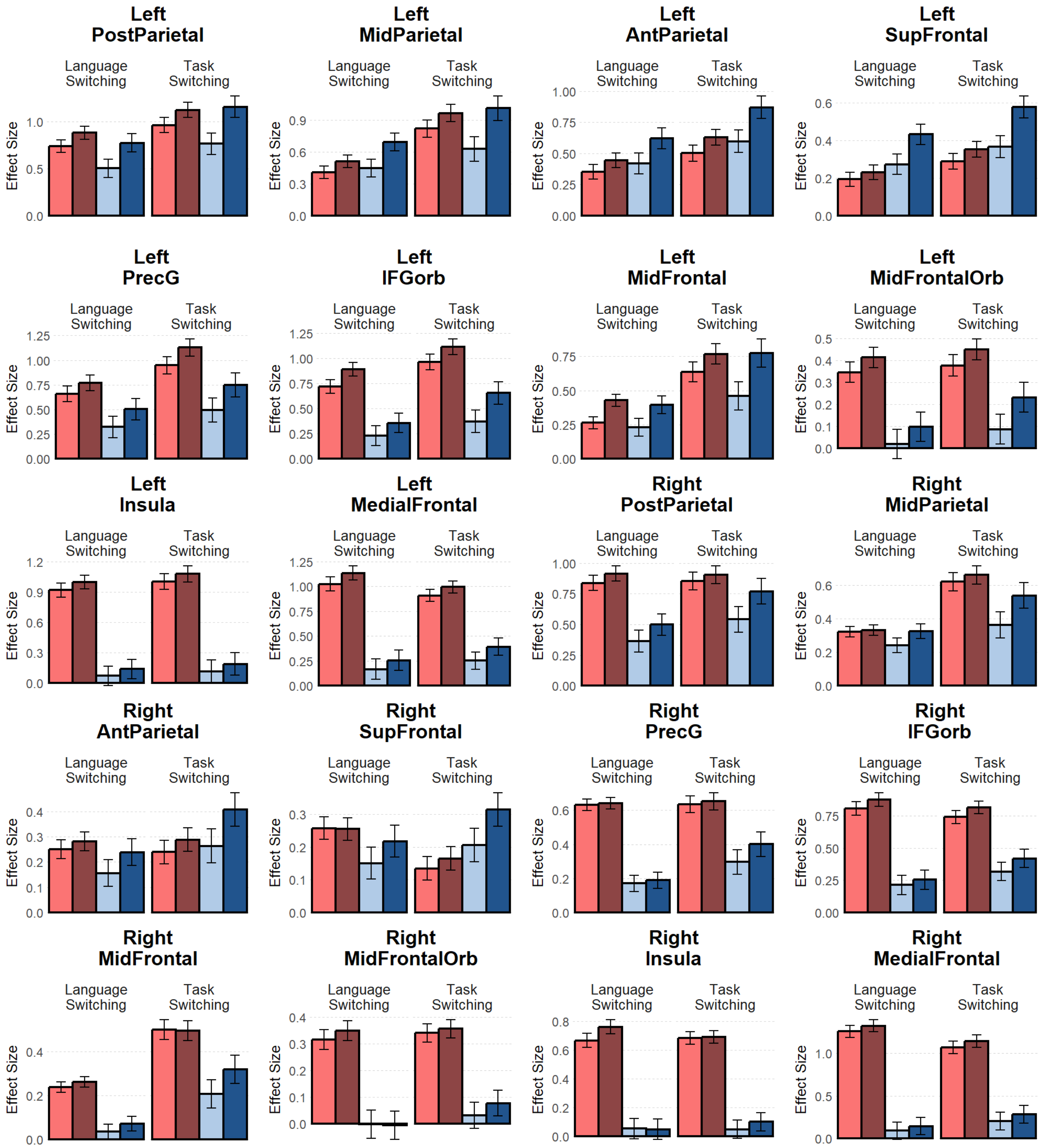


**Figure S13. By-ROI Response Estimates within the MD Parcels.** The figure shows bar plots of mean response estimates, expressed as % BOLD signal change, across tasks and trial types. Each sub-figure corresponds to an individual fROI within the MD parcels. Within each sub-figure, the 4 bars in the left panel depict results from the language-switching localizer, and the 4 bars in the right panel depict results from the nonverbal task-switching localizer.

**Table S15. By-ROI Response Estimates within MD Parcels Using the Language-Switching Localizer.** The

Table of Contents

table presents the by-ROI results of a mixed-effects model examining the effects of Task, Trial Type, and their interaction on response estimates, using the language-switching localizer.

| ROI | term | estimate | SE | t-value | p-value | CI |
|---|---|---|---|---|---|---|
| leftPostParietal | Intercept | 0.72 | 0.10 | 6.87 | < 0.001 | [0.51 0.94] |
| leftPostParietal | Task | -0.17 | 0.06 | -2.84 | < 0.01 | [-0.29 -0.05] |
| leftPostParietal | Trial Type | 0.21 | 0.06 | 3.38 | < 0.001 | [0.09 0.32] |
| leftPostParietal | Task x Trial Type | 0.13 | 0.12 | 1.05 | 0.30 | [-0.11 0.37] |
| leftMidParietal | Intercept | 0.52 | 0.09 | 5.52 | < 0.001 | [0.33 0.71] |
| leftMidParietal | Task | 0.11 | 0.06 | 1.85 | 0.06 | [-0.01 0.22] |
| leftMidParietal | Trial Type | 0.18 | 0.06 | 2.98 | < 0.01 | [0.06 0.29] |
| leftMidParietal | Task x Trial Type | 0.14 | 0.12 | 1.20 | 0.23 | [-0.09 0.37] |
| leftAntParietal | Intercept | 0.46 | 0.09 | 5.09 | < 0.001 | [0.28 0.64] |
| leftAntParietal | Task | 0.12 | 0.05 | 2.52 | < 0.05 | [0.03 0.22] |
| leftAntParietal | Trial Type | 0.15 | 0.05 | 3.02 | < 0.01 | [0.05 0.24] |
| leftAntParietal | Task x Trial Type | 0.11 | 0.10 | 1.16 | 0.25 | [-0.08 0.30] |
| leftSupFrontal | Intercept | 0.28 | 0.06 | 4.73 | < 0.001 | [0.16 0.40] |
| leftSupFrontal | Task | 0.14 | 0.04 | 3.76 | < 0.001 | [0.07 0.21] |
| leftSupFrontal | Trial Type | 0.10 | 0.04 | 2.64 | < 0.01 | [0.03 0.17] |
| leftSupFrontal | Task x Trial Type | 0.12 | 0.07 | 1.65 | 0.10 | [-0.02 0.27] |
| leftPrecentralPrecG | Intercept | 0.56 | 0.12 | 4.76 | < 0.001 | [0.32 0.80] |
| leftPrecentralPrecG | Task | -0.30 | 0.06 | -4.79 | < 0.001 | [-0.43 -0.18] |
| leftPrecentralPrecG | Trial Type | 0.15 | 0.06 | 2.32 | < 0.05 | [0.02 0.27] |
| leftPrecentralPrecG | Task x Trial Type | 0.07 | 0.13 | 0.53 | 0.59 | [-0.18 0.32] |
| leftPrecentralIFGorb | Intercept | 0.55 | 0.11 | 4.97 | < 0.001 | [0.33 0.77] |
| leftPrecentralIFGorb | Task | -0.51 | 0.07 | -6.86 | < 0.001 | [-0.66 -0.36] |
| leftPrecentralIFGorb | Trial Type | 0.15 | 0.07 | 2.00 | < 0.05 | [0.002 0.30] |
| leftPrecentralIFGorb | Task x Trial Type | -0.04 | 0.15 | -0.29 | 0.78 | [-0.34 0.25] |
| leftMidFrontal | Intercept | 0.33 | 0.07 | 4.49 | < 0.001 | [0.18 0.48] |
| leftMidFrontal | Task | -0.03 | 0.05 | -0.63 | 0.53 | [-0.14 0.07] |
| leftMidFrontal | Trial Type | 0.16 | 0.05 | 3.11 | < 0.01 | [0.06 0.27] |
| leftMidFrontal | Task x Trial Type | 0.002 | 0.11 | 0.01 | 0.99 | [-0.21 0.21] |
| leftMidFrontalOrb | Intercept | 0.22 | 0.08 | 2.79 | < 0.01 | [0.06 0.38] |
| leftMidFrontalOrb | Task | -0.32 | 0.06 | -5.13 | < 0.001 | [-0.44 -0.20] |
| leftMidFrontalOrb | Trial Type | 0.07 | 0.06 | 1.16 | 0.25 | [-0.05 0.20] |

| ROI | term | estimate | SE | t-value | p-value | CI |
|---|---|---|---|---|---|---|
| leftMidFrontalOrb | Task x Trial Type | 0.01 | 0.12 | 0.09 | 0.93 | [-0.24 0.26] |
| leftInsula | Intercept | 0.53 | 0.11 | 4.98 | < 0.001 | [0.32 0.75] |
| leftInsula | Task | -0.86 | 0.07 | -12.70 | < 0.001 | [-0.99 -0.72] |
| leftInsula | Trial Type | 0.07 | 0.07 | 1.09 | 0.28 | [-0.06 0.21] |
| leftInsula | Task x Trial Type | -0.01 | 0.14 | -0.08 | 0.94 | [-0.28 0.26] |
| leftMedialFrontal | Intercept | 0.64 | 0.11 | 5.76 | < 0.001 | [0.42 0.87] |
| leftMedialFrontal | Task | -0.87 | 0.07 | -12.63 | < 0.001 | [-1.00 -0.73] |
| leftMedialFrontal | Trial Type | 0.10 | 0.07 | 1.46 | 0.15 | [-0.04 0.24] |
| leftMedialFrontal | Task x Trial Type | -0.02 | 0.14 | -0.15 | 0.88 | [-0.29 0.25] |
| rightPostParietal | Intercept | 0.65 | 0.10 | 6.69 | < 0.001 | [0.46 0.85] |
| rightPostParietal | Task | -0.45 | 0.06 | -7.30 | < 0.001 | [-0.57 -0.32] |
| rightPostParietal | Trial Type | 0.11 | 0.06 | 1.74 | 0.08 | [-0.01 0.23] |
| rightPostParietal | Task x Trial Type | 0.06 | 0.12 | 0.47 | 0.64 | [-0.18 0.30] |
| rightMidParietal | Intercept | 0.30 | 0.05 | 5.81 | < 0.001 | [0.20 0.41] |
| rightMidParietal | Task | -0.04 | 0.04 | -1.02 | 0.31 | [-0.13 0.04] |
| rightMidParietal | Trial Type | 0.05 | 0.04 | 1.08 | 0.28 | [-0.04 0.13] |
| rightMidParietal | Task x Trial Type | 0.08 | 0.09 | 0.86 | 0.39 | [-0.10 0.25] |
| rightAntParietal | Intercept | 0.23 | 0.06 | 3.88 | < 0.001 | [0.11 0.35] |
| rightAntParietal | Task | -0.07 | 0.04 | -1.62 | 0.11 | [-0.15 0.01] |
| rightAntParietal | Trial Type | 0.06 | 0.04 | 1.33 | 0.19 | [-0.03 0.14] |
| rightAntParietal | Task x Trial Type | 0.05 | 0.09 | 0.62 | 0.54 | [-0.12 0.22] |
| rightSupFrontal | Intercept | 0.22 | 0.06 | 3.92 | < 0.001 | [0.11 0.33] |
| rightSupFrontal | Task | -0.07 | 0.04 | -1.75 | 0.08 | [-0.15 0.01] |
| rightSupFrontal | Trial Type | 0.03 | 0.04 | 0.77 | 0.44 | [-0.05 0.11] |
| rightSupFrontal | Task x Trial Type | 0.07 | 0.08 | 0.84 | 0.40 | [-0.09 0.23] |
| rightPrecentralPrecG | Intercept | 0.41 | 0.06 | 7.24 | < 0.001 | [0.30 0.52] |
| rightPrecentralPrecG | Task | -0.46 | 0.05 | -9.70 | < 0.001 | [-0.55 -0.36] |
| rightPrecentralPrecG | Trial Type | 0.01 | 0.05 | 0.29 | 0.77 | [-0.08 0.11] |
| rightPrecentralPrecG | Task x Trial Type | 0.01 | 0.09 | 0.09 | 0.93 | [-0.18 0.19] |
| rightPrecentralIFGorb | Intercept | 0.54 | 0.09 | 6.31 | < 0.001 | [0.37 0.71] |
| rightPrecentralIFGorb | Task | -0.61 | 0.06 | -9.69 | < 0.001 | [-0.73 -0.48] |
| rightPrecentralIFGorb | Trial Type | 0.06 | 0.06 | 0.87 | 0.38 | [-0.07 0.18] |
| rightPrecentralIFGorb | Task x Trial Type | -0.03 | 0.13 | -0.25 | 0.80 | [-0.28 0.22] |

Table of Contents

| ROI | term | estimate | SE | t-value | p-value | CI |
|---|---|---|---|---|---|---|
| rightMidFrontal | Intercept | 0.15 | 0.04 | 3.64 | < 0.001 | [0.07 0.23] |
| rightMidFrontal | Task | -0.20 | 0.04 | -5.27 | < 0.001 | [-0.27 -0.12] |
| rightMidFrontal | Trial Type | 0.03 | 0.04 | 0.81 | 0.42 | [-0.04 0.10] |
| rightMidFrontal | Task x Trial Type | 0.01 | 0.07 | 0.17 | 0.86 | [-0.13 0.16] |
| rightMidFrontalOrb | Intercept | 0.16 | 0.06 | 2.68 | < 0.01 | [0.04 0.29] |
| rightMidFrontalOrb | Task | -0.34 | 0.05 | -7.34 | < 0.001 | [-0.43 -0.25] |
| rightMidFrontalOrb | Trial Type | 0.01 | 0.05 | 0.32 | 0.75 | [-0.08 0.10] |
| rightMidFrontalOrb | Task x Trial Type | -0.04 | 0.09 | -0.39 | 0.69 | [-0.22 0.14] |
| rightInsula | Intercept | 0.38 | 0.08 | 4.85 | < 0.001 | [0.22 0.54] |
| rightInsula | Task | -0.66 | 0.05 | -12.86 | < 0.001 | [-0.77 -0.56] |
| rightInsula | Trial Type | 0.04 | 0.05 | 0.87 | 0.38 | [-0.06 0.15] |
| rightInsula | Task x Trial Type | -0.10 | 0.10 | -0.97 | 0.34 | [-0.30 0.10] |
| rightMedialFrontal | Intercept | 0.70 | 0.12 | 6.12 | < 0.001 | [0.47 0.94] |
| rightMedialFrontal | Task | -1.17 | 0.08 | -14.77 | < 0.001 | [-1.33 -1.01] |
| rightMedialFrontal | Trial Type | 0.06 | 0.08 | 0.75 | 0.46 | [-0.10 0.22] |
| rightMedialFrontal | Task x Trial Type | -0.01 | 0.16 | -0.08 | 0.94 | [-0.32 0.30] |

**Table S16. By-ROI Response Estimates within MD Parcels Using the Task-Switching Localizer.** The table presents the by-ROI results of a mixed-effects model examining the effects of Task, Trial Type, and their interaction on response estimates, using the task-switching localizer.

| ROI | term | estimate | SE | t-value | p-value | CI |
|---|---|---|---|---|---|---|
| leftPostParietal | Intercept | 1.00 | 0.12 | 8.16 | < 0.001 | [0.76 1.25] |
| leftPostParietal | Task | -0.08 | 0.07 | -1.22 | 0.22 | [-0.22 0.05] |
| leftPostParietal | Trial Type | 0.28 | 0.07 | 4.12 | < 0.001 | [0.14 0.41] |
| leftPostParietal | Task x Trial Type | 0.23 | 0.14 | 1.72 | 0.09 | [-0.03 0.50] |
| leftMidParietal | Intercept | 0.86 | 0.12 | 6.85 | < 0.001 | [0.60 1.11] |
| leftMidParietal | Task | -0.07 | 0.07 | -1.01 | 0.31 | [-0.22 0.07] |
| leftMidParietal | Trial Type | 0.26 | 0.07 | 3.64 | < 0.001 | [0.12 0.41] |

| ROI | term | estimate | SE | t-value | p-value | CI |
|---|---|---|---|---|---|---|
| leftMidParietal | Task x Trial Type | 0.24 | 0.14 | 1.66 | 0.10 | [-0.05 0.53] |
| leftAntParietal | Intercept | 0.65 | 0.10 | 6.62 | < 0.001 | [0.45 0.85] |
| leftAntParietal | Task | 0.17 | 0.06 | 2.97 | < 0.01 | [0.06 0.28] |
| leftAntParietal | Trial Type | 0.20 | 0.06 | 3.54 | < 0.001 | [0.09 0.31] |
| leftAntParietal | Task x Trial Type | 0.14 | 0.11 | 1.28 | 0.20 | [-0.08 0.37] |
| leftSupFrontal | Intercept | 0.40 | 0.07 | 5.96 | < 0.001 | [0.26 0.53] |
| leftSupFrontal | Task | 0.15 | 0.05 | 3.21 | < 0.01 | [0.06 0.24] |
| leftSupFrontal | Trial Type | 0.14 | 0.05 | 2.91 | < 0.01 | [0.04 0.23] |
| leftSupFrontal | Task x Trial Type | 0.15 | 0.09 | 1.56 | 0.12 | [-0.04 0.33] |
| leftPrecentralPrecG | Intercept | 0.83 | 0.13 | 6.28 | < 0.001 | [0.56 1.10] |
| leftPrecentralPrecG | Task | -0.42 | 0.07 | -5.83 | < 0.001 | [-0.56 -0.28] |
| leftPrecentralPrecG | Trial Type | 0.22 | 0.07 | 3.05 | < 0.01 | [0.08 0.36] |
| leftPrecentralPrecG | Task x Trial Type | 0.07 | 0.14 | 0.52 | 0.60 | [-0.21 0.36] |
| leftPrecentralIFGorb | Intercept | 0.78 | 0.12 | 6.24 | < 0.001 | [0.52 1.03] |
| leftPrecentralIFGorb | Task | -0.53 | 0.08 | -6.52 | < 0.001 | [-0.69 -0.37] |
| leftPrecentralIFGorb | Trial Type | 0.22 | 0.08 | 2.69 | < 0.01 | [0.06 0.38] |
| leftPrecentralIFGorb | Task x Trial Type | 0.13 | 0.16 | 0.81 | 0.42 | [-0.19 0.45] |
| leftMidFrontal | Intercept | 0.66 | 0.11 | 5.78 | < 0.001 | [0.43 0.89] |
| leftMidFrontal | Task | -0.09 | 0.07 | -1.22 | 0.22 | [-0.22 0.05] |
| leftMidFrontal | Trial Type | 0.22 | 0.07 | 3.20 | < 0.01 | [0.09 0.36] |
| leftMidFrontal | Task x Trial Type | 0.18 | 0.14 | 1.29 | 0.20 | [-0.10 0.45] |
| leftMidFrontalOrb | Intercept | 0.29 | 0.08 | 3.58 | < 0.001 | [0.12 0.45] |
| leftMidFrontalOrb | Task | -0.25 | 0.06 | -4.10 | < 0.001 | [-0.38 -0.13] |

| ROI | term | estimate | SE | t-value | p-value | CI |
|---|---|---|---|---|---|---|
| leftMidFrontalOrb | Trial Type | 0.11 | 0.06 | 1.75 | 0.08 | [-0.01 0.23] |
| leftMidFrontalOrb | Task x Trial Type | 0.07 | 0.12 | 0.59 | 0.56 | [-0.17 0.32] |
| leftInsula | Intercept | 0.60 | 0.13 | 4.74 | < 0.001 | [0.34 0.85] |
| leftInsula | Task | -0.89 | 0.08 | -10.61 | < 0.001 | [-1.06 -0.73] |
| leftInsula | Trial Type | 0.07 | 0.08 | 0.88 | 0.38 | [-0.09 0.24] |
| leftInsula | Task x Trial Type | -0.002 | 0.17 | -0.01 | 0.99 | [-0.33 0.33] |
| leftMedialFrontal | Intercept | 0.64 | 0.10 | 6.66 | < 0.001 | [0.44 0.83] |
| leftMedialFrontal | Task | -0.63 | 0.06 | -10.51 | < 0.001 | [-0.75 -0.51] |
| leftMedialFrontal | Trial Type | 0.11 | 0.06 | 1.89 | 0.06 | [-0.005 0.23] |
| leftMedialFrontal | Task x Trial Type | 0.05 | 0.12 | 0.42 | 0.67 | [-0.18 0.29] |
| rightPostParietal | Intercept | 0.77 | 0.11 | 6.94 | < 0.001 | [0.54 0.99] |
| rightPostParietal | Task | -0.22 | 0.06 | -4.01 | < 0.001 | [-0.34 -0.11] |
| rightPostParietal | Trial Type | 0.14 | 0.06 | 2.49 | < 0.05 | [0.03 0.25] |
| rightPostParietal | Task x Trial Type | 0.18 | 0.11 | 1.59 | 0.11 | [-0.04 0.40] |
| rightMidParietal | Intercept | 0.55 | 0.09 | 6.35 | < 0.001 | [0.37 0.72] |
| rightMidParietal | Task | -0.19 | 0.06 | -3.43 | < 0.001 | [-0.30 -0.08] |
| rightMidParietal | Trial Type | 0.11 | 0.06 | 1.94 | 0.05 | [-0.002 0.22] |
| rightMidParietal | Task x Trial Type | 0.14 | 0.11 | 1.23 | 0.22 | [-0.08 0.36] |
| rightAntParietal | Intercept | 0.30 | 0.07 | 4.10 | < 0.001 | [0.15 0.45] |
| rightAntParietal | Task | 0.07 | 0.05 | 1.57 | 0.12 | [-0.02 0.16] |
| rightAntParietal | Trial Type | 0.10 | 0.05 | 2.11 | < 0.05 | [0.01 0.19] |
| rightAntParietal | Task x Trial Type | 0.10 | 0.09 | 1.04 | 0.30 | [-0.09 0.28] |
| rightSupFrontal | Intercept | 0.20 | 0.06 | 3.64 | < 0.001 | [0.09 0.32] |

| ROI | term | estimate | SE | t-value | p-value | CI |
|---|---|---|---|---|---|---|
| rightSupFrontal | Task | 0.11 | 0.04 | 3.05 | < 0.01 | [0.04 0.18] |
| rightSupFrontal | Trial Type | 0.07 | 0.04 | 1.92 | 0.06 | [-0.002 0.14] |
| rightSupFrontal | Task x Trial Type | 0.08 | 0.07 | 1.09 | 0.28 | [-0.06 0.22] |
| rightPrecentralPrecG | Intercept | 0.50 | 0.08 | 6.36 | < 0.001 | [0.34 0.65] |
| rightPrecentralPrecG | Task | -0.30 | 0.04 | -6.50 | < 0.001 | [-0.38 -0.20] |
| rightPrecentralPrecG | Trial Type | 0.06 | 0.04 | 1.34 | 0.18 | [-0.03 0.15] |
| rightPrecentralPrecG | Task x Trial Type | 0.09 | 0.09 | 0.96 | 0.34 | [-0.09 0.27] |
| rightPrecentralIFGorb | Intercept | 0.57 | 0.08 | 7.22 | < 0.001 | [0.41 0.73] |
| rightPrecentralIFGorb | Task | -0.41 | 0.05 | -7.80 | < 0.001 | [-0.51 -0.31] |
| rightPrecentralIFGorb | Trial Type | 0.09 | 0.05 | 1.65 | 0.10 | [-0.02 0.19] |
| rightPrecentralIFGorb | Task x Trial Type | 0.03 | 0.10 | 0.25 | 0.80 | [-0.18 0.23] |
| rightMidFrontal | Intercept | 0.38 | 0.07 | 5.40 | < 0.001 | [0.24 0.52] |
| rightMidFrontal | Task | -0.23 | 0.04 | -5.31 | < 0.001 | [-0.32 -0.15] |
| rightMidFrontal | Trial Type | 0.05 | 0.04 | 1.21 | 0.23 | [-0.03 0.14] |
| rightMidFrontal | Task x Trial Type | 0.12 | 0.09 | 1.31 | 0.19 | [-0.06 0.29] |
| rightMidFrontalOrb | Intercept | 0.20 | 0.06 | 3.59 | < 0.001 | [0.09 0.32] |
| rightMidFrontalOrb | Task | -0.30 | 0.04 | -6.90 | < 0.001 | [-0.38 -0.21] |
| rightMidFrontalOrb | Trial Type | 0.03 | 0.04 | 0.72 | 0.48 | [-0.05 0.12] |
| rightMidFrontalOrb | Task x Trial Type | 0.03 | 0.09 | 0.35 | 0.72 | [-0.14 0.20] |
| rightInsula | Intercept | 0.38 | 0.07 | 5.10 | < 0.001 | [0.23 0.53] |
| rightInsula | Task | -0.61 | 0.06 | -10.69 | < 0.001 | [-0.73 -0.50] |
| rightInsula | Trial Type | 0.03 | 0.06 | 0.52 | 0.60 | [-0.08 0.14] |
| rightInsula | Task x Trial Type | 0.05 | 0.12 | 0.41 | 0.68 | [-0.18 0.27] |

| ROI | term | estimate | SE | t-value | p-value | CI |
|---|---|---|---|---|---|---|
| rightMedialFrontal | Intercept | 0.68 | 0.12 | 5.83 | < 0.001 | [0.44 0.91] |
| rightMedialFrontal | Task | -0.86 | 0.08 | -11.34 | < 0.001 | [-1.01 -0.71] |
| rightMedialFrontal | Trial Type | 0.08 | 0.08 | 1.05 | 0.30 | [-0.07 0.23] |
| rightMedialFrontal | Task x Trial Type | 0.01 | 0.15 | 0.06 | 0.96 | [-0.29 0.31] |

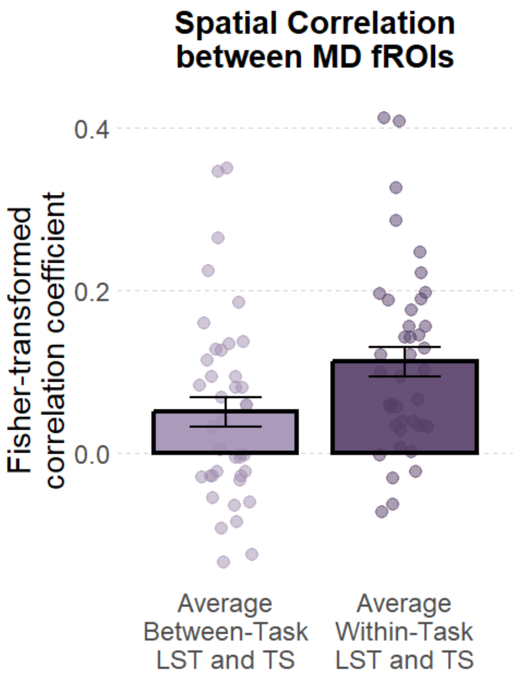


**Figure S14. Spatial Correlation between MD fROIs.** The figure shows a bar plot of the fisher-transformed correlation coefficient between MD fROIs. For the between-task condition, we computed the correlation coefficients across different tasks (e.g., LST-run1 with TS-run1) and averaged the values. For the within-task condition, we computed the correlation coefficients between runs of the same task (e.g., LST-run1 with LST-run2) and averaged the two values. LST: language switching task, TS: nonverbal task switching task.

**Table S17. Spatial Correlation between MD fROIs.** The table presents the results of a mixed-effects model examining the effects of Task Type on spatial correlation between the MD fROIs.

| term | estimate | SE | t-value | p-value | CI |
|---|---|---|---|---|---|
| Intercept | 0.08 | 0.02 | 3.74 | < 0.001 | [0.04 0.13] |
| Task Type | -0.06 | 0.01 | -5.76 | < 0.001 | [-0.08 -0.04] |

**Figure S15. By-ROI Spatial Correlation between MD fROIs.** The figure shows bar plots of the fisher-transformed correlation coefficient in MD fROIs. For the between-task condition, we computed the correlation coefficients across different tasks (e.g., LST-run1 with TS-run1) and averaged the values. For the within-task condition, we computed the correlation coefficients between runs of the same task (e.g., LST-run1 with LST-run2) and averaged the two values. LST: language switching task, TS: nonverbal task switching task.

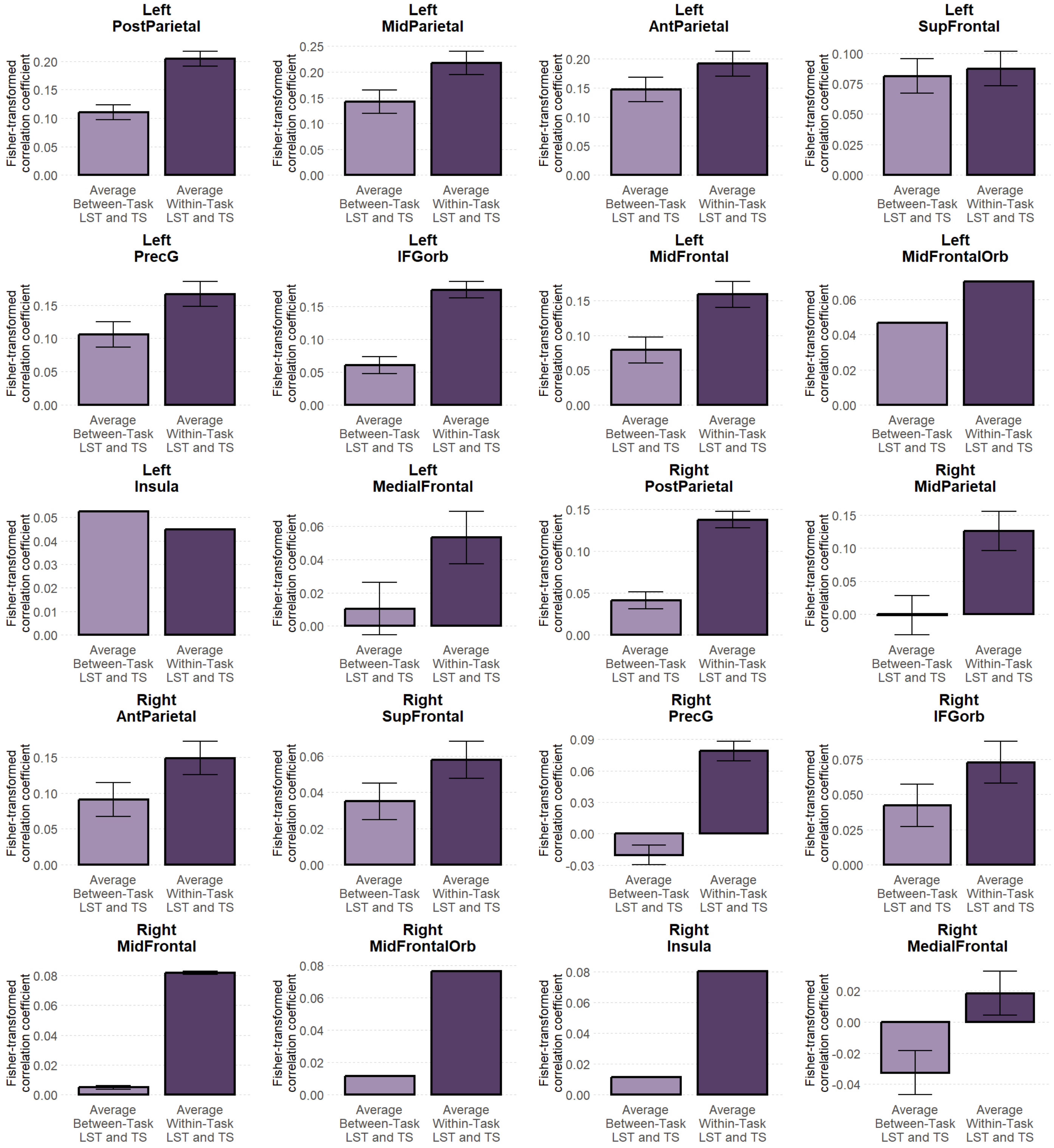


**Table S18. By-ROI Spatial Correlation in each MD fROI.** The table presents the by-ROI results of a mixed-effects model examining the effects of Task Type on spatial correlation in each MD fROI.

Table of Contents

| ROI | term | estimate | SE | t-value | p-value | CI |
|---|---|---|---|---|---|---|
| leftPostParietal | Intercept | 0.16 | 0.04 | 4.53 | < 0.001 | [0.09 0.23] |
| leftPostParietal | Task Type | -0.09 | 0.06 | -1.72 | 0.09 | [-0.20 0.02] |
| leftMidParietal | Intercept | 0.18 | 0.04 | 4.53 | < 0.001 | [0.10 0.26] |
| leftMidParietal | Task Type | -0.07 | 0.05 | -1.44 | 0.16 | [-0.18 0.03] |
| leftAntParietal | Intercept | 0.17 | 0.03 | 5.36 | < 0.001 | [0.11 0.23] |
| leftAntParietal | Task Type | -0.04 | 0.04 | -1.23 | 0.22 | [-0.12 0.03] |
| leftSupFrontal | Intercept | 0.08 | 0.03 | 2.94 | < 0.01 | [0.03 0.14] |
| leftSupFrontal | Task Type | -0.01 | 0.04 | -0.15 | 0.88 | [-0.09 0.08] |
| leftPrecentralPrecG | Intercept | 0.14 | 0.04 | 3.25 | < 0.01 | [0.05 0.22] |
| leftPrecentralPrecG | Task Type | -0.06 | 0.06 | -0.98 | 0.34 | [-0.19 0.06] |
| leftPrecentralIFGorb | Intercept | 0.12 | 0.04 | 3.20 | < 0.01 | [0.04 0.19] |
| leftPrecentralIFGorb | Task Type | -0.12 | 0.06 | -1.92 | 0.06 | [-0.24 0.01] |
| leftMidFrontal | Intercept | 0.12 | 0.03 | 3.69 | < 0.001 | [0.05 0.18] |
| leftMidFrontal | Task Type | -0.08 | 0.04 | -1.90 | 0.06 | [-0.16 0.005] |
| leftMidFrontalOrb | Intercept | 0.06 | 0.03 | 2.31 | < 0.05 | [0.01 0.11] |
| leftMidFrontalOrb | Task Type | -0.02 | 0.05 | -0.46 | 0.65 | [-0.12 0.08] |
| leftInsula | Intercept | 0.05 | 0.02 | 2.09 | < 0.05 | [0.002 0.10] |
| leftInsula | Task Type | 0.01 | 0.05 | 0.17 | 0.87 | [-0.09 0.10] |
| leftMedialFrontal | Intercept | 0.03 | 0.03 | 1.07 | 0.29 | [-0.03 0.09] |
| leftMedialFrontal | Task Type | -0.04 | 0.04 | -1.05 | 0.30 | [-0.12 0.04] |
| rightPostParietal | Intercept | 0.09 | 0.03 | 3.07 | < 0.01 | [0.03 0.15] |
| rightPostParietal | Task Type | -0.10 | 0.05 | -2.04 | < 0.05 | [-0.19 -0.0007] |
| rightMidParietal | Intercept | 0.06 | 0.04 | 1.48 | 0.15 | [-0.02 0.15] |

| ROI | term | estimate | SE | t-value | p-value | CI |
|---|---|---|---|---|---|---|
| rightMidParietal | Task Type | -0.13 | 0.04 | -2.82 | < 0.01 | [-0.22 -0.04] |
| rightAntParietal | Intercept | 0.12 | 0.03 | 3.56 | < 0.001 | [0.05 0.19] |
| rightAntParietal | Task Type | -0.06 | 0.04 | -1.55 | 0.13 | [-0.13 0.02] |
| rightSupFrontal | Intercept | 0.05 | 0.03 | 1.82 | 0.08 | [-0.005 0.10] |
| rightSupFrontal | Task Type | -0.02 | 0.04 | -0.58 | 0.57 | [-0.10 0.06] |
| rightPrecentralPrecG | Intercept | 0.03 | 0.03 | 1.08 | 0.28 | [-0.03 0.08] |
| rightPrecentralPrecG | Task Type | -0.10 | 0.04 | -2.28 | < 0.05 | [-0.19 -0.01] |
| rightPrecentralIFGorb | Intercept | 0.06 | 0.03 | 1.80 | 0.08 | [-0.01 0.12] |
| rightPrecentralIFGorb | Task Type | -0.03 | 0.05 | -0.66 | 0.52 | [-0.12 0.06] |
| rightMidFrontal | Intercept | 0.04 | 0.02 | 2.27 | < 0.05 | [0.005 0.08] |
| rightMidFrontal | Task Type | -0.08 | 0.04 | -2.07 | < 0.05 | [-0.15 -0.002] |
| rightMidFrontalOrb | Intercept | 0.04 | 0.02 | 2.21 | < 0.05 | [0.004 0.08] |
| rightMidFrontalOrb | Task Type | -0.06 | 0.04 | -1.64 | 0.10 | [-0.14 0.01] |
| rightInsula | Intercept | 0.05 | 0.02 | 2.31 | < 0.05 | [0.01 0.09] |
| rightInsula | Task Type | -0.07 | 0.04 | -1.73 | 0.09 | [-0.15 0.01] |
| rightMedialFrontal | Intercept | -0.01 | 0.03 | -0.22 | 0.83 | [-0.07 0.06] |
| rightMedialFrontal | Task Type | -0.05 | 0.05 | -1.05 | 0.30 | [-0.15 0.05] |

### Spatial Overlap Collapsed over the MD Parcels

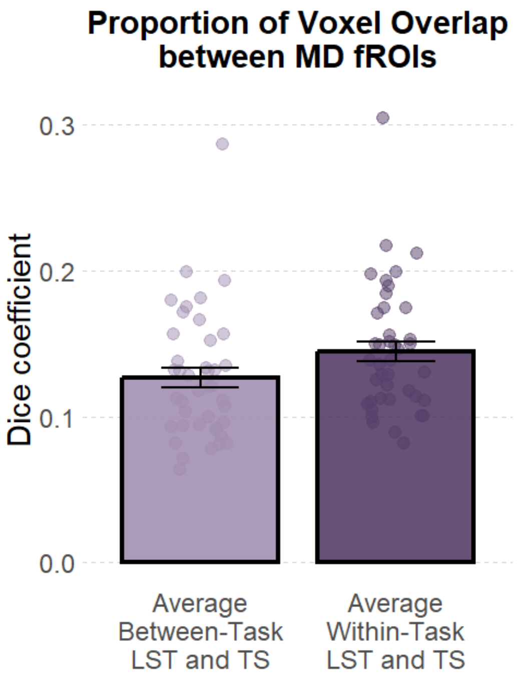


**Figure S16. Proportion of Spatial Overlap between MD fROIs.** The figure shows a bar plot of the proportion of spatial overlap between MD fROIs. For the between-task condition, we computed the dice coefficients across different tasks (e.g., LST-run1 with TS-run1) and averaged the values. For the within-task condition, we computed the dice coefficients between runs of the same task (e.g., LST-run1 with LST-run2) and averaged the two values. LST: language switching task, TS: nonverbal task switching task.

**Table S19. Proportion of Spatial Overlap between MD fROIs.** The table presents the results of a mixed-effects model examining the effects of Task Type on spatial overlap between the MD fROIs.

| term | estimate | SE | t-value | p-value | CI |
|---|---|---|---|---|---|
| Intercept | 0.14 | 0.01 | 16.10 | < 0.001 | [0.12 0.15] |
| Task Type | -0.02 | 0.005 | -3.92 | < 0.001 | [-0.03 -0.01] |

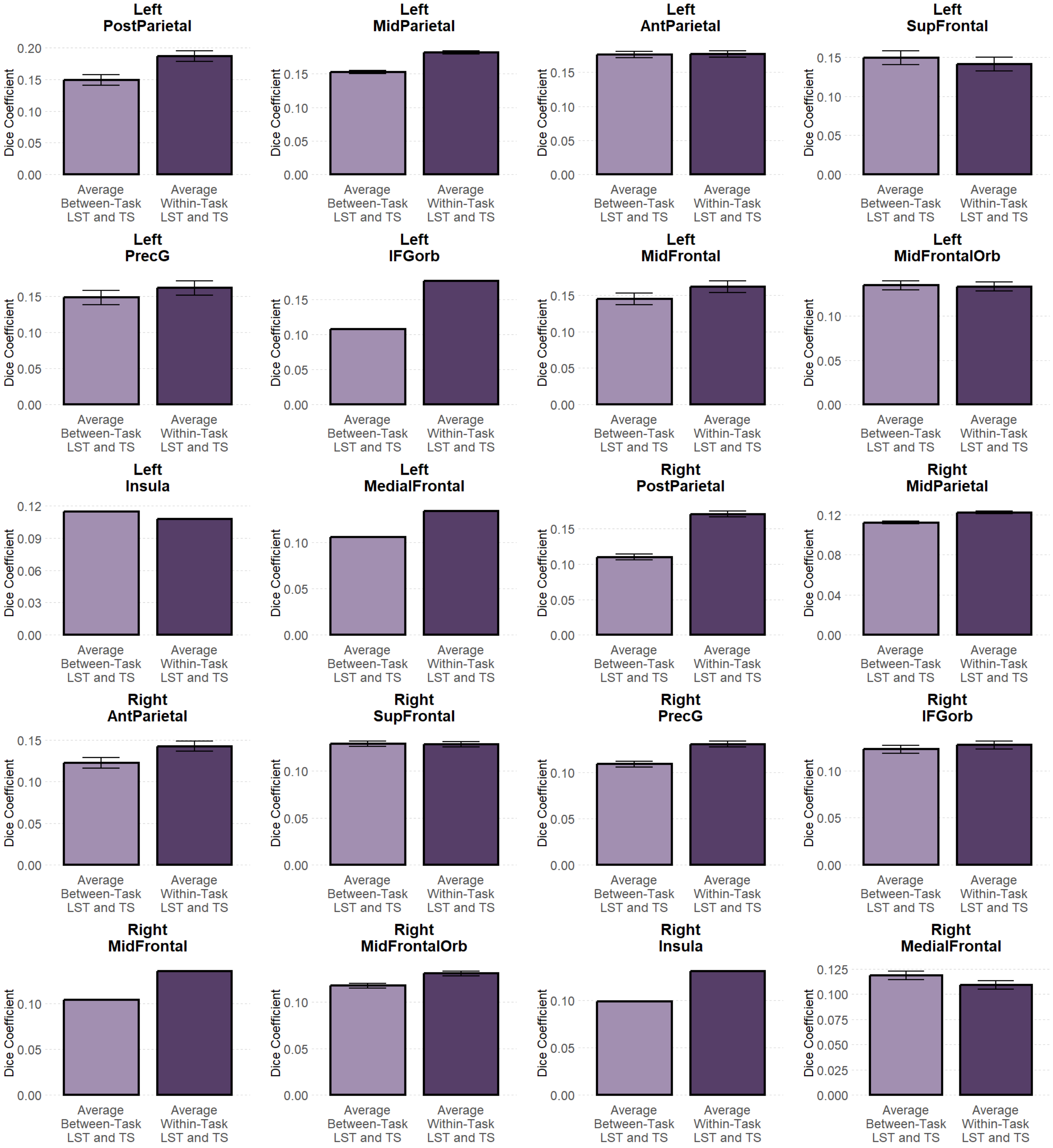


**Figure S17. By-ROI Proportion of Spatial Overlap in MD fROIs.** The figure shows bar plots of the dice coefficient in MD fROIs. For the between-task condition, we computed the dice coefficients across different tasks (e.g., LST-run1 with TS-run1) and averaged the values. For the within-task condition, we computed the dice coefficients between runs of the same task (e.g., LST-run1 with LST-run2) and averaged the two values. LST: language switching task, TS: nonverbal task switching task.

**Table S20. By-ROI Proportion of Spatial Overlap in MD fROIs.** The table presents the by-ROI results of a mixed-effects model examining the effects of Task Type on proportion of spatial overlap in each MD fROI.

| ROI | term | estimate | SE | t-value | p-value | CI |
|---|---|---|---|---|---|---|
| leftPostParietal | Intercept | 0.17 | 0.01 | 12.11 | < 0.001 | [0.14 0.20] |
| leftPostParietal | Task Type | -0.04 | 0.02 | -2.18 | < 0.05 | [-0.07 -0.003] |
| leftMidParietal | Intercept | 0.17 | 0.02 | 10.48 | < 0.001 | [0.14 0.20] |
| leftMidParietal | Task Type | -0.03 | 0.03 | -0.99 | 0.33 | [-0.09 0.03] |
| leftAntParietal | Intercept | 0.18 | 0.01 | 12.27 | < 0.001 | [0.15 0.20] |
| leftAntParietal | Task Type | -0.001 | 0.02 | -0.05 | 0.96 | [-0.05 0.05] |
| leftSupFrontal | Intercept | 0.14 | 0.01 | 10.66 | < 0.001 | [0.12 0.17] |
| leftSupFrontal | Task Type | 0.01 | 0.02 | 0.49 | 0.63 | [-0.03 0.04] |
| leftPrecentralPrecG | Intercept | 0.16 | 0.02 | 8.95 | < 0.001 | [0.12 0.19] |
| leftPrecentralPrecG | Task Type | -0.01 | 0.02 | -0.58 | 0.57 | [-0.06 0.03] |
| leftPrecentralIFGorb | Intercept | 0.14 | 0.01 | 11.31 | < 0.001 | [0.12 0.17] |
| leftPrecentralIFGorb | Task Type | -0.07 | 0.03 | -2.76 | < 0.01 | [-0.12 -0.02] |
| leftMidFrontal | Intercept | 0.15 | 0.02 | 9.23 | < 0.001 | [0.12 0.19] |
| leftMidFrontal | Task Type | -0.02 | 0.02 | -0.70 | 0.49 | [-0.06 0.03] |
| leftMidFrontalOrb | Intercept | 0.13 | 0.01 | 9.29 | < 0.001 | [0.10 0.16] |
| leftMidFrontalOrb | Task Type | 0.001 | 0.02 | 0.05 | 0.96 | [-0.05 0.05] |
| leftInsula | Intercept | 0.11 | 0.01 | 10.13 | < 0.001 | [0.09 0.13] |
| leftInsula | Task Type | 0.01 | 0.02 | 0.32 | 0.75 | [-0.04 0.05] |
| leftMedialFrontal | Intercept | 0.12 | 0.01 | 12.77 | < 0.001 | [0.10 0.14] |
| leftMedialFrontal | Task Type | -0.03 | 0.02 | -1.50 | 0.14 | [-0.06 0.01] |
| rightPostParietal | Intercept | 0.14 | 0.01 | 12.98 | < 0.001 | [0.12 0.16] |
| rightPostParietal | Task Type | -0.06 | 0.02 | -3.58 | < 0.001 | [-0.10 -0.03] |

| ROI | term | estimate | SE | t-value | p-value | CI |
|---|---|---|---|---|---|---|
| rightMidParietal | Intercept | 0.12 | 0.01 | 10.75 | < 0.001 | [0.10 0.14] |
| rightMidParietal | Task Type | -0.01 | 0.02 | -0.49 | 0.62 | [-0.05 0.03] |
| rightAntParietal | Intercept | 0.13 | 0.01 | 11.22 | < 0.001 | [0.11 0.16] |
| rightAntParietal | Task Type | -0.02 | 0.02 | -1.22 | 0.23 | [-0.05 0.01] |
| rightSupFrontal | Intercept | 0.13 | 0.01 | 12.74 | < 0.001 | [0.11 0.15] |
| rightSupFrontal | Task Type | 0.0007 | 0.02 | 0.04 | 0.96 | [-0.03 0.04] |
| rightPrecentralPrecG | Intercept | 0.12 | 0.01 | 12.10 | < 0.001 | [0.10 0.14] |
| rightPrecentralPrecG | Task Type | -0.02 | 0.02 | -1.32 | 0.19 | [-0.06 0.01] |
| rightPrecentralIFGorb | Intercept | 0.12 | 0.01 | 10.81 | < 0.001 | [0.10 0.15] |
| rightPrecentralIFGorb | Task Type | -0.004 | 0.02 | -0.24 | 0.81 | [-0.04 0.03] |
| rightMidFrontal | Intercept | 0.12 | 0.01 | 11.38 | < 0.001 | [0.10 0.14] |
| rightMidFrontal | Task Type | -0.03 | 0.02 | -1.50 | 0.14 | [-0.07 0.01] |
| rightMidFrontalOrb | Intercept | 0.12 | 0.01 | 10.77 | < 0.001 | [0.10 0.15] |
| rightMidFrontalOrb | Task Type | -0.01 | 0.02 | -0.64 | 0.52 | [-0.05 0.03] |
| rightInsula | Intercept | 0.12 | 0.01 | 11.00 | < 0.001 | [0.09 0.14] |
| rightInsula | Task Type | -0.03 | 0.02 | -1.53 | 0.13 | [-0.07 0.01] |
| rightMedialFrontal | Intercept | 0.11 | 0.01 | 10.74 | < 0.001 | [0.09 0.14] |
| rightMedialFrontal | Task Type | 0.01 | 0.02 | 0.58 | 0.56 | [-0.02 0.04] |

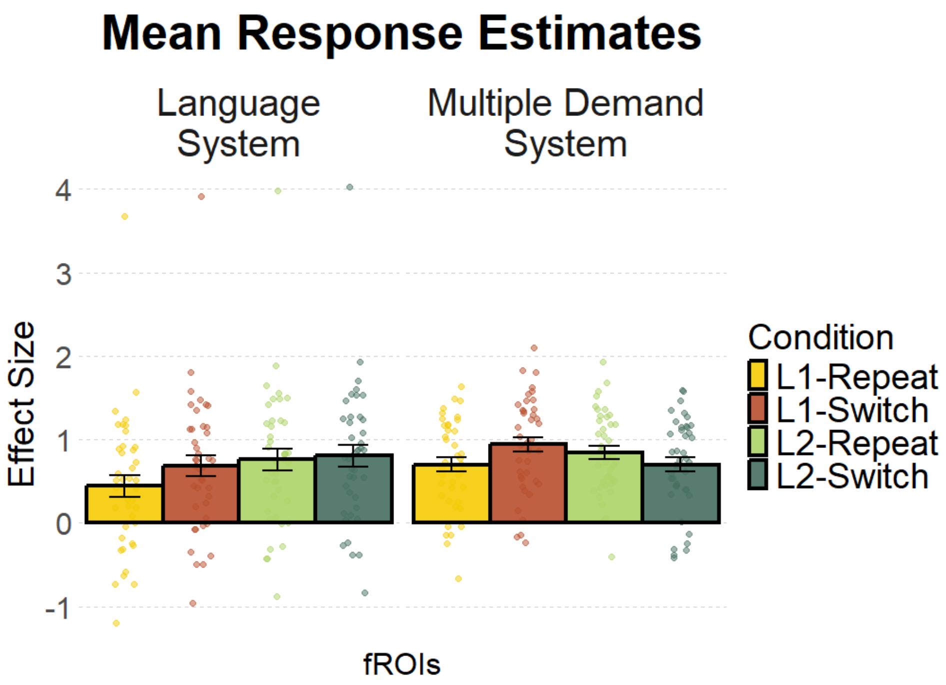


**Figure S18. Mean Response Estimates across Languages and Trial Types within the Language and MD Systems.** The figure shows bar plots of the mean response estimates, corresponding to the % BOLD signal change, across languages and trial types. The 4 bars in the left panel depict results within the language system, and the 4 bars in the right panel depict results within the MD system.

**Table S21. Mean Response Estimates across Languages and Trial Types within the Language System.** The table presents the results of a mixed-effects model examining the effects of Language, Trial Type, and their interaction on response estimates within the language system.

| term | estimate | SE | t-value | p-value | CI |
|---|---|---|---|---|---|
| Intercept | 0.67 | 0.22 | 3.00 | < 0.05 | [0.17 1.18] |
| Language | 0.22 | 0.06 | 3.45 | < 0.05 | [0.04 0.39] |
| Language Switch Type | 0.14 | 0.05 | 2.64 | < 0.01 | [0.04 0.25] |
| Language x Language Switch Type | -0.19 | 0.11 | -1.78 | 0.08 | [-0.41 0.02] |

**Table S22. Mean Response Estimates across Languages and Trial Types within the Multiple Demand System.** The table presents the results of a mixed-effects model examining the effects of Language, Trial Type, and their interaction on response estimates within the MD system.

| term | estimate | SE | t-value | p-value | CI |
|---|---|---|---|---|---|
| Intercept | 0.79 | 0.11 | 7.35 | < 0.001 | [0.58 1.01] |
| Language | -0.05 | 0.05 | -1.06 | 0.30 | [-0.14 0.04] |
| Language Switch Type | 0.05 | 0.04 | 1.13 | 0.27 | [-0.04 0.14] |
| Language x Language Switch Type | -0.38 | 0.09 | -4.25 | < 0.001 | [-0.57 -0.20] |

**Figure S19. By-ROI Mean Response Estimates across Languages and Trial Types within the Language System.** The figure shows bar plots of the by-ROI mean response estimates within the language system, corresponding to the % BOLD signal change, across languages and trial types.

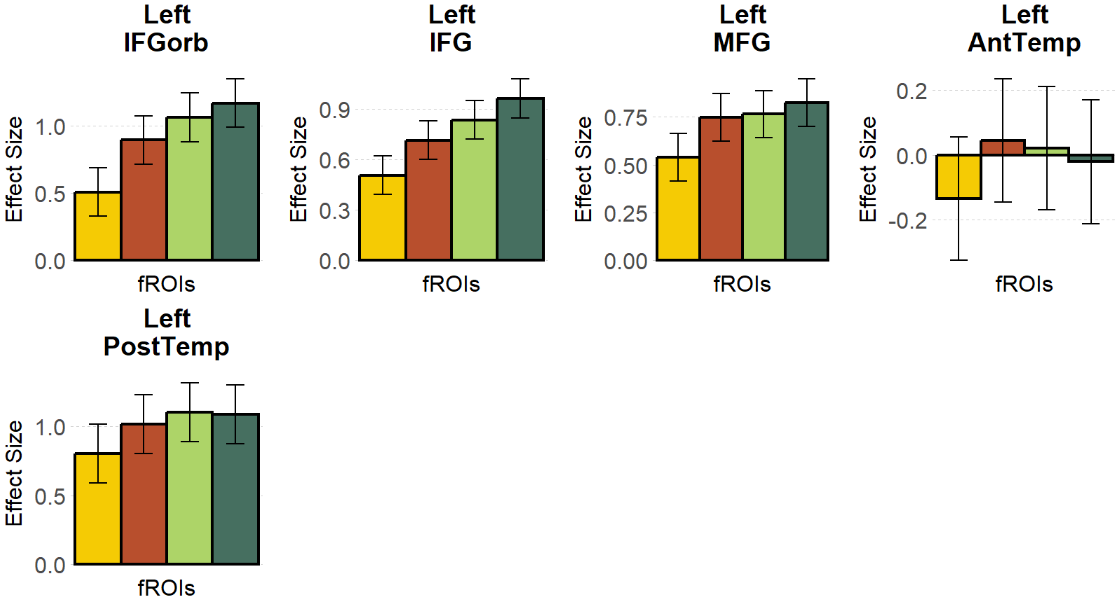


**Table S23. By-ROI Mean Response Estimates across Languages and Trial Types within the Language System.** The table presents the results of a mixed-effects model examining the effects of Language, Trial Type, and their interaction on response estimates within the language system.

| ROI | term | estimate | SE | t-value | p-value | CI |
|---|---|---|---|---|---|---|
| leftIFGorb | Intercept | 0.91 | 0.19 | 4.83 | < 0.001 | [0.53 1.29] |
| leftIFGorb | Language | 0.42 | 0.07 | 5.72 | < 0.001 | [0.27 0.56] |
| leftIFGorb | Language Switch Type | 0.25 | 0.07 | 3.39 | < 0.001 | [0.10 0.39] |
| leftIFGorb | Language x Language Switch Type | -0.28 | 0.14 | -1.95 | 0.05 | [-0.57 0.004] |
| leftIFG | Intercept | 0.75 | 0.12 | 6.18 | < 0.001 | [0.51 1.00] |

| ROI | term | estimate | SE | t-value | p-value | CI |
|---|---|---|---|---|---|---|
| leftIFG | Language | 0.29 | 0.06 | 5.23 | < 0.001 | [0.18 0.40] |
| leftIFG | Language Switch Type | 0.17 | 0.06 | 3.04 | < 0.01 | [0.06 0.28] |
| leftIFG | Language x Language Switch Type | -0.08 | 0.11 | -0.72 | 0.47 | [-0.30 0.14] |
| leftMFG | Intercept | 0.72 | 0.13 | 5.61 | < 0.001 | [0.46 0.98] |
| leftMFG | Language | 0.15 | 0.05 | 3.30 | < 0.001 | [0.06 0.24] |
| leftMFG | Language Switch Type | 0.14 | 0.05 | 2.92 | < 0.01 | [0.04 0.23] |
| leftMFG | Language x Language Switch Type | -0.15 | 0.09 | -1.61 | 0.11 | [-0.33 0.03] |
| leftAntTemp | Intercept | -0.02 | 0.19 | -0.12 | 0.91 | [-0.41 0.37] |
| leftAntTemp | Language | 0.04 | 0.04 | 1.19 | 0.23 | [-0.03 0.12] |
| leftAntTemp | Language Switch Type | 0.07 | 0.04 | 1.80 | 0.07 | [-0.01 0.14] |
| leftAntTemp | Language x Language Switch Type | -0.22 | 0.07 | -2.93 | < 0.01 | [-0.37 -0.07] |
| leftPostTemp | Intercept | 1.00 | 0.22 | 4.64 | < 0.001 | [0.57 1.44] |
| leftPostTemp | Language | 0.18 | 0.05 | 3.73 | < 0.001 | [0.09 0.28] |
| leftPostTemp | Language Switch Type | 0.10 | 0.05 | 2.02 | < 0.05 | [0.002 0.20] |
| leftPostTemp | Language x Language Switch Type | -0.23 | 0.10 | -2.34 | < 0.05 | [-0.43 -0.04] |

**Table S24. Post-hoc Comparison of Interactions for the by-ROI Mean Response Estimates across Languages and Trial Types within the Language System.** The table presents the results of post-hoc comparisons for interaction effects within the language system.

| ROI | estimate | SE | t-ratio | p-value | t-value | condition |
|---|---|---|---|---|---|---|
| leftAntTemp | -0.16 | 0.05 | -2.9157309 | < 0.01 | -2.92 | L1 - L2 within Repeat |
| leftAntTemp | 0.07 | 0.05 | 1.2278675 | 0.22 | 1.23 | L1 - L2 within Switch |
| leftPostTemp | -0.30 | 0.07 | -4.2911966 | < 0.001 | -4.29 | L1 - L2 within Repeat |
| leftPostTemp | -0.07 | 0.07 | -0.9887059 | 0.32 | -0.99 | L1 - L2 within Switch |

Table of Contents

**Figure S20. By-ROI Mean Response Estimates across Languages and Trial Types within the Multiple Demand System.** The figure shows bar plots of the by-ROI mean response estimates within the MD system, corresponding to the % BOLD signal change, across languages and trial types.

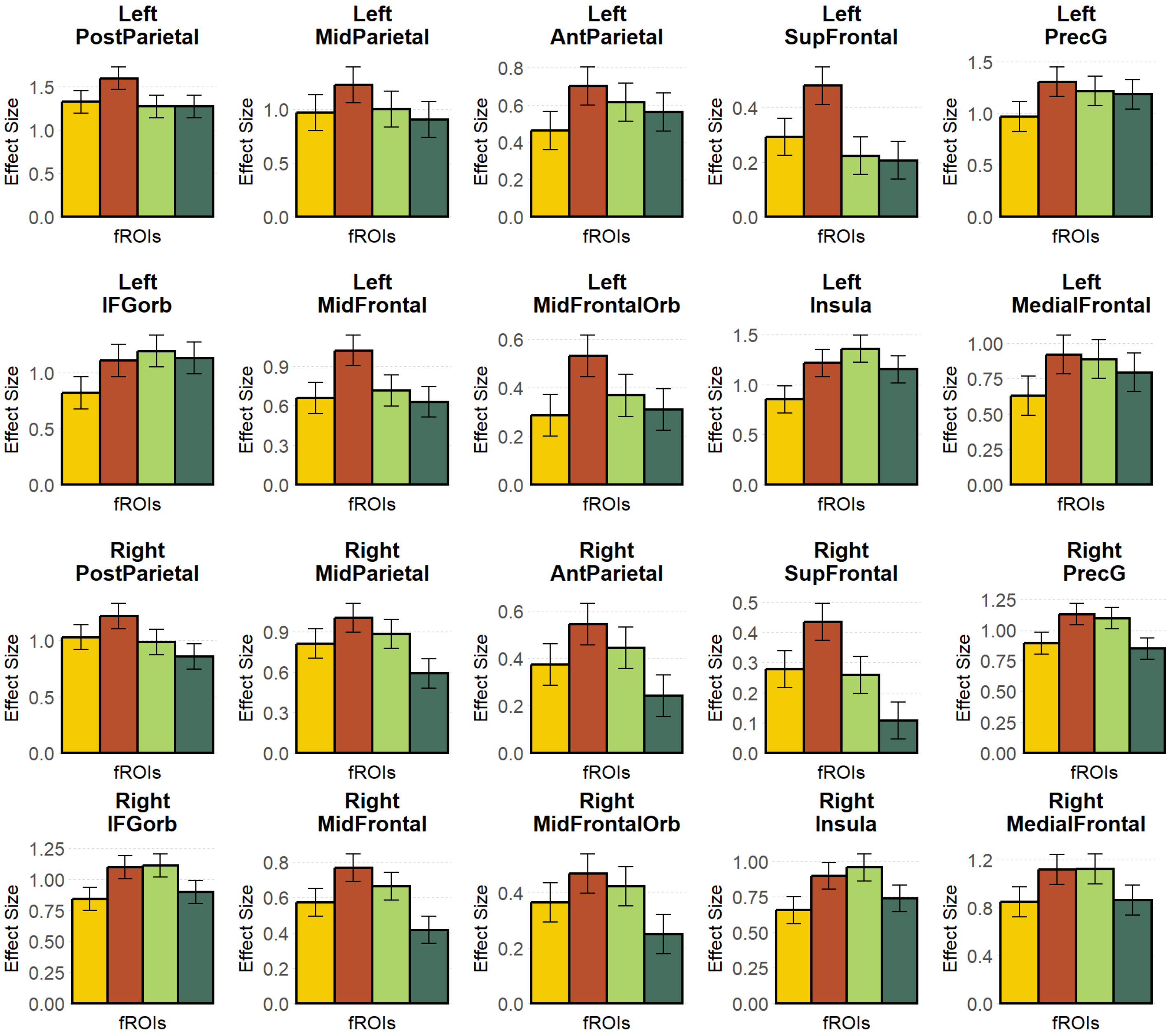


**Table S25. By-ROI Mean Response Estimates across Languages and Trial Types within the MD System.** The table presents the results of a mixed-effects model examining the effects of Language, Trial Type, and their interaction on response estimates within the MD system.

| ROI | term | estimate | SE | t-value | p-value | CI |
|---|---|---|---|---|---|---|
| leftPostParietal | Intercept | 1.36 | 0.14 | 9.98 | < 0.001 | [1.09 1.64] |

| ROI | term | estimate | SE | t-value | p-value | CI |
|---|---|---|---|---|---|---|
| leftPostParietal | Language | -0.19 | 0.06 | -3.15 | < 0.01 | [-0.31 -0.07] |
| leftPostParietal | Language Switch Type | 0.14 | 0.06 | 2.28 | < 0.05 | [0.02 0.26] |
| leftPostParietal | Language x Language Switch Type | -0.27 | 0.12 | -2.26 | < 0.05 | [-0.51 -0.03] |
| leftMidParietal | Intercept | 1.02 | 0.17 | 5.92 | < 0.001 | [0.67 1.37] |
| leftMidParietal | Language | -0.14 | 0.06 | -2.24 | < 0.05 | [-0.27 -0.02] |
| leftMidParietal | Language Switch Type | 0.08 | 0.06 | 1.27 | 0.21 | [-0.05 0.21] |
| leftMidParietal | Language x Language Switch Type | -0.36 | 0.13 | -2.76 | < 0.01 | [-0.61 -0.10] |
| leftAntParietal | Intercept | 0.58 | 0.11 | 5.40 | < 0.001 | [0.37 0.80] |
| leftAntParietal | Language | 0.005 | 0.05 | 0.10 | 0.92 | [-0.10 0.11] |
| leftAntParietal | Language Switch Type | 0.09 | 0.05 | 1.77 | 0.08 | [-0.01 0.20] |
| leftAntParietal | Language x Language Switch Type | -0.29 | 0.10 | -2.78 | < 0.01 | [-0.50 -0.08] |
| leftSupFrontal | Intercept | 0.30 | 0.08 | 3.98 | < 0.001 | [0.15 0.45] |
| leftSupFrontal | Language | -0.17 | 0.05 | -3.73 | < 0.001 | [-0.26 -0.08] |
| leftSupFrontal | Language Switch Type | 0.09 | 0.05 | 1.86 | 0.07 | [-0.01 0.18] |
| leftSupFrontal | Language x Language Switch Type | -0.20 | 0.09 | -2.22 | < 0.05 | [-0.39 -0.02] |
| leftPrecentralPrecG | Intercept | 1.17 | 0.15 | 7.72 | < 0.001 | [0.86 1.47] |
| leftPrecentralPrecG | Language | 0.06 | 0.07 | 0.88 | 0.38 | [-0.08 0.20] |
| leftPrecentralPrecG | Language Switch Type | 0.15 | 0.07 | 2.13 | < 0.05 | [0.01 0.30] |
| leftPrecentralPrecG | Language x Language Switch Type | -0.37 | 0.14 | -2.57 | < 0.05 | [-0.65 -0.09] |
| leftPrecentralIFGorb | Intercept | 1.06 | 0.15 | 7.01 | < 0.001 | [0.76 1.37] |
| leftPrecentralIFGorb | Language | 0.20 | 0.07 | 2.67 | < 0.01 | [0.05 0.34] |
| leftPrecentralIFGorb | Language Switch Type | 0.11 | 0.07 | 1.54 | 0.13 | [-0.03 0.26] |
| leftPrecentralIFGorb | Language x Language Switch Type | -0.35 | 0.15 | -2.37 | < 0.05 | [-0.64 -0.06] |
| leftMidFrontal | Intercept | 0.76 | 0.12 | 6.11 | < 0.001 | [0.51 1.01] |

| ROI | term | estimate | SE | t-value | p-value | CI |
|---|---|---|---|---|---|---|
| leftMidFrontal | Language | -0.17 | 0.06 | -2.88 | < 0.01 | [-0.28 -0.05] |
| leftMidFrontal | Language Switch Type | 0.14 | 0.06 | 2.36 | < 0.05 | [0.02 0.25] |
| leftMidFrontal | Language x Language Switch Type | -0.45 | 0.12 | -3.86 | < 0.001 | [-0.68 -0.22] |
| leftMidFrontalOrb | Intercept | 0.37 | 0.09 | 3.98 | < 0.001 | [0.18 0.56] |
| leftMidFrontalOrb | Language | -0.07 | 0.05 | -1.31 | 0.19 | [-0.17 0.04] |
| leftMidFrontalOrb | Language Switch Type | 0.09 | 0.05 | 1.76 | 0.08 | [-0.01 0.20] |
| leftMidFrontalOrb | Language x Language Switch Type | -0.30 | 0.11 | -2.87 | < 0.01 | [-0.51 -0.09] |
| leftInsula | Intercept | 1.15 | 0.14 | 8.03 | < 0.001 | [0.86 1.43] |
| leftInsula | Language | 0.22 | 0.06 | 3.70 | < 0.001 | [0.10 0.34] |
| leftInsula | Language Switch Type | 0.08 | 0.06 | 1.31 | 0.19 | [-0.04 0.20] |
| leftInsula | Language x Language Switch Type | -0.57 | 0.12 | -4.75 | < 0.001 | [-0.81 -0.33] |
| leftMedialFrontal | Intercept | 0.81 | 0.14 | 5.66 | < 0.001 | [0.52 1.10] |
| leftMedialFrontal | Language | 0.07 | 0.05 | 1.26 | 0.21 | [-0.04 0.17] |
| leftMedialFrontal | Language Switch Type | 0.10 | 0.05 | 1.88 | 0.06 | [-0.005 0.20] |
| leftMedialFrontal | Language x Language Switch Type | -0.38 | 0.10 | -3.65 | < 0.001 | [-0.59 -0.18] |
| rightPostParietal | Intercept | 1.02 | 0.12 | 8.64 | < 0.001 | [0.78 1.26] |
| rightPostParietal | Language | -0.20 | 0.05 | -3.71 | < 0.001 | [-0.31 -0.09] |
| rightPostParietal | Language Switch Type | 0.03 | 0.05 | 0.53 | 0.60 | [-0.08 0.14] |
| rightPostParietal | Language x Language Switch Type | -0.32 | 0.11 | -2.93 | < 0.01 | [-0.54 -0.10] |
| rightMidParietal | Intercept | 0.82 | 0.12 | 6.87 | < 0.001 | [0.58 1.06] |
| rightMidParietal | Language | -0.17 | 0.07 | -2.30 | < 0.05 | [-0.32 -0.02] |
| rightMidParietal | Language Switch Type | -0.05 | 0.07 | -0.70 | 0.49 | [-0.20 0.10] |
| rightMidParietal | Language x Language Switch Type | -0.48 | 0.15 | -3.26 | < 0.001 | [-0.78 -0.19] |
| rightAntParietal | Intercept | 0.40 | 0.09 | 4.26 | < 0.001 | [0.21 0.59] |

| ROI | term | estimate | SE | t-value | p-value | CI |
|---|---|---|---|---|---|---|
| rightAntParietal | Language | -0.12 | 0.05 | -2.30 | < 0.05 | [-0.22 -0.02] |
| rightAntParietal | Language Switch Type | -0.02 | 0.05 | -0.32 | 0.75 | [-0.12 0.08] |
| rightAntParietal | Language x Language Switch Type | -0.37 | 0.10 | -3.68 | < 0.001 | [-0.57 -0.17] |
| rightSupFrontal | Intercept | 0.27 | 0.07 | 3.95 | < 0.001 | [0.13 0.41] |
| rightSupFrontal | Language | -0.17 | 0.04 | -3.96 | < 0.001 | [-0.26 -0.09] |
| rightSupFrontal | Language Switch Type | 0.003 | 0.04 | 0.06 | 0.95 | [-0.08 0.09] |
| rightSupFrontal | Language x Language Switch Type | -0.31 | 0.09 | -3.55 | < 0.001 | [-0.48 -0.14] |
| rightPrecentralPrecG | Intercept | 0.99 | 0.10 | 10.34 | < 0.001 | [0.80 1.18] |
| rightPrecentralPrecG | Language | -0.04 | 0.06 | -0.70 | 0.49 | [-0.15 0.07] |
| rightPrecentralPrecG | Language Switch Type | -0.01 | 0.06 | -0.10 | 0.92 | [-0.12 0.10] |
| rightPrecentralPrecG | Language x Language Switch Type | -0.48 | 0.11 | -4.35 | < 0.001 | [-0.70 -0.26] |
| rightPrecentralIFGorb | Intercept | 0.99 | 0.10 | 9.69 | < 0.001 | [0.78 1.19] |
| rightPrecentralIFGorb | Language | 0.04 | 0.06 | 0.57 | 0.57 | [-0.09 0.16] |
| rightPrecentralIFGorb | Language Switch Type | 0.02 | 0.06 | 0.32 | 0.75 | [-0.10 0.14] |
| rightPrecentralIFGorb | Language x Language Switch Type | -0.47 | 0.12 | -3.76 | < 0.001 | [-0.72 -0.22] |
| rightMidFrontal | Intercept | 0.60 | 0.08 | 7.17 | < 0.001 | [0.44 0.78] |
| rightMidFrontal | Language | -0.13 | 0.05 | -2.75 | < 0.01 | [-0.22 -0.04] |
| rightMidFrontal | Language Switch Type | -0.03 | 0.05 | -0.55 | 0.58 | [-0.12 0.07] |
| rightMidFrontal | Language x Language Switch Type | -0.44 | 0.10 | -4.68 | < 0.001 | [-0.63 -0.26] |
| rightMidFrontalOrb | Intercept | 0.38 | 0.08 | 4.82 | < 0.001 | [0.22 0.54] |
| rightMidFrontalOrb | Language | -0.08 | 0.05 | -1.73 | 0.09 | [-0.17 0.01] |
| rightMidFrontalOrb | Language Switch Type | -0.04 | 0.05 | -0.74 | 0.46 | [-0.13 0.06] |
| rightMidFrontalOrb | Language x Language Switch Type | -0.28 | 0.09 | -2.96 | < 0.01 | [-0.46 -0.09] |
| rightInsula | Intercept | 0.81 | 0.10 | 7.99 | < 0.001 | [0.61 1.02] |

| ROI | term | estimate | SE | t-value | p-value | CI |
|---|---|---|---|---|---|---|
| rightInsula | Language | 0.07 | 0.05 | 1.33 | 0.19 | [-0.04 0.18] |
| rightInsula | Language Switch Type | 0.01 | 0.05 | 0.22 | 0.83 | [-0.10 0.12] |
| rightInsula | Language x Language Switch Type | -0.46 | 0.11 | -4.23 | < 0.001 | [-0.67 -0.24] |
| rightMedialFrontal | Intercept | 0.99 | 0.13 | 7.52 | < 0.001 | [0.72 1.26] |
| rightMedialFrontal | Language | 0.01 | 0.06 | 0.15 | 0.88 | [-0.11 0.12] |
| rightMedialFrontal | Language Switch Type | 0.004 | 0.06 | 0.06 | 0.95 | [-0.11 0.12] |
| rightMedialFrontal | Language x Language Switch Type | -0.53 | 0.12 | -4.53 | < 0.001 | [-0.76 -0.30] |

**Table S26. Post-hoc Comparison of Interactions for the by-ROI Mean Response Estimates across Languages and Trial Types within the MD System.** The table presents the results of post-hoc comparisons for interaction effects within the MD system.

| ROI | estimate | SE | t-ratio | p-value | t-value | condition |
|---|---|---|---|---|---|---|
| leftPostParietal | 0.05 | 0.08 | 0.6357291 | 0.53 | 0.64 | L1 - L2 within Repeat |
| leftPostParietal | 0.32 | 0.08 | 3.8261303 | < 0.001 | 3.83 | L1 - L2 within Switch |
| leftMidParietal | -0.03 | 0.09 | -0.3676080 | 0.71 | -0.37 | L1 - L2 within Repeat |
| leftMidParietal | 0.32 | 0.09 | 3.5348831 | < 0.001 | 3.53 | L1 - L2 within Switch |
| leftAntParietal | -0.15 | 0.07 | -2.0354977 | < 0.05 | -2.04 | L1 - L2 within Repeat |
| leftAntParietal | 0.14 | 0.07 | 1.8932361 | 0.06 | 1.89 | L1 - L2 within Switch |
| leftSupFrontal | 0.07 | 0.07 | 1.0628888 | 0.29 | 1.06 | L1 - L2 within Repeat |
| leftSupFrontal | 0.27 | 0.07 | 4.2075709 | < 0.001 | 4.21 | L1 - L2 within Switch |
| leftPrecentralPrecG | -0.25 | 0.10 | -2.4379929 | < 0.05 | -2.44 | L1 - L2 within Repeat |
| leftPrecentralPrecG | 0.12 | 0.10 | 1.1989145 | 0.23 | 1.20 | L1 - L2 within Switch |
| leftPrecentralIFGorb | -0.37 | 0.10 | -3.5638273 | < 0.001 | -3.56 | L1 - L2 within Repeat |
| leftPrecentralIFGorb | -0.02 | 0.10 | -0.2099183 | 0.83 | -0.21 | L1 - L2 within Switch |

| ROI | estimate | SE | t-ratio | p-value | t-value | condition |
|---|---|---|---|---|---|---|
| leftMidFrontal | -0.06 | 0.08 | -0.6914394 | 0.49 | -0.69 | L1 - L2 within Repeat |
| leftMidFrontal | 0.39 | 0.08 | 4.7714463 | < 0.001 | 4.77 | L1 - L2 within Switch |
| leftMidFrontalOrb | -0.08 | 0.08 | -1.1013959 | 0.27 | -1.10 | L1 - L2 within Repeat |
| leftMidFrontalOrb | 0.22 | 0.08 | 2.9523051 | < 0.01 | 2.95 | L1 - L2 within Switch |
| leftInsula | -0.51 | 0.08 | -5.9781104 | < 0.001 | -5.98 | L1 - L2 within Repeat |
| leftInsula | 0.06 | 0.08 | 0.7391305 | 0.46 | 0.74 | L1 - L2 within Switch |
| leftMedialFrontal | -0.26 | 0.07 | -3.4744885 | < 0.001 | -3.47 | L1 - L2 within Repeat |
| leftMedialFrontal | 0.13 | 0.07 | 1.6922608 | 0.09 | 1.69 | L1 - L2 within Switch |
| rightPostParietal | 0.04 | 0.08 | 0.5464977 | 0.59 | 0.55 | L1 - L2 within Repeat |
| rightPostParietal | 0.36 | 0.08 | 4.6963908 | < 0.001 | 4.70 | L1 - L2 within Switch |
| rightMidParietal | -0.07 | 0.10 | -0.6770172 | 0.50 | -0.68 | L1 - L2 within Repeat |
| rightMidParietal | 0.41 | 0.10 | 3.9356988 | < 0.001 | 3.94 | L1 - L2 within Switch |
| rightAntParietal | -0.07 | 0.07 | -0.9773530 | 0.33 | -0.98 | L1 - L2 within Repeat |
| rightAntParietal | 0.30 | 0.07 | 4.2285181 | < 0.001 | 4.23 | L1 - L2 within Switch |
| rightSupFrontal | 0.02 | 0.06 | 0.2896122 | 0.77 | 0.29 | L1 - L2 within Repeat |
| rightSupFrontal | 0.33 | 0.06 | 5.3131958 | < 0.001 | 5.31 | L1 - L2 within Switch |
| rightPrecentralPrecG | -0.20 | 0.08 | -2.5810023 | < 0.05 | -2.58 | L1 - L2 within Repeat |
| rightPrecentralPrecG | 0.28 | 0.08 | 3.5692351 | < 0.001 | 3.57 | L1 - L2 within Switch |
| rightPrecentralIFGorb | -0.27 | 0.09 | -3.0612923 | < 0.01 | -3.06 | L1 - L2 within Repeat |
| rightPrecentralIFGorb | 0.20 | 0.09 | 2.2531517 | < 0.05 | 2.25 | L1 - L2 within Switch |
| rightMidFrontal | -0.09 | 0.07 | -1.3615133 | 0.18 | -1.36 | L1 - L2 within Repeat |
| rightMidFrontal | 0.35 | 0.07 | 5.2528507 | < 0.001 | 5.25 | L1 - L2 within Switch |
| rightMidFrontalOrb | -0.06 | 0.07 | -0.8669199 | 0.39 | -0.87 | L1 - L2 within Repeat |

| ROI | estimate | SE | t-ratio | p-value | t-value | condition |
|---|---|---|---|---|---|---|
| rightMidFrontalOrb | 0.22 | 0.07 | 3.3186005 | < 0.01 | 3.32 | L1 - L2 within Switch |
| rightInsula | -0.30 | 0.08 | -3.9325768 | < 0.001 | -3.93 | L1 - L2 within Repeat |
| rightInsula | 0.16 | 0.08 | 2.0504424 | < 0.05 | 2.05 | L1 - L2 within Switch |
| rightMedialFrontal | -0.27 | 0.08 | -3.3107752 | < 0.01 | -3.31 | L1 - L2 within Repeat |
| rightMedialFrontal | 0.26 | 0.08 | 3.0988728 | < 0.01 | 3.10 | L1 - L2 within Switch |

**Table S27. Corresponding Polish and English Names of the Experimental Stimuli Used in the Language Switching Task.** The table presents the target names in Polish and English that were used as stimuli in the language-switching task for speech production.

| Picture Name in Polish | Picture Name in English |
|---|---|
| samolot | plane |
| mysz | mouse2n |
| krawat | tie |
| strugaczka | sharpener |
| wózek | pram |
| samochód | car |
| długopis | pen |
| żółw | turtle |
| dach | roof |
| kamizelka | vest1 |
| kwiatek | flower |
| chmura | cloud |
| zjeżdżalnia | slide |
| pomidor | tomato |
| motyl | butterfly |
| widelec | fork |
| szafa | wardrobe |
| ręka | hand |

| Picture Name in Polish | Picture Name in English |
|---|---|
| butelka | bottle |
| noga | leg |
| dom | house |
| lew | lion |
| szczoteczka | toothbrush |
| gruszka | pear |
| cebula | onion |
| kość | bone |
| koszyk | basket |
| mucha | fly |
| żaba | frog |
| nóż | knife |
| liść | leaf2n |
| gniazdo | nest |
| bęben | drum2n |
| okulary | sunglasses |
| koń | horse |
| zapałka | match3n |
| łyżka | spoon |
| garnek | pot2n |

| Picture Name in Polish | Picture Name in English |
|---|---|
| waga | scale2 |
| biurko | desk1 |
| małpa | monkey |
| klucz | key |
| wałek | rolling pin |
| koza | goat |
| kaczka | duck |
| smoczek | dummy |
| wiewiórka | squirrel2 |
| mrówka | ant |
